# The cognitive and neural bases of creative thought: a cross-domain meta-analysis of transcranial direct current stimulation studies

**DOI:** 10.1101/2025.03.18.644047

**Authors:** Melody M.Y. Chan, Eugene Cho, Matthew A. Lambon Ralph, Gail A. Robinson

**Author notes:** Co-first authors (Melody M.Y. Chan, Eugene Cho). Joint senior authors (Matthew A. Lambon Ralph, Gail A. Robinson). **Correspondence to**: Prof Gail A. Robinson Neuropsychology Research Clinic, School of Psychology The University of Queensland, Brisbane QLD 4072 Australia. **Open access statement** For the purpose of open access, the authors have applied a CC BY public copyright licence to any Author Accepted Manuscript version arising from this submission.

## Abstract

Creative thought enables humans to flexibly generate, evaluate and select novel and adaptive ideas according to different contexts. Decades of creativity research indicates that it involves at least two aspects: retrieval of previously acquired knowledge and manipulation of that knowledge. However, the cognitive processes underpinning these two aspects of creative thought remain underspecified. The broader clinical-cognitive neuroscience literature suggests that retrieval and manipulation of knowledge is underpinned by general purpose cognitive mechanisms supporting semantic cognition, controlled episodic memory retrieval, and executive mechanisms. To identify commonalities from converging evidence that points towards a unifying theory for the neurocognitive bases of creative thought, we reviewed and meta-analysed 152 studies from creativity and the relevant parallel cognitive neuroscience literature using transcranial direct current stimulation (tDCS). The results revealed three things: 1) current tDCS studies are heavily biased towards the frontal cortex (459/591 effect sizes; 77.7%); 2) only anodal tDCS over the left lateral frontal cortex promotes creativity (p <.01); and 3) anodal tDCS stimulation over the same region also promotes improvement in many other cognitive processes. The latter includes more efficient processing of semantic knowledge (p <.05), more accurate episodic memory retrieval (p <.05), better and more efficient manipulation of buffered knowledge (all p <.001), and more efficient response selection amongst competing options (i.e., task-setting; p <.01). By merging these previously separate literatures, tDCS studies support the notion that creative thought arises from general purpose cognitive mechanisms including controlled retrieval and temporary storage of semantic and episodic information, as well as executive mechanisms.

## 1. Introduction

What are the cognitive and neural bases of creative thought, a fundamental capacity of humans that underpins everyday problem solving and scientific breakthroughs? Although this question is crucial to humanity, it remains largely unanswered. Early conceptualisations of creativity suggest it comprises a generative process (also known as divergent thinking; Guilford, 1967) and an evaluation-selection process (akin to convergent thinking; Campbell, 1960; Mednick, 1962). However, our current understanding of the cognitive and neural substrates underpinning creativity is limited by the common approach embraced in cognitive psychology, functional neuroimaging and related neuroscience fields, in which one specific cognitive domain is the sole focus of investigation (i.e., domain-specific). Rather than defining the specific cognitive processes underlying creativity, the widely adopted approach has been to use a restricted set of cognitive tasks for measuring creativity (**Table 1**).

**Table 1:** Tasks used in creativity research or when tapping cognitive processes implicated in creative thought – domain-specific vs domain-general approaches.

| Perspective | Tasks commonly used in experimental settings |  |  |  |
| --- | --- | --- | --- | --- |
| Domain-specific | Subset |  | Definition | Commonly used cognitive tasks |
|  | Divergent thinking |  | Generate a range of verbal or nonverbal ideas within limited time | • Alternate Uses Task (AUT; Guilford, 1967): participants are asked to generate various unconventional/original uses for familiar objects e.g. within two minutes (Gilhooly et al., 2007).<br>Better performance<br>• Fluency: generation of more unique and unconventional uses<br>• Originality: greater mean semantic distance of generated ideas (Chrysikou et al., 2021) |
|  | Convergent thinking |  | Produce a single solution to a list of problems/questions | Remote associates test (RAT; Mednick, 1962): generate the fourth word that is related to the three given cue words<br>Better performance: greater number of correct problems solved |
| Domain-general | Cognitive Cornerstone | Components | Definition | Commonly used cognitive tasks |
|  | Pre-existing knowledge | Semantic cognition | Flexible use of lifelong acquired knowledge about the world (e.g., names and properties of objects, relationships between concepts) | • Semantic representation: Picture/word matching, picture/letter/digit naming; Word/sentence reading (including stroop – congruent condition); semantic judgement (e.g., synonym judgement)<br>• Semantic control: semantic relatedness (amplification of less dominant aspects of a concept), semantic congruence (resolution of incongruent meaning) |
|  |  | Controlled episodic memory retrieval | New learning of verbal/visual information, as well as personal recollections | • Episodic memory: Paired associates learning; word list learning (recognition)<br>• Controlled episodic memory retrieval: word list learning (recall) |
|  |  | Multimodal knowledge buffer | Temporary memory storage for knowledge to be manipulated in a goal-directed manner | • Visuospatial/verbal working memory tasks<br>• N-back paradigms (semantic/less semantic) |
|  | Executive mechanisms | Energization | Self-initiation and sustaining response generation in the absence of cues | • Word generation tasks (semantic/phonemic verbal fluency, ideational fluency (i.e., generation of conventional/unconventional uses of daily objects)<br>• Nonverbal response generation tasks |
|  |  | Task-setting | Selecting task-relevant responses from competing responses by establishing stimulus-response relationship | • Set shifting (repeat condition)<br>• Stroop (incongruent)<br>• Flanker (incongruent)<br>• Attention network test [incongruent/(incongruent-congruent)]<br>• Choice RT tasks |
|  |  | Monitoring | Checking of responses and ensuring the responses are task-relevant by inhibiting task-irrelevant responses | • Go/no-go (no-go trial)<br>• Stop signal task (stop-signal reaction time)<br>• Continuous performance test<br>• Set shifting (shift condition)<br>• Post error slowing in task-setting/other EF tasks |
|  |  | Self-regulation | Processing risk/reward | • Risk-taking tasks (e.g., Balloon Analogue Risk Task)<br>• Intertemporal Choice task |
|  |  | Metacognition | Integrating self-regulatory and executive functions | • Confidence ratings in memory/perceptual tasks |

Empirical findings yielded from these tasks suggest that creativity may involve at least two cognitive aspects – 1) pre-existing knowledge that provides “raw materials” for idea generation (Beaty & Kenett, 2023; Benedek et al., 2023; Cropley, 2006; Dane & Pratt, 2007; Ericsson et al., 2007; Sternberg, 1988), and 2) an operational system that flexibly manipulates ideas in a goal-directed manner (Abraham, 2019; Benedek & Fink, 2019; Chrysikou, 2019; Flaherty, 2005; Gruber & Wallace, 1999; Lebuda & Benedek, 2023; Mekern et al., 2019).

Furthermore, these findings suggest that creativity primarily recruits fronto-parietal-temporal brain regions (Gonen-Yaacovi et al., 2013; Kuang et al., 2022). However, when considering contributing cognitive domains and the broader clinical-cognitive neuroscience literature, it becomes apparent that the tasks listed in **Table 1** are not specific to creativity. Instead, these tasks tap multiple cognitive processes, and most (if not all) of these processes serve general purpose cognitive mechanisms in support of knowledge sources – semantic cognition (Lambon Ralph et al., 2017) and episodic memory (Madore et al., 2015), and executive control functions (Robinson et al., 2012; Stuss 2011).

To understand the cognitive and neural foundations of creative thought, the first necessary step is to define the specific processes that underly creative thought based on empirical theories. Therefore, from the broader clinical-cognitive neuroscience perspective, we have proposed that creative thought is supported by two core “cognitive cornerstones” (Chan et al., 2023). These cornerstones are supported by domain general neurocognitive mechanisms: 1) pre-existing knowledge (i.e., semantic and episodic memories), the source material from which knowledge can be flexibly retrieved and temporarily “buffered” for further manipulation, and 2) executive mechanisms that manipulate knowledge in a goal-directed manner. The latter executive mechanisms entail energization (i.e., self-initiation and sustaining of responses in the absence of external cues), task-setting (i.e., selection of goal-relevant responses amongst competing/distracting stimuli), monitoring (i.e., anticipation of errors and checking of ongoing task performance by inhibiting goal-irrelevant responses), self-regulation (i.e., motivation/risk/reward processing), and metacognition (i.e., integration of motivational and executive capacities). Some tasks are well established for assessing the components of these two cornerstones (**Table 1**).

An important next step is to analyse the large quantity of empirical findings across specific cognitive domains by means of meta-analytic methods (Humphreys & Lambon Ralph, 2015; Poldrack & Yarkoni, 2016), in order to identify the converging evidence and search for a unifying explanation. In this study, we focussed on transcranial direct current stimulation (tDCS) studies because tDCS enables the investigation of the causal relationships between the activity of a brain region and behaviour (Bradley et al., 2022; Miniussi et al., 2013). tDCS is a non-invasive neuromodulation technique operated by delivering a constant direct current that flows between electrodes (Thair et al., 2017). The direct current temporarily excites (by anodal stimulation) or suppresses (by cathodal stimulation) cortical neuronal populations of one or more brain region(s) and hence facilitates or inhibits cognitive processes associated with that region (Polanía et al., 2018). In parallel literatures, tDCS has been widely used in both the creativity domain (Lucchiari et al., 2018) and other cognitive domains (Narmashiri & Akbari, 2023). Meta-analysis is an especially useful approach to adopt because the tDCS creativity studies have (i) typically comprised small sample sizes with diverse participants’ demographic background (Ivancovsky et al., 2019) and (ii) utilised different brain stimulation protocols/outcome measures. The extant individual studies suggest that frontal (Mayseless & Shamay-Tsoory, 2015), parietal (Pick & Lavidor, 2019) and temporal (Ruggiero et al., 2018) brain regions are all necessary for spontaneous production of novel ideas and goal-directed evaluation-selection of appropriate ideas. In parallel, numerous tDCS cognitive neuroscience studies indicate that similar frontal, parietal and temporal regions are implicated in cognitive processes that support semantic cognition (Joyal & Fecteau, 2016), episodic memory (Bjekić et al., 2019; Galli et al., 2019), and executive control functions (de Boer et al., 2021; Schroeder et al., 2020). Thus, parallel examination of creativity and other relevant cognitive neuroscience tDCS studies offers the opportunity to identify the shared and distinct cognitive processes, and associated brain regions, that underlie creative thought.

In this study, we began with systematically sourcing and summarising tDCS studies that employed tasks tapping creativity and relevant cognitive domains (semantic cognition, controlled episodic memory retrieval, and executive mechanisms). We then classified these studies by tDCS-targeted brain regions (frontal, parietal, temporal) and stimulation polarity (anodal, cathodal, bilateral). Two parallel meta-analyses – one for the creativity literature and another for the cognitive neuroscience literature – were conducted to assess which brain regions, when stimulated, result in significant enhancement or disruption of creativity or domain general cognitive processes, indicating that the targeted brain region is critical for that process. If any brain region targeted by tDCS for enhancing or disrupting creativity overlaps with brain regions identified as essential for the cognitive cornerstones, this overlap would provide evidence supporting our hypothesis: creative thought arises from general purpose cognitive mechanisms supporting semantic cognition, controlled episodic memory retrieval, and executive functions.

## 2. Methods

### 2.1 Literature Search

This study was conducted in accordance with the Preferred Reporting Items for Systematic Reviews and Meta-Analyses (PRISMA) 2020 guideline (Page et al., 2021). In order to obtain tDCS studies on creativity and cognition, multiple literature search methods were adopted.

To obtain tDCS studies on creativity, candidate search terms were identified by reading relevant reviews (Benedek et al., 2016; Chrysikou, 2019; Lucchiari et al., 2018). Text mining was performed using PubReminer for keyword refinement (O’Mara-Eves et al., 2015) with the search terms reported in the previous reviews mentioned above. A preliminary search using Scopus was conducted in October 2022 to confirm the search strategy. Formal electronic database search using Embase, MEDLINE, Web of Science and PsyINFO was conducted on 6^th^ November 2022 with the following keywords: creative* AND (process* OR thought* OR think* OR executive OR semantic OR cognit* OR control) AND (brain OR neural OR neuroscien* OR neurolog*). No limitations were set on publication dates. Search strategies for each of the databases were listed in **Table S1**. To obtain up-to-date records, an additional search was performed using Google Scholar (year of publication: 2023 – present) with the above search terms in October 2024 with the first 300 records (Haddaway et al., 2015) being screened. We also searched for eligible records from a recently published meta-analysis of tDCS creativity studies (Chen et al., 2024).

Published tDCS studies on cognition were obtained from the reference lists of the eight published reviews (de Boer et al., 2021; Jacobson et al., 2012; Joyal & Fecteau, 2016; Ostrowski et al., 2022; Schroeder et al., 2020; Strobach & Antonenko, 2017; Tremblay et al., 2014; Westwood & Romani, 2017). These tDCS reviews collectively provide a comprehensive set of empirical studies that examined the effects of tDCS on cognition in healthy individuals across various cognitive domains, including attention, memory, executive functions, decision making and language. While we primarily extracted studies from reviews in the memory and executive function domains, we also search for studies from broader domains because tasks tapping semantic cognition, episodic memory retrieval, and executive control functions sometimes appear in the attention, decision-making, and language literatures. As these reviews only included studies published until 2021, supplementary electronic database searches (conducted in August/September and December 2023) were conducted using PubMed, Web of Science and Google Scholar with publication dates limited (2022-present) in order to gather up-to-date relevant studies. Key search terms were: (“tDCS” OR “transcranial direct current stimulation”) AND “healthy” AND (“cognition” OR “attention” OR “working memory” OR “semantic memory” OR “episodic memory” OR “executive function” OR “self-regulation” OR “language”).

The titles and abstracts of records retrieved from the above search procedures were obtained and imported to Endnote 20 reference manager tool for further screening.

### 2.2 Article screening and inclusion

Published randomised/crossover, sham-controlled trials that investigated the effects of conventional/high-definition tDCS on creativity, as well as the putative cognitive processes underlying creative thought in healthy individuals [age between 18-40yrs to reduce heterogeneity of dataset contributed by developmental and aging factors (Hodgson et al., 2023)] were included. Titles and abstracts of the retrieved records were first screened to exclude the following: 1) duplicated records, 2) non-empirical studies (e.g., book chapters, reviews, commentaries), 3) animal studies, and 4) studies that did not involve the use of transcranial electrical stimulation. Then, full texts of the included records during title/abstract screening were retrieved. During full-text screening, records were first screened to see whether they fulfil these inclusion criteria: 1) randomised/crossover and sham-controlled studies; 2) studies that applied conventional tDCS or high-definition tDCS (HD-tDCS) protocols on healthy individuals aged between 18-40yrs; 3) studies that used well-established cognitive tasks to measure creativity or the proposed general purpose cognitive mechanisms (i.e., cognitive cornerstones) implicated in creative thought (**Table 1**), 4) studies that measured “offline” or “online” tDCS effects (**Figure 1a**).

To focus on the fronto-parieto-temporal involvement in putative cognitive processes underlying creative thought, we only included studies that fulfil the above inclusion criteria, as well as with 5) (HD-)tDCS stimulation applied over the predefined fronto-parieto-temporal brain regions shown to be associated with creative thought (**Figure 1b**). In addition, we only included studies that reported objective (but not subjective, e.g., confidence ratings) behavioural outcome measures (i.e., accuracy/error rates/reaction time). As a result, tDCS studies tapping metacognition [e.g., Bona and Silvanto (2014); Han et al. (2023)], which typically assess participants’ subjective ratings of their performance in memory/perceptual tasks, were excluded from this review.

### 2.3 Data extraction and recoding

From each included study, design (i.e., crossover/parallel, sham-controlled, participant-blind and/or assessor-blind), demographic details of the participants, tDCS protocol, and outcome measures were extracted. Demographic data included the number of participants and their mean age. tDCS protocol data included stimulation modality (i.e., conventional tDCS or HD-tDCS), position of the anode and cathode (according to standard electrode position nomenclature; Acharya et al., 2016), or standardised brain templates; Evans et al., 2012), number of active/sham stimulation sessions, duration of stimulation, current intensity, and electrode size. The outcome measure data encompassed cognitive task names and task descriptions, outcome measure time frame (i.e., before/during/after stimulation), and the numerical values of outcomes and/or narrative descriptions of results.

Some extracted qualitative data was recoded into nominal data to facilitate subsequent narrative/quantitative synthesis. First, given previous meta-analyses showed that different tDCS stimulation polarities (e.g., anodal/cathodal stimulation) have differential effects on cognition (Jacobson et al., 2012; Narmashiri & Akbari, 2023), a variable “stimulation polarity” was created. To excite/suppress a brain region, the most common montage (i.e., unipolar anodal/cathodal stimulation) involves the placement of one stimulating (anodal/cathodal) electrode(s) over the scalp, and a returning electrode placed over the forehead just above the supraorbital ridge, or other extracephalic regions (e.g., upper arm). Recently, a montage that is believed to improve tDCS spatial focality has gained popularity (Datta et al., 2009) and involves the placement of one active electrode in the centre, surrounded by several returning electrodes (i.e., unipolar anodal/cathodal high-definition tDCS). Researchers have also investigated how cognitive processes are altered by simultaneously exciting a brain region and suppressing its contralateral hemisphere analogue (i.e., bilateral stimulation). Hence, the “stimulation polarity” variable contained three categories (i.e., anodal, cathodal, bilateral), which was created by recoding information about anode and cathode placement of each study based on established frameworks (Chan et al., 2021; Nasseri et al., 2015). For detailed recoding procedures, see **Figure 2a**.

Second, the cognitive tasks used by different studies may vary in the nature of stimuli and response type, of which the processing of these different stimuli has been shown to involve different brain regions (Robinson et al., 2012). To understand how this could potentially modulate the results, a set of dichotomous variables (semantic *vs.* less semantic, visual *vs.* auditory, verbal *vs.* nonverbal) describing the nature of task was created. Of note, a cognitive task presenting words/sentences, familiar faces, photos of famous places, familiar music, body language, cartoon, and well-rehearsed arithmetic problems (e.g., simple addition/subtraction) was classified as a “semantic” task; otherwise, it was classified “less semantic” (e.g., tasks presenting nonwords, alphabets and digits). Meanwhile, tasks that required the manipulation of words (e.g., overt/covert speech, typing, writing) were considered a verbal task; otherwise, it was regarded a nonverbal task.

### 2.4 Meta-analysis

Studies that examined differences between active– and sham-tDCS groups were included in the meta-analysis. The descriptive data/inferential statistics were used for effect size calculation. Effect sizes (Cohen’s *d*) were calculated using online calculators (Lenhard & Lenhard, 2022; Uanhoro, 2017; Wilson, 2023); for detailed effect size (and variance) calculation procedures, see **Figure 2b**. These procedures are commonly adopted in meta-analyses (e.g., Fox et al., 2016). However, for studies that did not report minimal information for effect size calculation, our approach would tend to underestimate the actual effects.

Due to the diversity of tDCS montages used across studies, it is essential to categorise effect sizes meaningfully to enable a robust meta-analysis. Previous cognitive neuroscience literature suggests that medial and lateral frontal/temporal/parietal regions are differentially associated with semantic cognition (Jackson, 2021), episodic memory (Daviddi et al., 2023) and executive control mechanisms (Domenech et al., 2020; Stuss, 2011). Thus, effect sizes were grouped by stimulation polarity (anodal, cathodal, bilateral) across predefined regions of interest (ROIs; **Figure 1b**) – left/right lateral frontal cortex, medial frontal cortex, left/right lateral parietal cortex, medial parietal cortex, and left/right lateral temporal cortex. To ensure consistency, 1) only studies/experiments that involved the stimulation of 1 ROI were included in the meta-analysis, and 2) for bilateral stimulation, only studies that involved left anodal (L+)/right cathodal (R-) or left cathodal (L-)/right anodal (R+) stimulation over the same lobe (frontal/parietal/temporal) were analysed.

To examine the brain regions necessary for creative thought from the perspective of tDCS, two parallel meta-analyses were conducted. The first series of analyses on the creativity literature examined the tDCS-induced changes in creativity task performance (i.e., divergent and convergent thinking). The second series of analyses on the broader cognitive neuroscience literature examined the tDCS-induced changes (as reflected by accuracy and reaction time data) in domain general cognitive mechanisms (i.e., controlled retrieval of semantic knowledge and episodic memory, manipulation of buffered knowledge, energization, task-setting, monitoring, and self-regulation). A random-effects model was used for pooling and analysing the aggregated effect sizes. Pooled effects for the analyses were considered significant at p = .05 (uncorrected) level. A more liberal threshold was chosen due to the conservative effect size calculation, which likely underestimates the true tDCS effect. This decision also accounts for the small effect sizes typically observed and the limited statistical power of individual studies in the tDCS literature.

For montages where significant meta-analytic results emerged for both creativity and broader cognitive neuroscience studies, sensitivity analyses were performed to see if the effect sizes differ between study designs and nature of tasks. Specifically, Welch robust tests of equality of means were performed to examine whether the significant tDCS-induced effects were biased by 1) semantic/less semantic, 2) verbal/nonverbal, 3) online/offline tasks, or 4) the use of HD-tDCS/conventional tDCS. Between-study heterogeneity in outcome measures (creativity task performance/ACC/RT) was measured with the I-squared (I^2^) statistics (Borenstein et al., 2021), and the level of heterogeneity was classified based on Higgins et al. (2003), with I^2^ values of 25 %, 50 % and 75 % being considered indicative of low, medium, and high heterogeneity, respectively. To support the development of better-designed and adequately powered studies on the neurocognitive basis of creativity using tDCS, we calculated the minimum required sample size for repeated-measures design to detect a significant difference between active and sham tDCS groups. This calculation was based on the averaged Cohen’s d value for creativity tasks, with α = .05 (two-tailed), power = .8, number of measurements = 2 (i.e., single-session active vs. sham tDCS), and correlation among repeated measures = .5. Publication biases were visually inspected using funnel plots, which plotted the effect sizes of all included studies in the meta-analysis on the x-axis and standard error on the y-axis. In the presence of publication bias, the plot is expected to be symmetrical at the top, while an increasing number of data points are missing from the middle to the bottom of the plot (Borenstein, 2022).

Random-effects meta-analyses, between-study heterogeneity, and the evaluation of risk of publication bias were performed using IBM SPSS Statistics for Windows, version 29.0 (IBM Corp., Armonk, N.Y., USA). Sample size calculation was conducted using G*power version 3.1 (Faul et al., 2009).

## 3. Results

### 3.1 Study characteristics

The number of studies included in each stage of the article screening procedure is reported in **Figure 3**. When combining the studies yielded from all search methods, 9554 title/abstract records were screened, with 273 full-texts retrieved and examined. With inclusion and exclusion criteria applied, a total of 172 crossover/parallel, randomised, sham-controlled studies (20 creativity and 152 cognitive neuroscience studies) were included.

Within this set, all creativity studies and 130 cognitive neuroscience studies were eligible for meta-analysis. A total of 591 effect sizes were extracted and included in the meta-analysis (**Figure 4**). Of note, most of these effect sizes were derived from frontal stimulation studies (459 out of 591; 77.7%). Effect sizes from parietal (17.7%) and temporal (4.6%) stimulation studies were substantially less common.

Study design, participants’ demographic details, tDCS protocol and outcome measures of each of the studies included in this review are reported in **Table S2**. All creativity studies presented stimuli visually, and most of the studies involved the use of verbal semantic tasks (i.e., alternate uses task/remote associates test; 43 out of 52 effect sizes, 83%). Four studies [contributing six effect sizes; Aihara et al. (2017); Bartel et al. (2020); Chi and Snyder (2012); Luft et al. (2017)] involved the use of nonsemantic nonverbal tasks (e.g., matchstick problem solving, nine dot problem, inkblot image processing). Only two studies [contributing three effect sizes; Guo et al. (2023); Ruggiero et al. (2018)] involved the use of nonverbal semantic tasks (i.e., draw a picture within a frame and assign a caption to it, and draw multiple designs with annotations). Almost all cognitive neuroscience studies presented task stimuli visually. While tasks assessing semantic cognition and episodic memory were typically verbal in nature, tasks evaluating executive processes were generally nonsemantic and nonverbal.

### 3.2 Meta-analytic results

An overall summary of the meta-analytic results is shown in **Figure 5**. It illustrates the differential effect of tDCS stimulation polarity, montage, and cortical region (frontal, parietal, temporal) on various domain-specific and domain-general processes implicated in creativity. Frontal stimulation showed the strongest effects on both creativity tasks and cognitive processes underpinning creative thought, although this may partly reflect the disproportionate number of effect sizes from frontal studies. For parietal stimulation, meta-analysis was not performed for creativity studies as only one effect size was available. Nevertheless, one parietal stimulation montage showed some isolated effects on domain-general cognitive processes implicated in creative thought. In contrast, temporal stimulation results were largely nonsignificant.

#### 3.2.1 Frontal stimulation

Despite the variety of montages tested in the frontal region, only anodal stimulation over the left lateral frontal cortex consistently yielded improvements. These improvements were observed in both domain-specific processes of creativity and the proposed domain-general cognitive mechanisms underpinning creative thought. As reflected in the forest plot (**Figure 6**), anodal left lateral frontal stimulation does not only promote divergent thinking (*z* = 1.962, *d* = .540, 95% CI [.00004, 1.08], *p* = .0498) and convergent thinking (*z* = 2.095, *d* = .401, 95% CI [.026, .776], *p* = .036), but it also promoted behavioural improvements in many other domain general cognitive processes. The latter improvements included 1) more efficient processing of semantic knowledge (reaction time: *z* = –2.345, *d* = – .422, 95% CI [-.775, –.069], *p* = .019), 2) more accurate episodic memory retrieval (accuracy: *z* = 2.888, *d* = .240, 95% CI [.077, .402], *p* = .004), 3) better and more efficient manipulation of buffered knowledge (accuracy: *z* = 3.336, *d* = .213, 95% CI [.088, .338], *p* < .001; reaction time: *z* = –3.599, *d* = –.235, 95% CI [-.362, –.107], *p* < .001), as well as 4) more efficient response selection amongst competing options (i.e., task-setting; reaction time: *z* = –2.658, *d* = –.317, 95% CI [-.551, –.083], *p* = .008). However, broader cognitive neuroscience studies indicated that other montages also yield improvements in various cognitive processes beyond creativity. Specifically, right lateral and medial frontal anodal stimulation promoted more efficient task monitoring (right lateral frontal ROI p < .001, **Figure S1**; medial frontal ROI p = .002, **Figure S2**), bilateral (left-anodal/right-cathodal) frontal stimulation promoted more efficient processing of semantic knowledge (p = .028, **Figure S3**), bilateral (left-cathodal/right-anodal) frontal stimulation promoted more efficient task-setting (p = .029, **Figure S4**), and left lateral frontal cathodal stimulation promoted more accurate episodic memory retrieval and monitoring (all p < .05, **Figure S5**). Interestingly, right lateral and frontal cathodal stimulation specifically disrupted efficiency in task-setting (p = .028, **Figure S6**), while medial frontal cathodal stimulation had no significant effects on domain general cognitive processes (all p > .11; **Figure S7**).

#### 3.2.2 Parietal stimulation

Anodal stimulation over the left lateral parietal cortex consistently promoted more accurate episodic memory retrieval (p = .035, **Figure S8**). Other parietal montages showed no significant effect on domain general cognitive processes (all p > .25, **Figures S9-S13**).

#### 3.2.3 Temporal stimulation

Anodal stimulation over the right lateral temporal cortex (**Figure S15**) and bilateral lateral temporal stimulation (**Figures S16–S17**) showed no significant effects on creativity task performance (all p > .32). Similarly, tDCS studies in cognitive neuroscience yielded nonsignificant findings.

### 3.3 Additional analysis

As only anodal stimulation over the left lateral frontal cortex observed significant results from both the creativity and cognitive neuroscience literatures, sensitivity analysis was only conducted for this subset of effect sizes. Regarding the creativity literature, while the results were heavily biased towards visual tasks (noted above; see **Table S2**), a Welch test indicated that the observed significance was not significantly biased by verbal/semantic tasks (F_1,1.20_ = .007, p = .944), online measures (F_1,1.64_ = .268, p = .666), and the use of HD-tDCS (F_1,6.91_ = .990, p = .353). Regarding the cognitive neuroscience literature, when comparing effect sizes from semantic vs. less semantic tasks, effect sizes reflecting accuracy (F_1,77.44_ = 1.16, p = .285) or reaction time (F_1,36.88_ = .207, p = .652) changes did not differ significantly. When comparing effect sizes from verbal vs. nonverbal tasks, effect sizes reflecting accuracy (F_1,80.59_ = 1.68, p = .199) or reaction time (F_1,32.58_ = .179, p = .675) differences between active and sham tDCS groups were not statistically different. When comparing effect sizes from online vs. offline measures, effect sizes reflecting accuracy (F_1,70.53_ = .187, p = .666) or reaction time (F_1,57.15_ = 1.23, p = .273) changes also did not differ significantly. Similarly, the observed significance was not significantly biased by the use of HD-tDCS (ACC: F_1,6.26_ = .013, p = .911; RT: F_1,4.39_ = 1.82, p = .242).

The between-study heterogeneity (I²) in creativity studies varied across montages, ranging from <1% (very low heterogeneity) to 67% (moderate heterogeneity). The variability in heterogeneity across montages was even greater for cognitive neuroscience studies, ranging from <1% (very low heterogeneity) to 91% (high heterogeneity). Sample size calculation (based on an averaged Cohen’s d of 0.25 for all creativity studies) indicates the following: if one uses a repeated-measures design, at least 98 participants (i.e., n = 49 per group) are needed to detect a statistically significant difference between active and sham tDCS groups. Visual inspection of funnel plots did not reveal apparent publication bias (**Figure 7**).

## 4. Discussion

The neurocognitive mechanisms underlying many higher cognitive functions, including creative thought, remain elusive. In cognitive psychology and related cognitive neuroscience fields, to study how the brain supports a particular cognitive function, researchers often select a narrow set of tasks that are thought to tap a particular cognitive ability. Yet a century of clinical-cognitive neuroscience research has repeatedly shown that it is probably inadequate to investigate the mechanisms of a cognitive activity using a single (or few) cognitive tasks, because most (if not all) of these activities engage multiple distinct but partially interactive cognitive processes. Thus, distilling what processes are necessary for a cognitive function can be challenging. In this study, we conducted a parallel examination of findings from tDCS studies concerning creative thought across different research subfields (i.e., creativity vs. clinical-cognitive neuroscience). Using meta-analytic methods, we aimed to identify converging evidence and search for a unifying explanation for the cognitive and neural bases of creative thought. Our results indicated that 1) the left lateral frontal cortex is necessary for creativity, evident by improved creative performance (via anodal tDCS stimulation) when participants perform divergent/convergent thinking tasks; and 2) the left lateral frontal cortex is also associated with distinct cognitive processes implicated in semantic cognition, controlled episodic memory retrieval and executive control mechanisms. Further, enhancing the cortical excitability in this region by anodal tDCS not only promotes better and more efficient controlled retrieval of long-term representations and manipulation of temporarily stored information, but it also promotes more efficient selection of appropriate response by filtering competing ideas (i.e., task-setting). We discuss these results from the clinical-cognitive neuroscience perspective of creative thought and suggest future study directions.

Consistent with previous clinical-cognitive neuroscience findings (Heo et al., 2023; Robinson et al., 2012), the creativity literature suggests that the left lateral frontal cortex is necessary for novel idea generation and goal-directed idea evaluation-selection, the characteristic features of creative thought (Christoff et al., 2016). Of note, the included studies are heavily biased towards the use of verbal semantic tasks. Classically neuropsychological (Baker et al., 2010; Fridriksson et al., 2011), brain stimulation (Cattaneo et al., 2011; Fertonani et al., 2010) and neuroimaging (Hodgson et al., 2023) studies have tended to suggest that the left lateral frontal cortex is associated with language functions, while the right lateral frontal cortex is associated with nonverbal novel idea generation (Robinson et al., 2012). Following this traditional framework, one could contemplate the possibility that the left lateral frontal cortex is specific to verbal, but not nonverbal, novel idea generation. Interestingly, when we examined the cognitive processes associated with the left lateral prefrontal cortex by considering the tDCS studies tapping semantic cognition, episodic memory and executive control functions, we found that this region is not only involved in verbal, but also nonverbal and less semantic tasks: there were equivalent effect sizes for semantic vs. non-semantic tasks, and for verbal compared to nonverbal tasks. Thus unlike the traditional discrete notions, these results imply that the left lateral prefrontal cortex might be necessary for both verbal and nonverbal novel idea generation – and aligns with the broader cognitive-clinical neuroscience literatures on semantic control and executive functions (e.g., Hodgson et al., 2023; Corbett et al., 2009; Thompson et al., 2016; Humphreys & Lambon Ralph, 2017) which implicate (a) a more bilateral picture (albeit with graded hemispheric asymmetries) and (b) multimodal engagement by left frontal areas (Assem et al., 2024; Jung et al., 2022). To examine whether or not the left lateral prefrontal cortex is specific to verbal novel idea generation, future neuropsychological and neuromodulation studies should examine whether the left lateral prefrontal cortex enhances performance in nonverbal novel idea generation tasks.

Although tDCS is widely adopted in both creativity and clinical-cognitive neuroscience research, the tasks used in creativity research are rarely used in cognitive neuroscience investigations (**Table 1**) and thus, findings tend to be considered in parallel isolation. An important question regarding the cognitive and neural bases of creative thought remains unanswered – which brain regions are necessary for the specific cognitive processes implicated in creative thought? To date, there is no single tDCS creativity study that attempts to address this question. To address this question, we adopted an alternative approach – we meta-analysed both the creativity and clinical-cognitive neuroscience literatures in parallel. By integrating these meta-analytic results, we at least partially answer this question. More importantly, through comprehensively reviewing the literature from these different fields (**Table S2**), we can identify the research gaps and plan for future studies that bridge these gaps. Consistent with the findings of a recent meta-analysis on creativity and tDCS (Chen et al., 2024), the left lateral frontal cortex is shown to be essential for creativity. Through the perspective of clinical-cognitive neuroscience, we further reveal that the left lateral prefrontal cortex is associated with some, but not all, of the cognitive processes implicated in creative thought; namely, the 1) *controlled retrieval of long-term representations* (i.e., semantic and episodic memories), 2) *manipulation of temporarily stored information*, and the 3) *selection* of goal-relevant ideas amongst competing ideas/distractors (i.e., *task-setting*). These findings are largely consistent with lesion (Stuss, 2011; Stuss and Alexander, 2007) and neuroimaging studies (Jackson, 2021; Vatansever et al., 2021; Wang et al., 2019).

Broadly speaking, the meta-analytic results confirm our “cognitive cornerstones hypothesis” – creative thought recruits the same brain region that gives rise to general purpose cognitive mechanisms supporting controlled retrieval of stored semantic and episodic memory, as well as executive control mechanisms. However, current tDCS studies are heavily biased towards the frontal cortex, which limits our understanding of the roles played by other brain regions. To address this gap, future research using brain stimulation techniques, including HD-tDCS and transcranial magnetic stimulation (for better localisation of brain regions), is needed for determining the causal roles of the temporal and parietal cortices on creativity performance and the general purpose cognitive mechanisms underpinning creative thought.

This study serves as a worked example demonstrating how parallel meta-analyses of tDCS studies from different subfields could guide the discovery of unifying explanations regarding the cognitive and neural bases of a cognitive activity (as has been done with ALE meta-analysis of fMRI data: Humphreys & Lambon Ralph, 2023; Humphreys et al., 2024; Chan et al., 2025). We believe that this approach not only applies to the understanding of creative thought, but also many other higher cognitive abilities e.g., working memory.

Although our results have provided important insights about the brain regions necessary for the cognitive processes implicated in creative thought, some limitations should be noted. First in terms of study inclusion, we excluded tDCS studies that examined metacognition as these works primarily relied on subjective measures that were not standardised across studies.

Future empirical tDCS studies are needed to investigate the interplay between metacognitive processes and creative thought. Specifically, studies that include parallel assessments of creativity and metacognition could help clarify which brain regions are essential for which metacognitive processes underpinning creative thought. Secondly in terms of effect size calculation, we adopted a conservative approach for studies that did not report minimal information for effect size calculation. Most studies did not report the pre-/post-stimulation correlation coefficient, and assuming a fixed correlation of 0.5 could introduce potential bias. To avoid overestimation, we took a more cautious stance, treating pre-post assessments as independent when variance could not be directly calculated. This assumption increased variance, reduced statistical power, and consequently led to more conservative pooled effect size estimates. As a result, our conservative approach likely underestimated the true effect sizes and impacted how study imprecision was evaluated. Thirdly based on our sample size calculation, future studies should consider having larger sample sizes. For instance, we calculated that creativity studies may require more than 49 individuals per group for detecting significant differences with single session tDCS. One way to reduce the number of participants required is the consideration of applying multi-session tDCS, which usually yields a larger effect size given the accumulative effect of tDCS across sessions (Song et al., 2019).

## 5. Conclusion

Through systematically reviewing and meta-analysing the creativity and cognitive neuroscience tDCS literatures, we aimed to understand some of the key cognitive processes associated with brain regions necessary for creative thought. Our results showed that the left lateral frontal cortex is necessary for creative thought, and that this brain region is associated with general purpose cognitive mechanisms implicated in creative thought. While these results support the notion that creative thought arises from the interplay between controlled knowledge retrieval (including semantic and episodic memory), flexible manipulation of buffered knowledge, and executive control functions, much more must be done to deepen our understanding regarding the neurocognitive bases of creative thought – across multiple cognitive mechanisms and their associated neural bases. This includes exploring the role of the left lateral frontal cortex on nonverbal and less semantic creativity tasks, how controlled retrieval, flexible knowledge manipulation and executive mechanisms interact to support creative thought, and how temporal/parietal brain regions are implicated in the cognitive processes underpinning creative thought.

## 6. Acknowledgements

None.

## 7. Funding sources

This research was supported by the Australian Research Council grant (DP220103941) awarded to GAR and MALR. MALR is supported by intramural funding from the UKRI-MRC (MC_UU_00005/18). The authors have no relevant financial or non-financial interests to disclose.

## 8. Data availability

The dataset generated and analysed during the current study is available on GitHub repository, https://github.com/melodymychan/CreativeThoughtTDCS.git

## 9. Authors’ contribution

M.Y.M.C.: conceptualisation, methodology, data curation, formal analysis, software, visualisation, writing – original draft; E.C.: data curation; M.A.L.R.: conceptualisation, supervision, funding acquisition, writing – review and editing; G.A.R.: conceptualisation, supervision, funding acquisition, writing – review and editing.

---

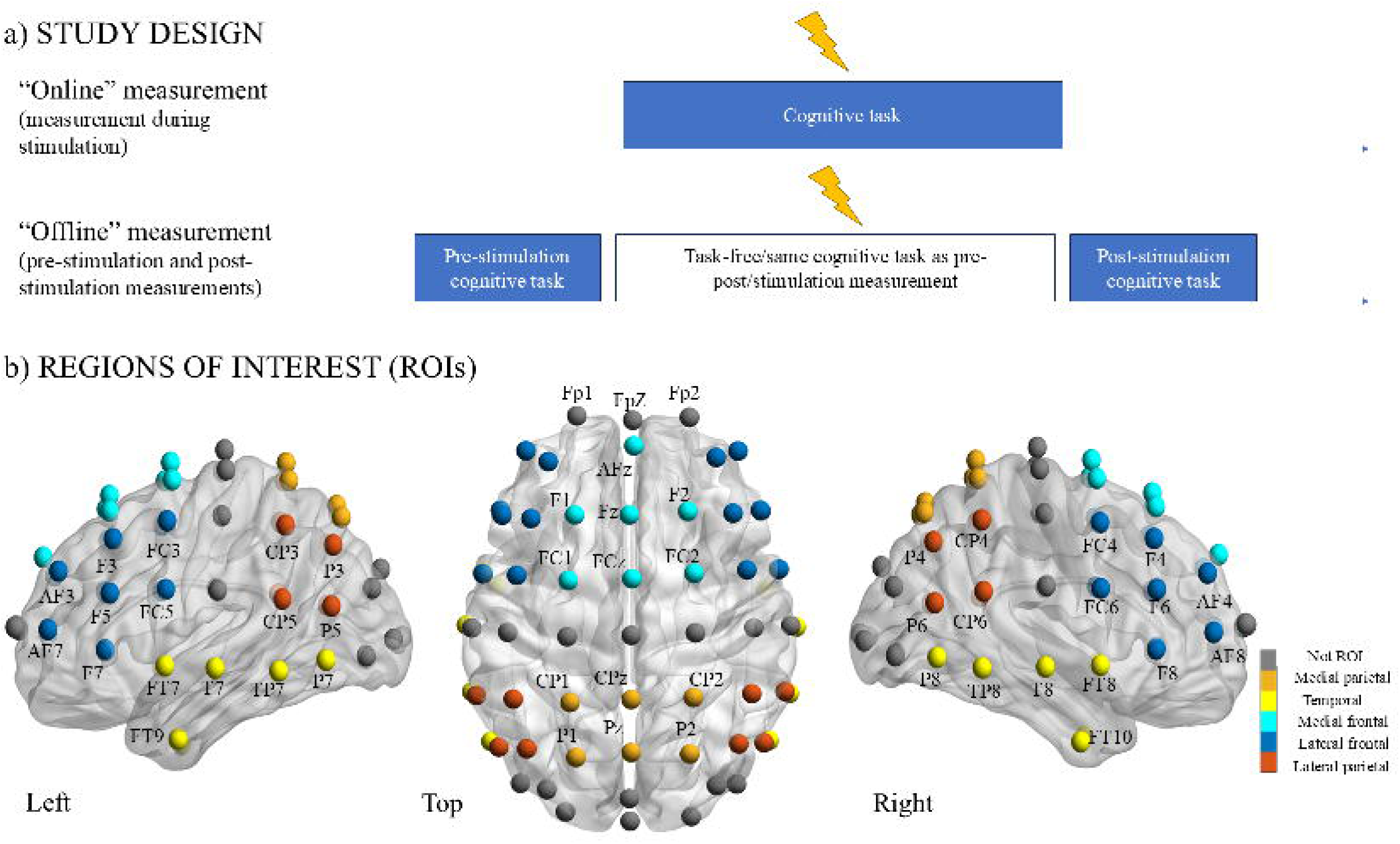

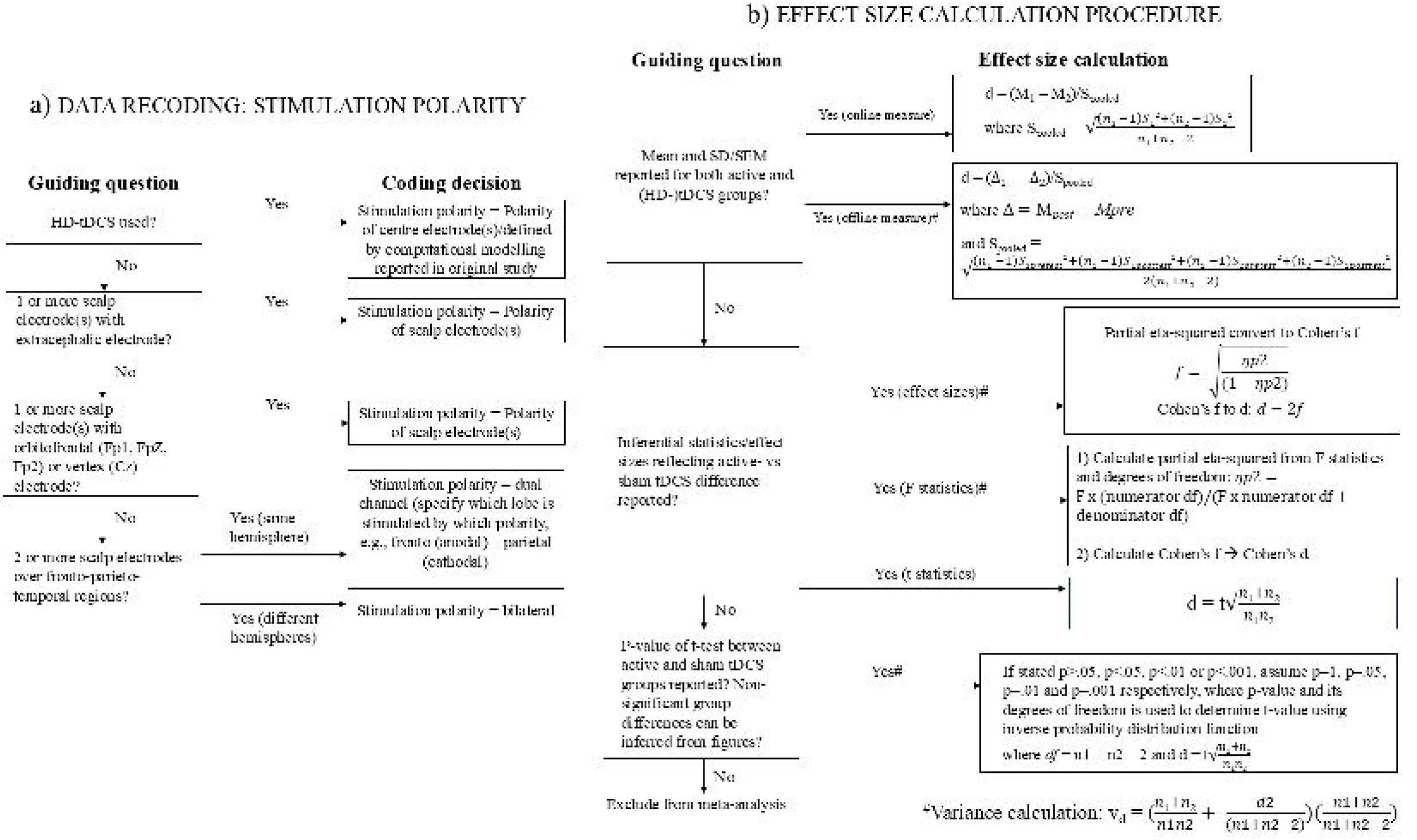

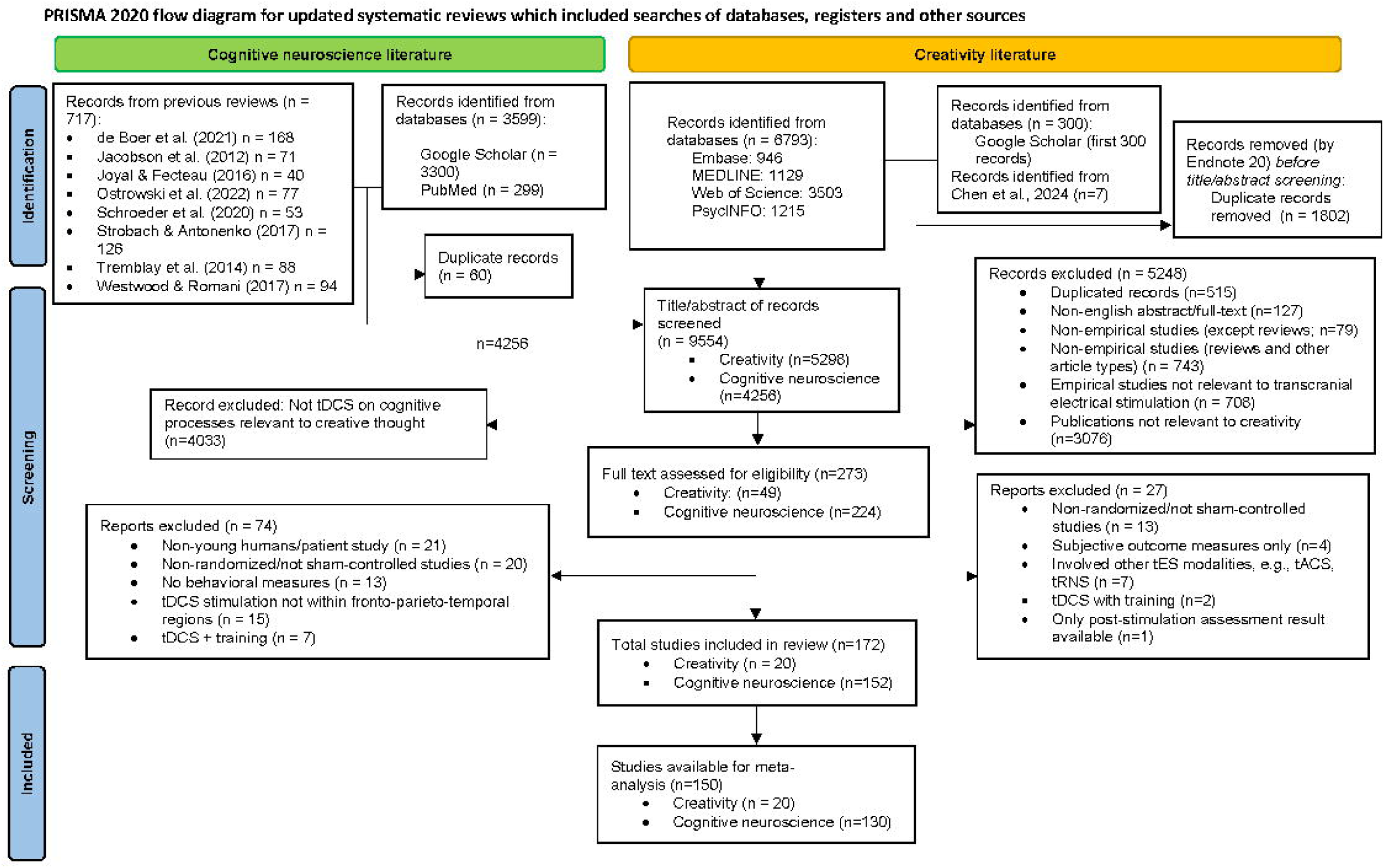

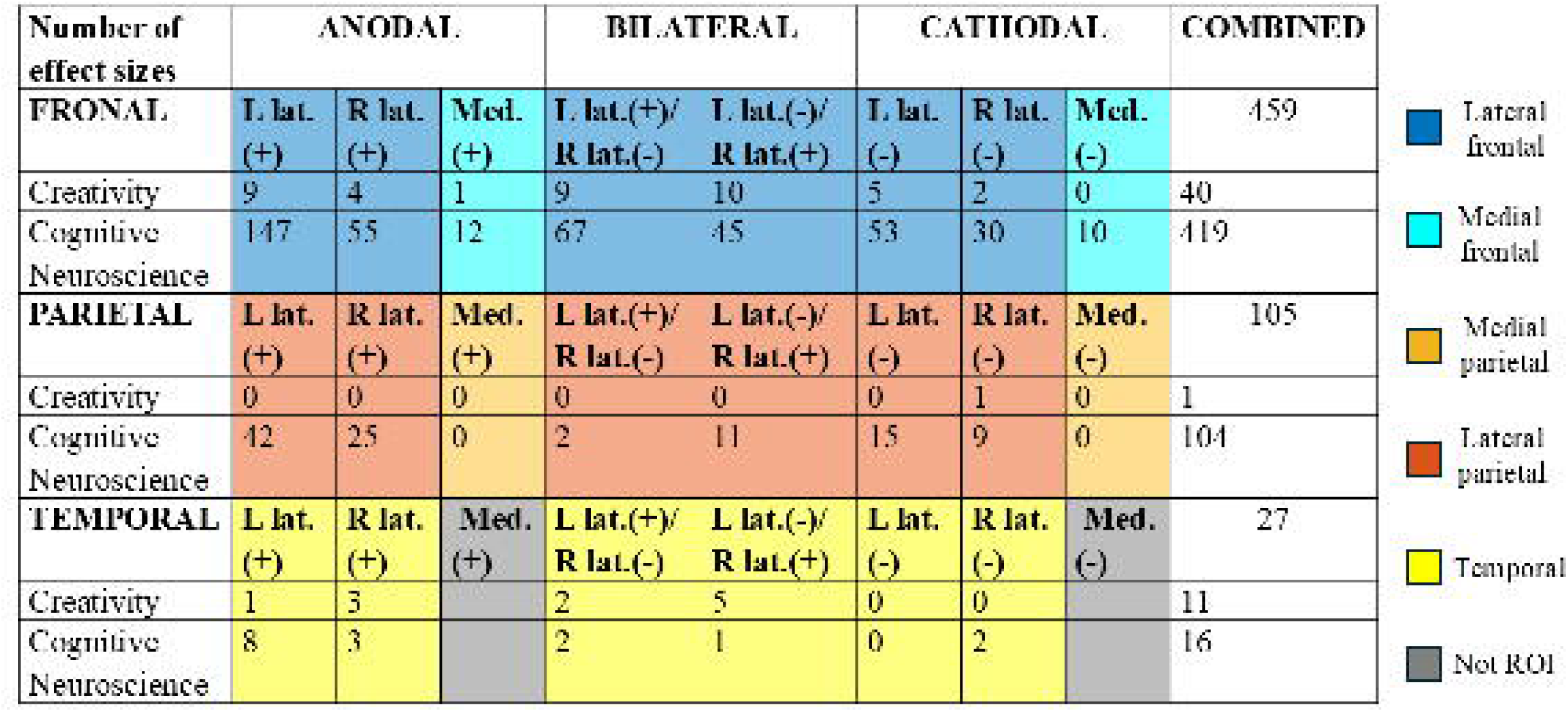

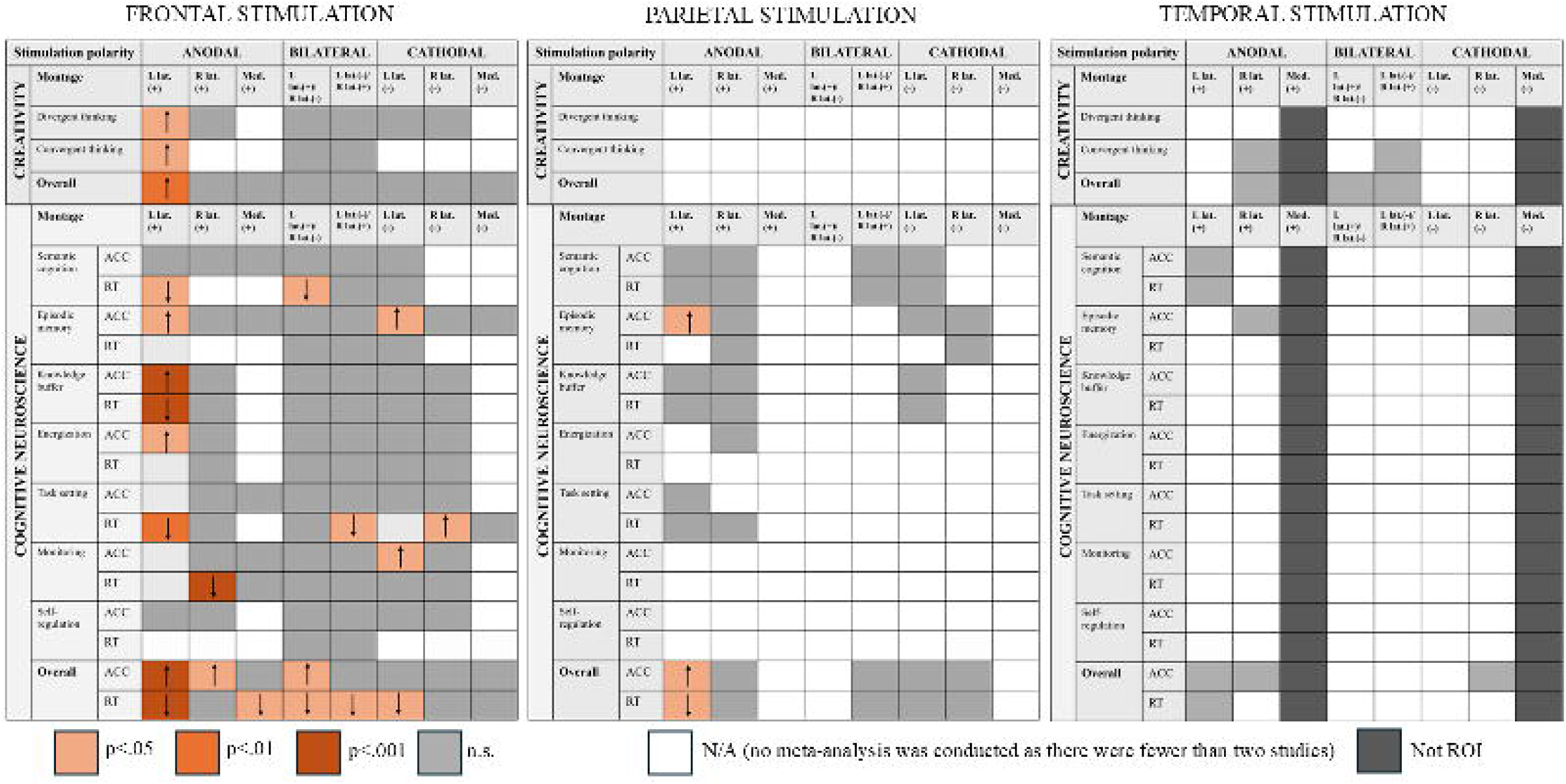

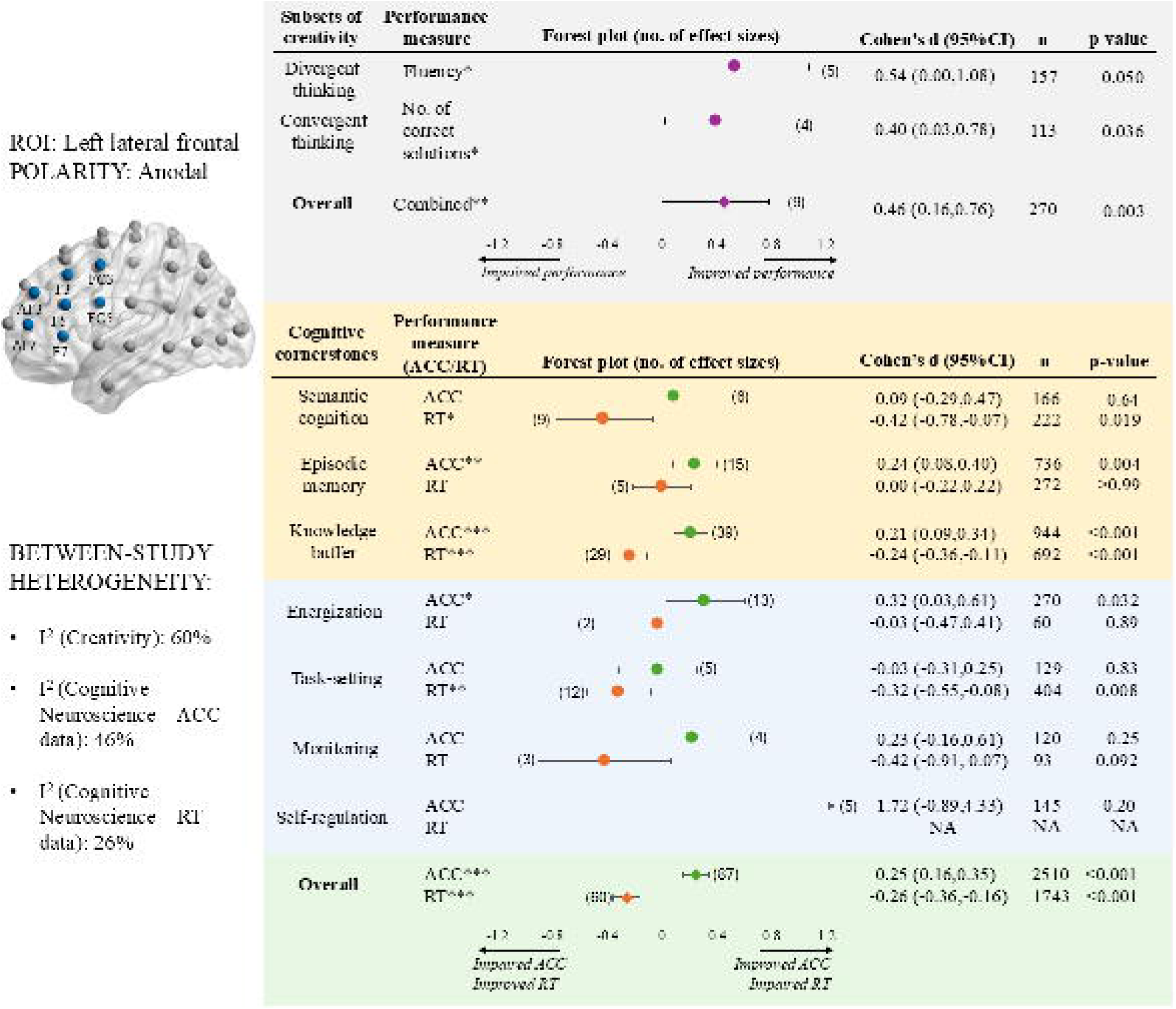

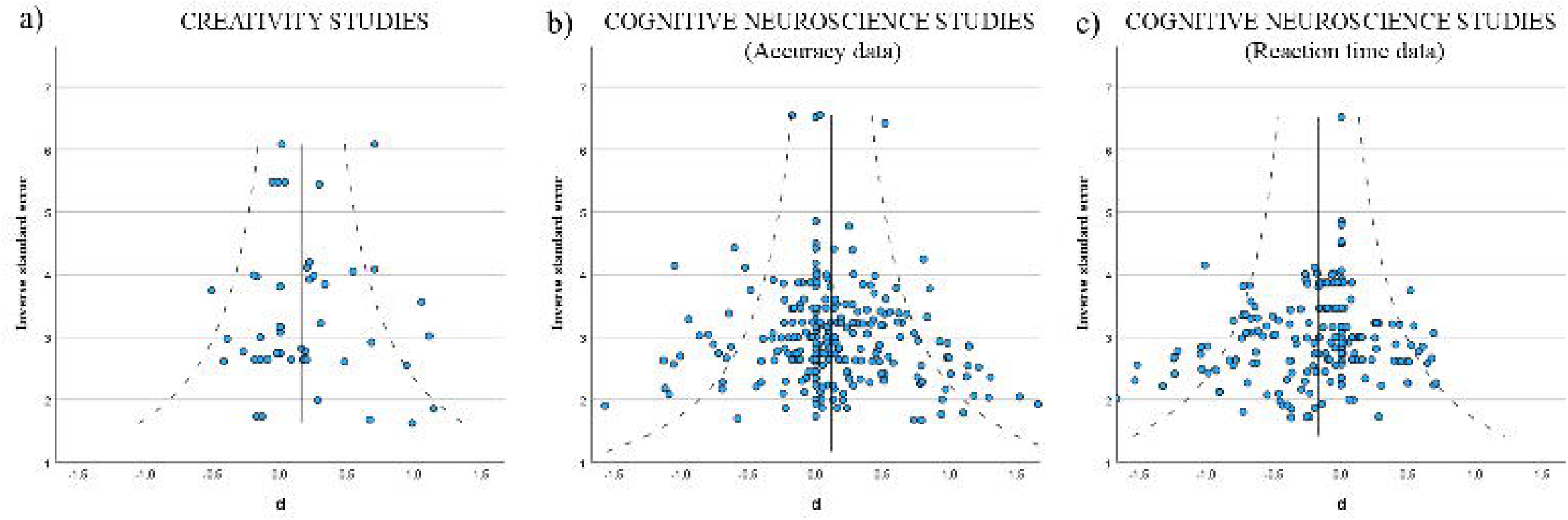

## Notes

https://github.com/melodymychan/CreativeThoughtTDCS.git

### Competing Interest Statement

The authors have declared no competing interest.

## References

1. Abraham, A. (2019). The neuropsychology of creativity. Current opinion in behavioral sciences, 27, 71–76.

2. Acharya, J. N., Hani, A. J., Cheek, J., Thirumala, P., & Tsuchida, T. N. (2016). American clinical neurophysiology society guideline 2: guidelines for standard electrode position nomenclature. The Neurodiagnostic Journal, 56(4), 245–252.

3. Aihara, T., Ogawa, T., Shimokawa, T., & Yamashita, O. (2017). Anodal transcranial direct current stimulation of the right anterior temporal lobe did not significantly affect verbal insight. Plos one, 12(9), e0184749.

4. Assem, M., Shashidhara, S., Glasser, M. F., & Duncan, J. (2024). Basis of executive functions in fine-grained architecture of cortical and subcortical human brain networks. Cerebral Cortex, 34(2), bhad537.

5. Baker, J. M., Rorden, C., & Fridriksson, J. (2010). Using transcranial direct-current stimulation to treat stroke patients with aphasia. Stroke, 41(6), 1229–1236.

6. Bartel, G., Marko, M., Rameses, I., Lamm, C., & Riečanský, I. (2020). Left prefrontal cortex supports the recognition of meaningful patterns in ambiguous stimuli. Frontiers in Neuroscience, 14, 152.

7. Beaty, R. E., & Kenett, Y. N. (2023). Associative thinking at the core of creativity. Trends in cognitive sciences.

8. Benedek, M., Beaty, R. E., Schacter, D. L., & Kenett, Y. N. (2023). The role of memory in creative ideation. Nature Reviews Psychology, 2(4), 246–257.

9. Benedek, M., & Fink, A. (2019). Toward a neurocognitive framework of creative cognition: The role of memory, attention, and cognitive control. Current opinion in behavioral sciences, 27, 116–122.

10. Benedek, M., Jauk, E., Beaty, R. E., Fink, A., Koschutnig, K., & Neubauer, A. C. (2016). Brain mechanisms associated with internally directed attention and self-generated thought. Scientific reports, 6(1), 22959.

11. Bjekić, J., Čolić, M. V., Živanović, M., Milanović, S. D., & Filipović, S. R. (2019). Transcranial direct current stimulation (tDCS) over parietal cortex improves associative memory. Neurobiology of Learning and Memory, 157, 114–120.

12. Bona, S., & Silvanto, J. (2014). Accuracy and confidence of visual short-term memory do not go hand-in-hand: behavioral and neural dissociations. Plos one, 9(3), e90808.

13. Borenstein, M. (2022). Comprehensive meta_analysis software. Systematic reviews in health research: metalJanalysis in context, 535–548.

14. Borenstein, M., Hedges, L. V., Higgins, J. P., & Rothstein, H. R. (2021). Introduction to meta-analysis. John Wiley & Sons.

15. Bradley, C., Nydam, A. S., Dux, P. E., & Mattingley, J. B. (2022). State-dependent effects of neural stimulation on brain function and cognition. Nature reviews neuroscience, 23(8), 459–475.

16. Cattaneo, Z., Pisoni, A., & Papagno, C. (2011). Transcranial direct current stimulation over Broca’s region improves phonemic and semantic fluency in healthy individuals. Neuroscience, 183, 64–70.

17. Chan, M. M., Lambon Ralph, M., & Robinson, G. A. (2025). The neural basis of creative thought: An activation likelihood estimation meta-analysis involving over 17,000 participants. bioRxiv, 2025–02. 10.1101/2025.02.19.639155

18. Chan, M. M., Lambon Ralph, M., & Robinson, G. (2023). The neurocognitive cornerstones of creative thought. 10.31234/osf.io/mv2q7

19. Chan, M. M., Yau, S. S., & Han, Y. M. (2021). The neurobiology of prefrontal transcranial direct current stimulation (tDCS) in promoting brain plasticity: A systematic review and meta-analyses of human and rodent studies. Neuroscience & biobehavioral reviews, 125, 392–416.

20. Chen, Q., Ding, K., Chen, Z., Yang, Y., Yu, R., Kenett, Y. N., & Qiu, J. (2024). A meta-analysis of the effects of transcranial direct current stimulation on creative thinking. *Psychology of Aesthetics*, Creativity, and the Arts.

21. Chi, R. P., & Snyder, A. W. (2012). Brain stimulation enables the solution of an inherently difficult problem. Neuroscience letters, 515(2), 121–124.

22. Christoff, K., Irving, Z. C., Fox, K. C., Spreng, R. N., & Andrews-Hanna, J. R. (2016). Mind-wandering as spontaneous thought: a dynamic framework. Nature reviews neuroscience, 17(11), 718–731.

23. Chrysikou, E. G. (2019). Creativity in and out of (cognitive) control. Current opinion in behavioral sciences, 27, 94–99.

24. Corbett, F., Jefferies, E., Ehsan, S., & Lambon Ralph, M. A. (2009). Different impairments of semantic cognition in semantic dementia and semantic aphasia: evidence from the non-verbal domain. Brain, 132(9), 2593–2608.

25. Cropley, A. (2006). In praise of convergent thinking. Creativity research journal, 18(3), 391–404.

26. Dane, E., & Pratt, M. G. (2007). Exploring intuition and its role in managerial decision making. Academy of management review, 32(1), 33–54.

27. Datta, A., Bansal, V., Diaz, J., Patel, J., Reato, D., & Bikson, M. (2009). Gyri-precise head model of transcranial direct current stimulation: improved spatial focality using a ring electrode versus conventional rectangular pad. Brain stimulation, 2(4), 201–207. e201.

28. Daviddi, S., Pedale, T., Jacques, P. L. S., Schacter, D. L., & Santangelo, V. (2023). Common and distinct correlates of construction and elaboration of episodic-autobiographical memory: An ALE meta-analysis. Cortex.

29. de Boer, N. S., Schluter, R. S., Daams, J. G., van der Werf, Y. D., Goudriaan, A. E., & van Holst, R. J. (2021). The effect of non-invasive brain stimulation on executive functioning in healthy controls: a systematic review and meta-analysis. Neuroscience & biobehavioral reviews, 125, 122–147.

30. Domenech, P., Rheims, S., & Koechlin, E. (2020). Neural mechanisms resolving exploitation-exploration dilemmas in the medial prefrontal cortex. Science, 369(6507), eabb0184.

31. Ericsson, K. A., Prietula, M. J., & Cokely, E. T. (2007). The making of an expert. Harvard business review, 85(7/8), 114.

32. Evans, A. C., Janke, A. L., Collins, D. L., & Baillet, S. (2012). Brain templates and atlases. NeuroImage, 62(2), 911–922.

33. Faul, F., Erdfelder, E., Buchner, A., & Lang, A.-G. (2009). Statistical power analyses using G* Power 3.1: Tests for correlation and regression analyses. Behavior research methods, 41(4), 1149–1160.

34. Fertonani, A., Rosini, S., Cotelli, M., Rossini, P. M., & Miniussi, C. (2010). Naming facilitation induced by transcranial direct current stimulation. Behavioural brain research, 208(2), 311–318.

35. Flaherty, A. W. (2005). Frontotemporal and dopaminergic control of idea generation and creative drive. Journal of Comparative Neurology, 493(1), 147–153.

36. Fox, N. A., Bakermans-Kranenburg, M. J., Yoo, K. H., Bowman, L. C., Cannon, E. N., Vanderwert, R. E., Ferrari, P. F., & Van IJzendoorn, M. H. (2016). Assessing human mirror activity with EEG mu rhythm: A meta-analysis. Psychological bulletin, 142(3), 291.

37. Fridriksson, J., Richardson, J. D., Baker, J. M., & Rorden, C. (2011). Transcranial direct current stimulation improves naming reaction time in fluent aphasia: a double-blind, sham-controlled study. Stroke, 42(3), 819–821.

38. Galli, G., Vadillo, M. A., Sirota, M., Feurra, M., & Medvedeva, A. (2019). A systematic review and meta-analysis of the effects of transcranial direct current stimulation (tDCS) on episodic memory. Brain stimulation, 12(2), 231–241.

39. Gonen-Yaacovi, G., de Souza, L. C., Levy, R., Urbanski, M., Josse, G., & Volle, E. (2013). Rostral and caudal prefrontal contribution to creativity: a meta-analysis of functional imaging data. Frontiers in human neuroscience, 7, 465.

40. Gruber, H. E., & Wallace, D. B. (1999). The case study method and evolving systems approach for understanding unique creative people at work.

41. Guo, J., Luo, J., An, Y., & Xia, T. (2023). tDCS Anodal Stimulation of the Right Dorsolateral Prefrontal Cortex Improves Creative Performance in Real-World Problem Solving. Brain Sciences, 13(3), 449.

42. Haddaway, N. R., Collins, A. M., Coughlin, D., & Kirk, S. (2015). The role of Google Scholar in evidence reviews and its applicability to grey literature searching. Plos one, 10(9), e0138237.

43. Han, L. T., Cohen, M. S., He, L. K., Green, L. M., Knowlton, B. J., Castel, A. D., & Rissman, J. (2023). Establishing a causal role for left ventrolateral prefrontal cortex in value-directed memory encoding with high-definition transcranial direct current stimulation. Neuropsychologia, 181, 108489.

44. Heo, J., Yi, K., Hong, J., & Kim, C. (2023). The role of the prefrontal cortex in semantic control for selecting weakly associated meanings in creative idea generation. Neuroscience letters, 802, 137177.

45. Higgins, J. P., Thompson, S. G., Deeks, J. J., & Altman, D. G. (2003). Measuring inconsistency in meta-analyses. bmj, 327(7414), 557–560.

46. Hodgson, V. J., Lambon Ralph, M. A., & Jackson, R. L. (2023). The cross-domain functional organization of posterior lateral temporal cortex: insights from ALE meta-analyses of 7 cognitive domains spanning 12,000 participants. Cerebral Cortex, 33(8), 4990–5006.

47. Humphreys, G. F., & Lambon Ralph, M. A. (2015). Fusion and fission of cognitive functions in the human parietal cortex. Cerebral Cortex, 25(10), 3547–3560.

48. Humphreys, G. F., & Lambon Ralph, M. A. (2017). Mapping domain-selective and counterpointed domain-general higher cognitive functions in the lateral parietal cortex: evidence from fMRI comparisons of difficulty-varying semantic versus visuo-spatial tasks, and functional connectivity analyses. Cerebral Cortex, 27(8), 4199–4212.

49. Humphreys, G. F., & Ralph, M. A. L. (2023). Mapping the task-general and task-specific neural correlates of speech production: meta-analysis and fMRI direct comparisons of category fluency and picture naming. bioRxiv, 2023–09.

50. Humphreys, G. F., Halai, A. D., Branzi, F. M., & Lambon Ralph, M. A. (2024). The left posterior angular gyrus is engaged by autobiographical recall not object-semantics, or event-semantics: Evidence from contrastive propositional speech production. Imaging Neuroscience, 2, 1–19.

51. Ivancovsky, T., Kurman, J., Morio, H., & Shamay-Tsoory, S. (2019). Transcranial direct current stimulation (tDCS) targeting the left inferior frontal gyrus: Effects on creativity across cultures. Social Neuroscience, 14(3), 277–285.

52. Jackson, R. L. (2021). The neural correlates of semantic control revisited. NeuroImage, 224, 117444.

53. Jacobson, L., Koslowsky, M., & Lavidor, M. (2012). tDCS polarity effects in motor and cognitive domains: a meta-analytical review. Experimental brain research, 216, 1–10.

54. Joyal, M., & Fecteau, S. (2016). Transcranial direct current stimulation effects on semantic processing in healthy individuals. Brain stimulation, 9(5), 682–691.

55. Jung, J., Ralph, M. A. L., & Jackson, R. L. (2022). Subregions of DLPFC display graded yet distinct structural and functional connectivity. Journal of Neuroscience, 42(15), 3241–3252.

56. Kuang, C., Chen, J., Chen, J., Shi, Y., Huang, H., Jiao, B., Lin, Q., Rao, Y., Liu, W., & Zhu, Y. (2022). Uncovering neural distinctions and commodities between two creativity subsets: A meta_analysis of fMRI studies in divergent thinking and insight using activation likelihood estimation. Human Brain Mapping, 43(16), 4864–4885.

57. Lambon Ralph, M. A., Jefferies, E., Patterson, K., & Rogers, T. T. (2017). The neural and computational bases of semantic cognition. Nature reviews neuroscience, 18(1), 42–55.

58. Lebuda, I., & Benedek, M. (2023). A systematic framework of creative metacognition. Physics of Life Reviews.

59. Lenhard, W., & Lenhard, A. (2022). Computation of effect sizes. https://www.psychometrica.de/effect_size.html

60. Lucchiari, C., Sala, P. M., & Vanutelli, M. E. (2018). Promoting creativity through transcranial direct current stimulation (tDCS). A critical review. Frontiers in behavioral neuroscience, 12, 354522.

61. Luft, C. D. B., Zioga, I., Banissy, M. J., & Bhattacharya, J. (2017). Relaxing learned constraints through cathodal tDCS on the left dorsolateral prefrontal cortex. Scientific reports, 7(1), 2916.

62. Madore, K. P., Addis, D. R., & Schacter, D. L. (2015). Creativity and memory: Effects of an episodic-specificity induction on divergent thinking. Psychological science, 26(9), 1461–1468.

63. Mayseless, N., & Shamay-Tsoory, S. (2015). Enhancing verbal creativity: modulating creativity by altering the balance between right and left inferior frontal gyrus with tDCS. Neuroscience, 291, 167–176.

64. Mekern, V., Hommel, B., & Sjoerds, Z. (2019). Computational models of creativity: a review of single-process and multi-process recent approaches to demystify creative cognition. Current opinion in behavioral sciences, 27, 47–54.

65. Miniussi, C., Harris, J. A., & Ruzzoli, M. (2013). Modelling non-invasive brain stimulation in cognitive neuroscience. Neuroscience & biobehavioral reviews, 37(8), 1702–1712.

66. Narmashiri, A., & Akbari, F. (2023). The Effects of Transcranial Direct Current Stimulation (tDCS) on the Cognitive Functions: A Systematic Review and Meta-analysis. Neuropsychology review, 1–27.

67. O’Mara-Eves, A., Thomas, J., McNaught, J., Miwa, M., & Ananiadou, S. (2015). Using text mining for study identification in systematic reviews: a systematic review of current approaches. Systematic reviews, 4, 1–22.

68. Ostrowski, J., Svaldi, J., & Schroeder, P. A. (2022). More focal, less heterogeneous? Multi-level meta-analysis of cathodal high-definition transcranial direct current stimulation effects on language and cognition. Journal of Neural Transmission, 129(7), 861–878.

69. Page, M. J., McKenzie, J. E., Bossuyt, P. M., Boutron, I., Hoffmann, T. C., Mulrow, C. D., Shamseer, L., Tetzlaff, J. M., Akl, E. A., & Brennan, S. E. (2021). The PRISMA 2020 statement: an updated guideline for reporting systematic reviews. bmj, 372.

70. Pick, H., & Lavidor, M. (2019). Modulation of automatic and creative features of the Remote Associates Test by angular gyrus stimulation. Neuropsychologia, 129, 348–356.

71. Polanía, R., Nitsche, M. A., & Ruff, C. C. (2018). Studying and modifying brain function with non-invasive brain stimulation. Nature neuroscience, 21(2), 174–187.

72. Poldrack, R. A., & Yarkoni, T. (2016). From brain maps to cognitive ontologies: informatics and the search for mental structure. Annual review of psychology, 67(1), 587–612.

73. Robinson, G., Shallice, T., Bozzali, M., & Cipolotti, L. (2012). The differing roles of the frontal cortex in fluency tests. Brain, 135(7), 2202–2214.

74. Ruggiero, F., Lavazza, A., Vergari, M., Priori, A., & Ferrucci, R. (2018). Transcranial direct current stimulation of the left temporal lobe modulates insight. Creativity research journal, 30(2), 143–151.

75. Schroeder, P. A., Schwippel, T., Wolz, I., & Svaldi, J. (2020). Meta-analysis of the effects of transcranial direct current stimulation on inhibitory control. Brain stimulation, 13(5), 1159–1167.

76. Song, S., Zilverstand, A., Gui, W., Li, H.-j., & Zhou, X. (2019). Effects of single-session versus multi-session non-invasive brain stimulation on craving and consumption in individuals with drug addiction, eating disorders or obesity: A meta-analysis. Brain stimulation, 12(3), 606–618.

77. Sternberg, R. J. (1988). A three-facet model of creativity.

78. Strobach, T., & Antonenko, D. (2017). tDCS-induced effects on executive functioning and their cognitive mechanisms: a review. Journal of Cognitive Enhancement, 1, 49–64.

79. Stuss, D. T. (2011). Functions of the frontal lobes: relation to executive functions. Journal of the international neuropsychological Society, 17(5), 759–765.

80. Thair, H., Holloway, A. L., Newport, R., & Smith, A. D. (2017). Transcranial direct current stimulation (tDCS): a beginner’s guide for design and implementation. Frontiers in Neuroscience, 11, 641.

81. Thompson, H. E., Henshall, L., & Jefferies, E. (2016). The role of the right hemisphere in semantic control: A case-series comparison of right and left hemisphere stroke. Neuropsychologia, 85, 44–61.

82. Tremblay, S., Lepage, J.-F., Latulipe-Loiselle, A., Fregni, F., Pascual-Leone, A., & Théoret, H. (2014). The uncertain outcome of prefrontal tDCS. Brain stimulation, 7(6), 773–783.

83. Uanhoro, J. O. (2017). Effect size calculators. https://effect-size-calculator.herokuapp.com/

84. Vatansever, D., Smallwood, J., & Jefferies, E. (2021). Varying demands for cognitive control reveals shared neural processes supporting semantic and episodic memory retrieval. Nature communications, 12(1), 2134.

85. Wang, H., He, W., Wu, J., Zhang, J., Jin, Z., & Li, L. (2019). A coordinate-based meta-analysis of the n-back working memory paradigm using activation likelihood estimation. Brain and cognition, 132, 1–12.

86. Westwood, S. J., & Romani, C. (2017). Transcranial direct current stimulation (tDCS) modulation of picture naming and word reading: A meta-analysis of single session tDCS applied to healthy participants. Neuropsychologia, 104, 234–249.

87. Wilson, D. B. (2023). Practical meta-analysis effect size calculator (Version 2023.11.27).

