## Supplementary material for "The cognitive and neural bases of creative thought: a cross-domain meta-analysis of transcranial direct current stimulation studies": Figure S1

**Supplementary figures**

**Figures S1-S7**: Forest plots for frontal montages

**Figures S8-S13**: Forest plots for parietal montages

**Figures S14-S18**: Forest plots for temporal montages

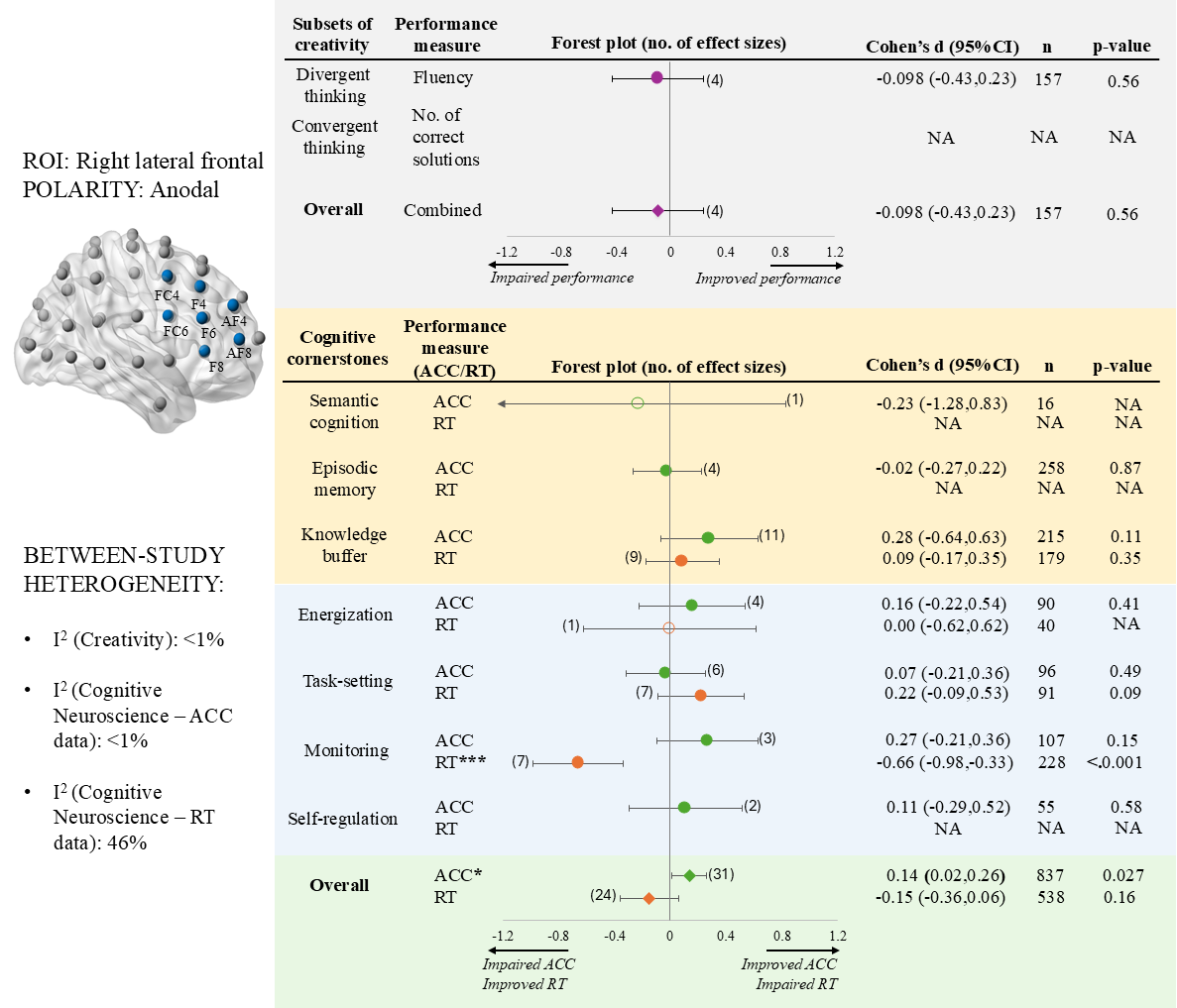

**Figure S1:** A forest plot summarising the right lateral frontal anodal tDCS effect on creativity and cognitive cornerstone components.

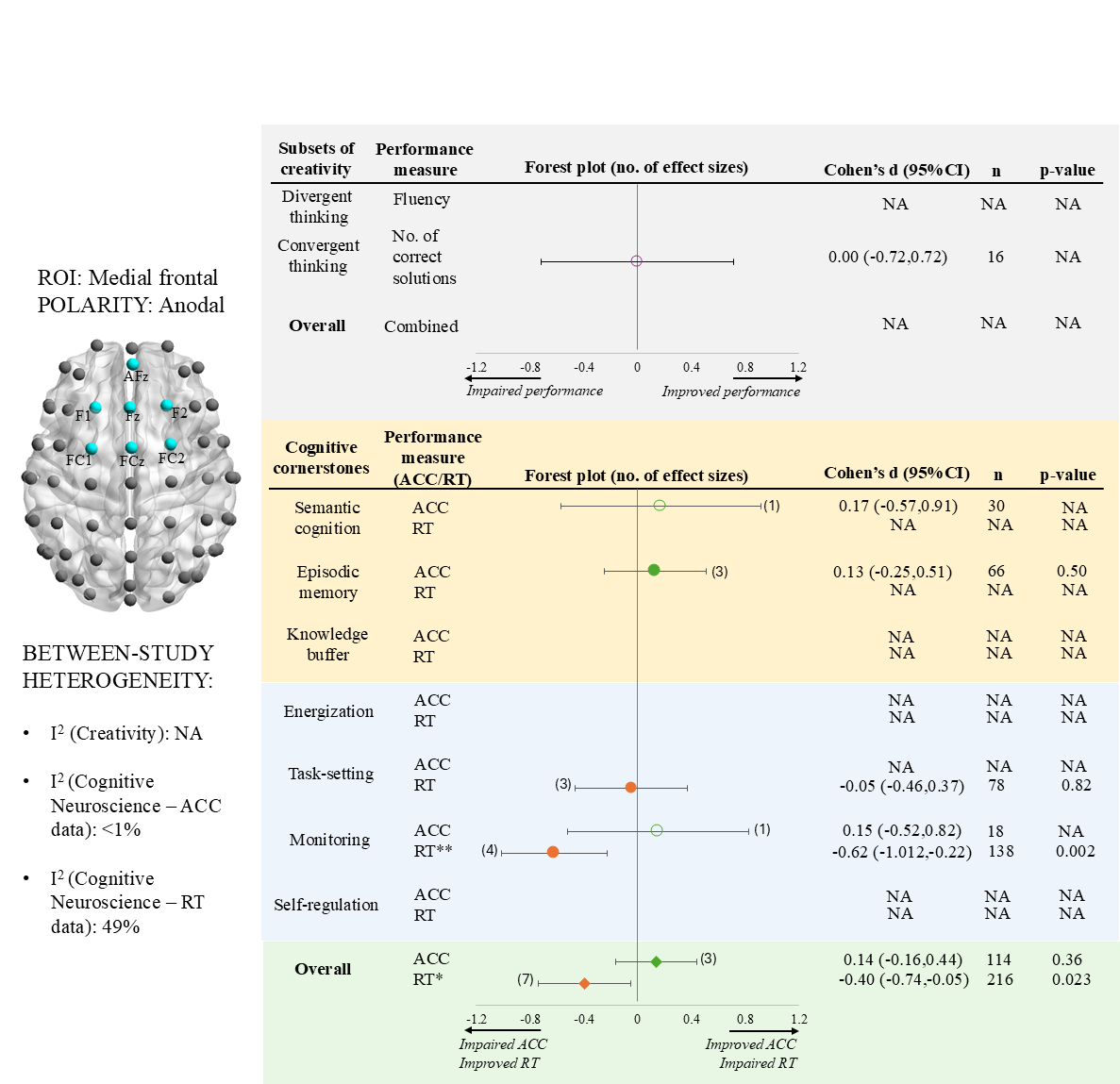
 **Figure S2:** A forest plot summarising the medial frontal anodal tDCS effect on creativity and cognitive cornerstone components.

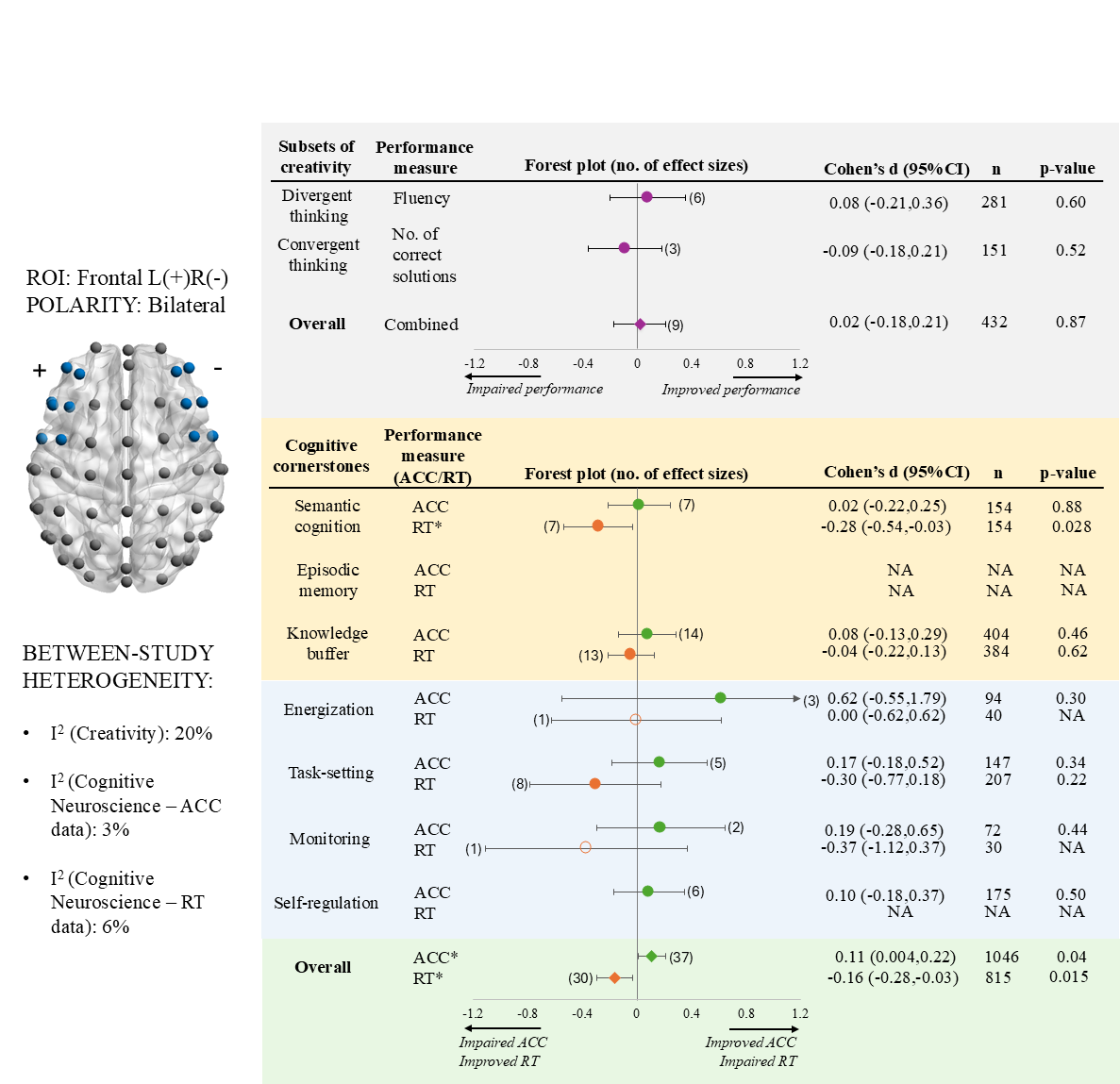

**Figure S3:** A forest plot summarising the left-frontal-anodal right-frontal-cathodal (frontal L+R-) tDCS effect on creativity and cognitive cornerstone components.

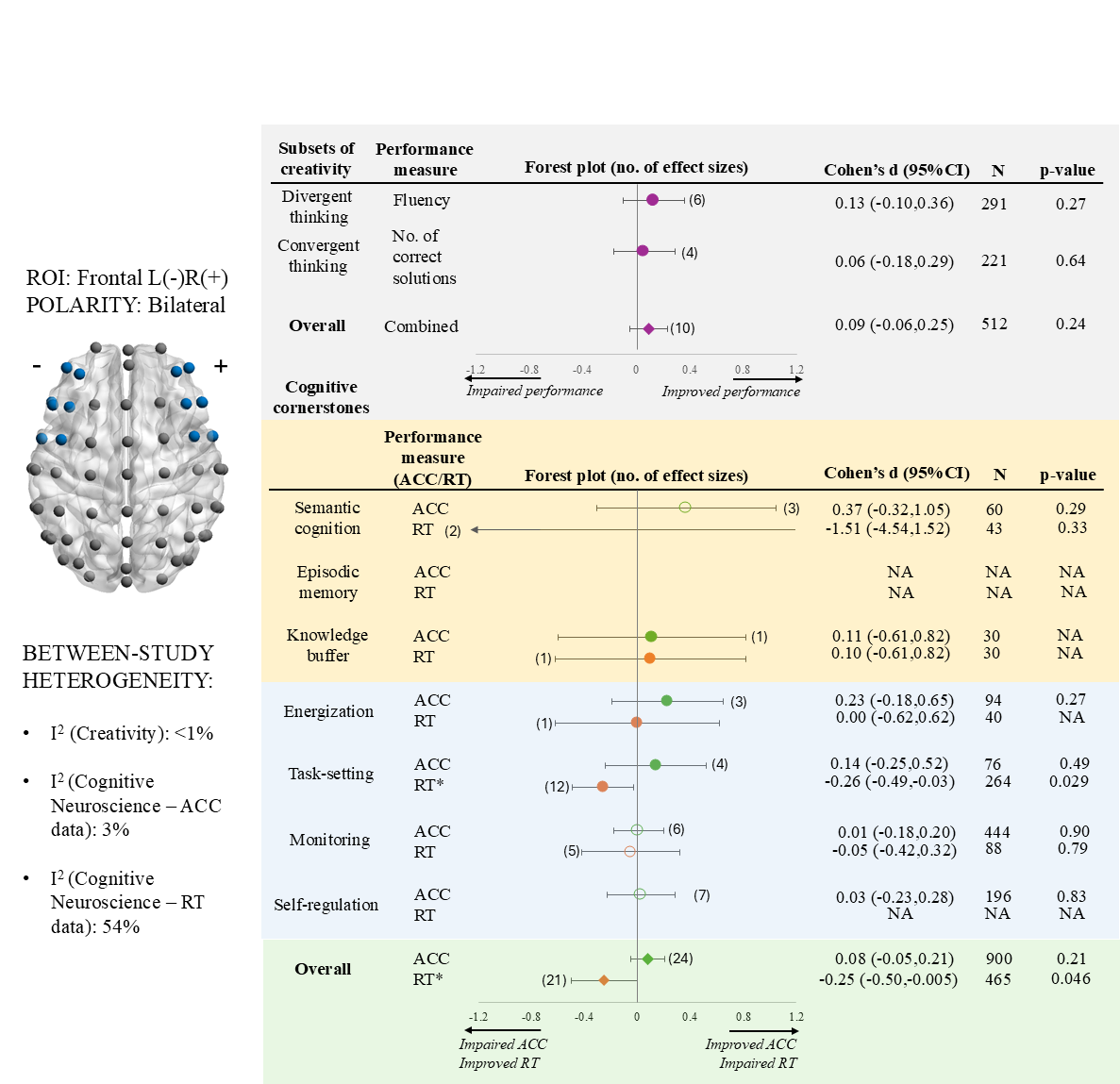

**Figure S4:** A forest plot summarising the left-frontal-cathodal right-frontal-anodal (frontal L-R+) tDCS effect on creativity and cognitive cornerstone components.

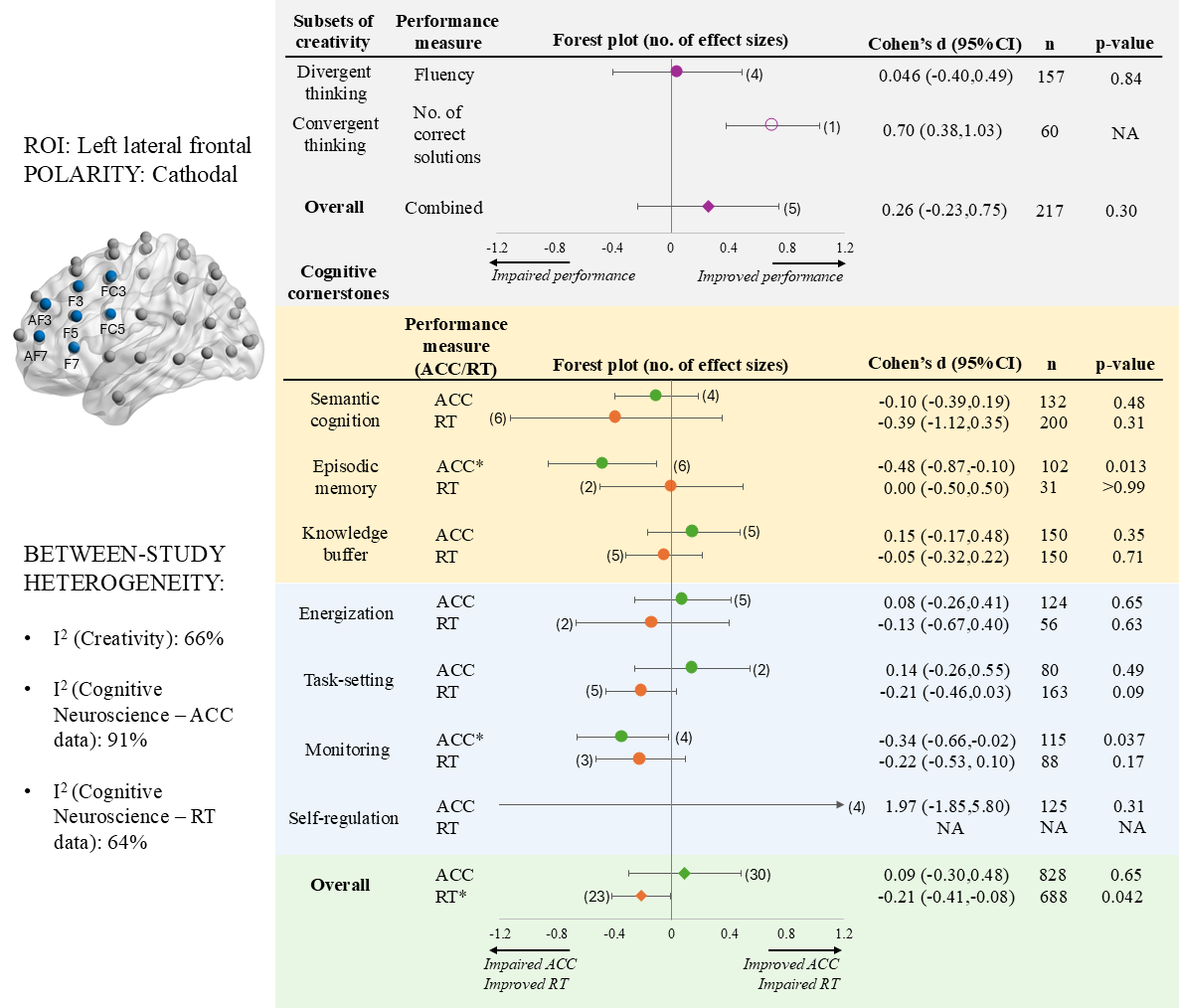

**Figure S5:** A forest plot summarising the left lateral frontal cathodal tDCS effect on creativity and cognitive cornerstone components.

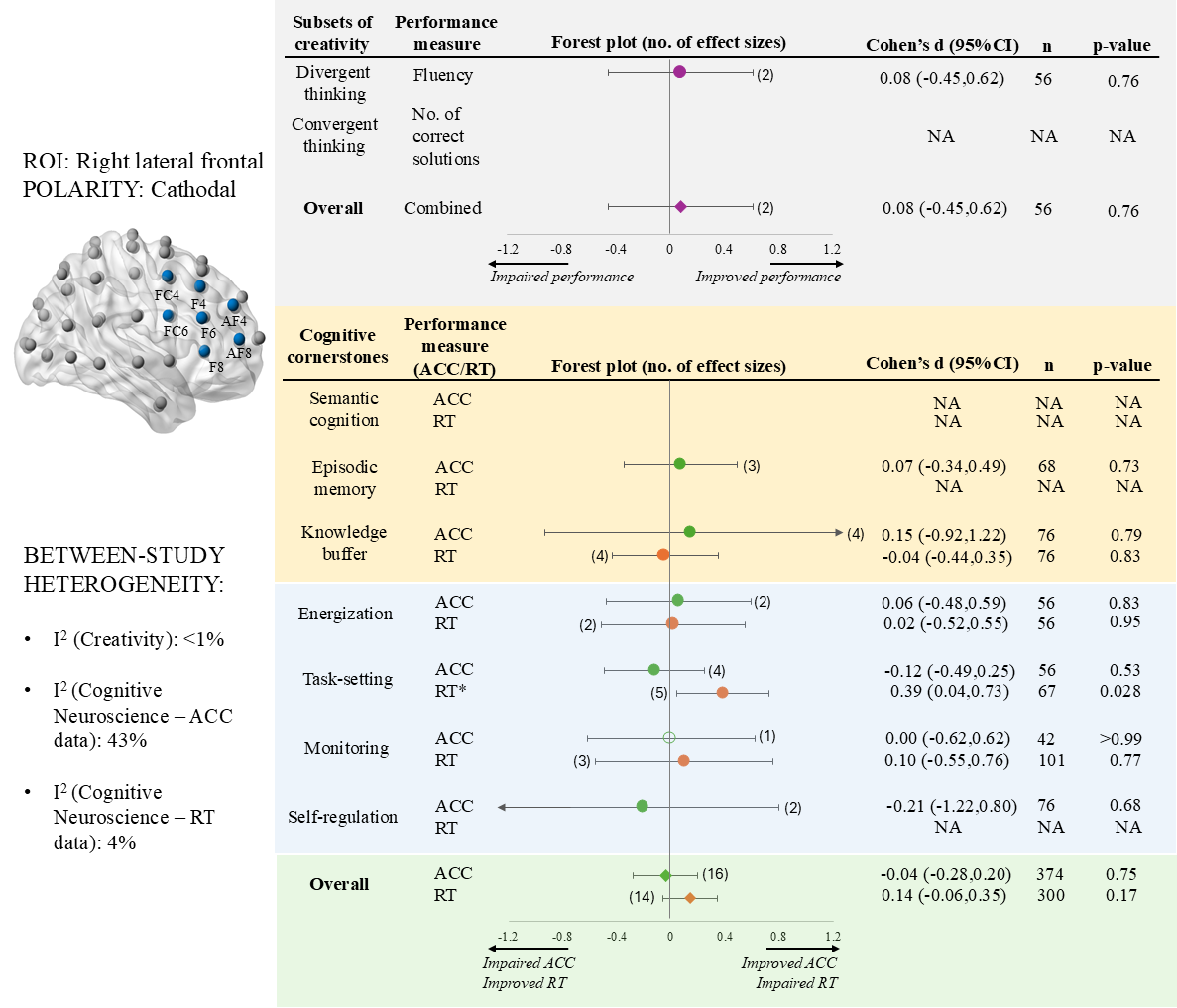

**Figure S6:** A forest plot summarising the right lateral frontal cathodal tDCS effect on creativity and cognitive cornerstone components.

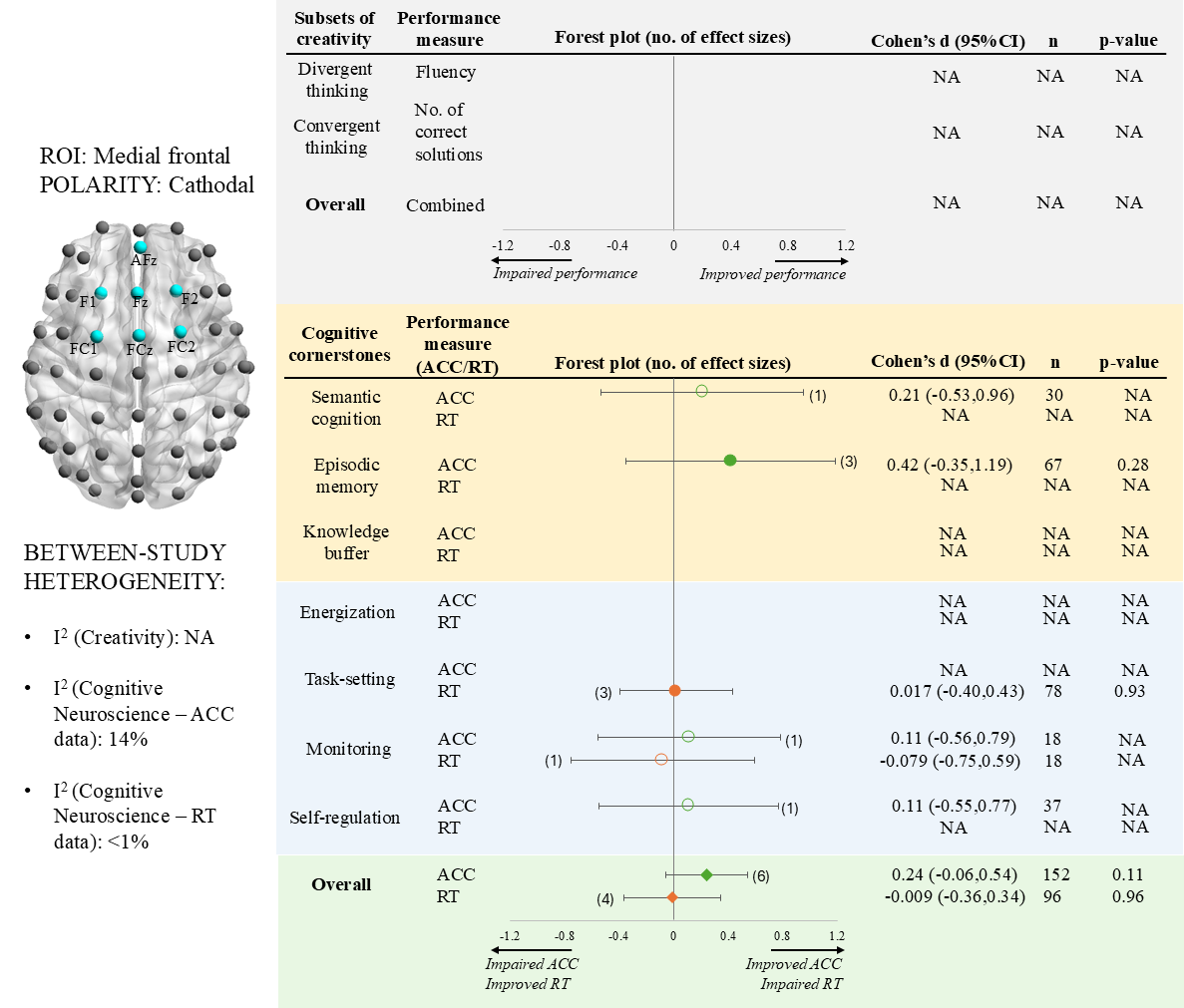

**Figure S7:** A forest plot summarising the medial frontal cathodal tDCS effect on creativity and cognitive cornerstone components.

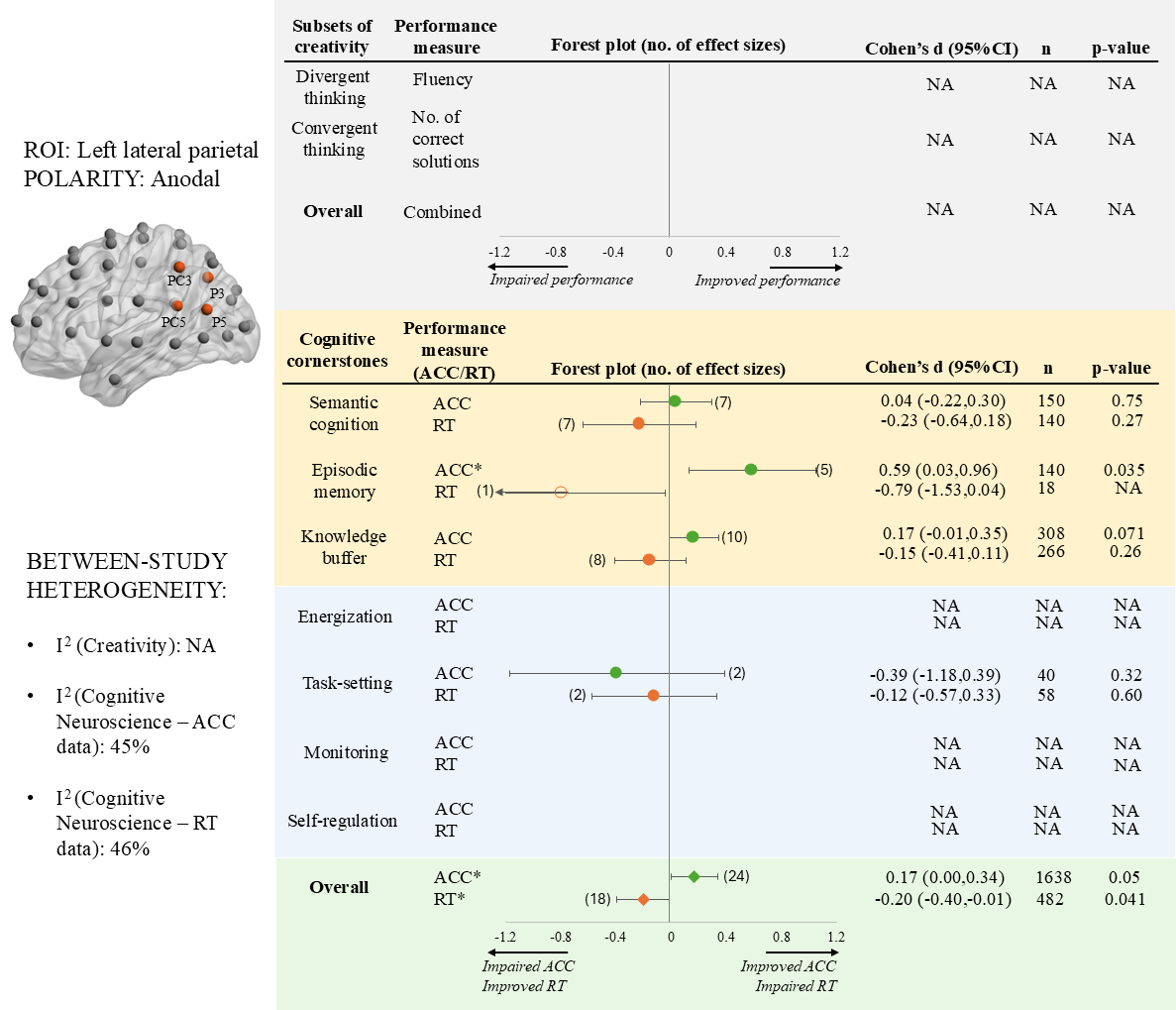

**Figure S8:** A forest plot summarising the left lateral parietal anodal tDCS effect on creativity and cognitive cornerstone components.

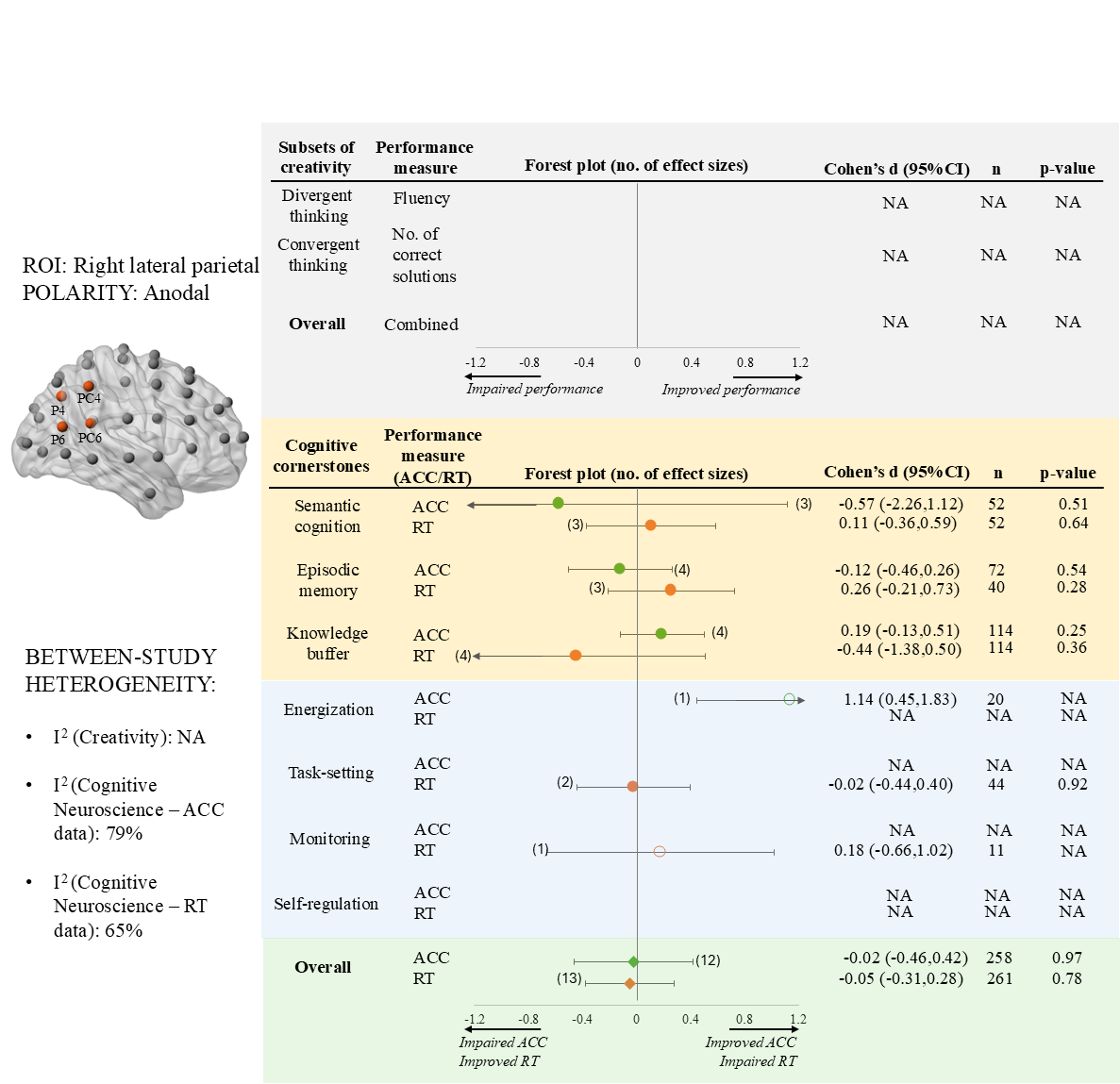

**Figure S9:** A forest plot summarising the right lateral parietal anodal tDCS effect on creativity and cognitive cornerstone components.

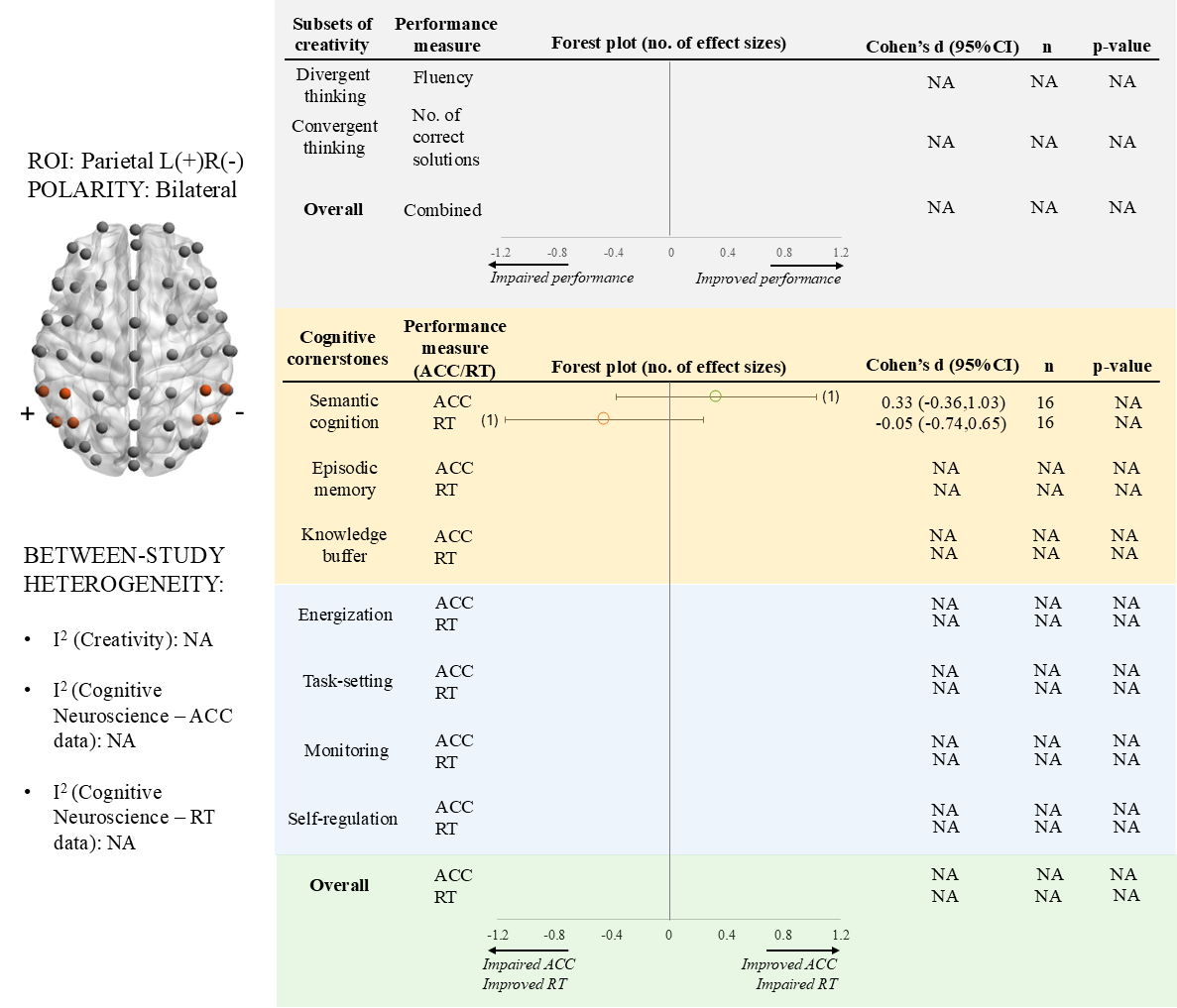

**Figure S10:** A forest plot summarising the left-parietal-anodal right-parietal-cathodal (parietal L+R-) tDCS effect on creativity and cognitive cornerstone components.

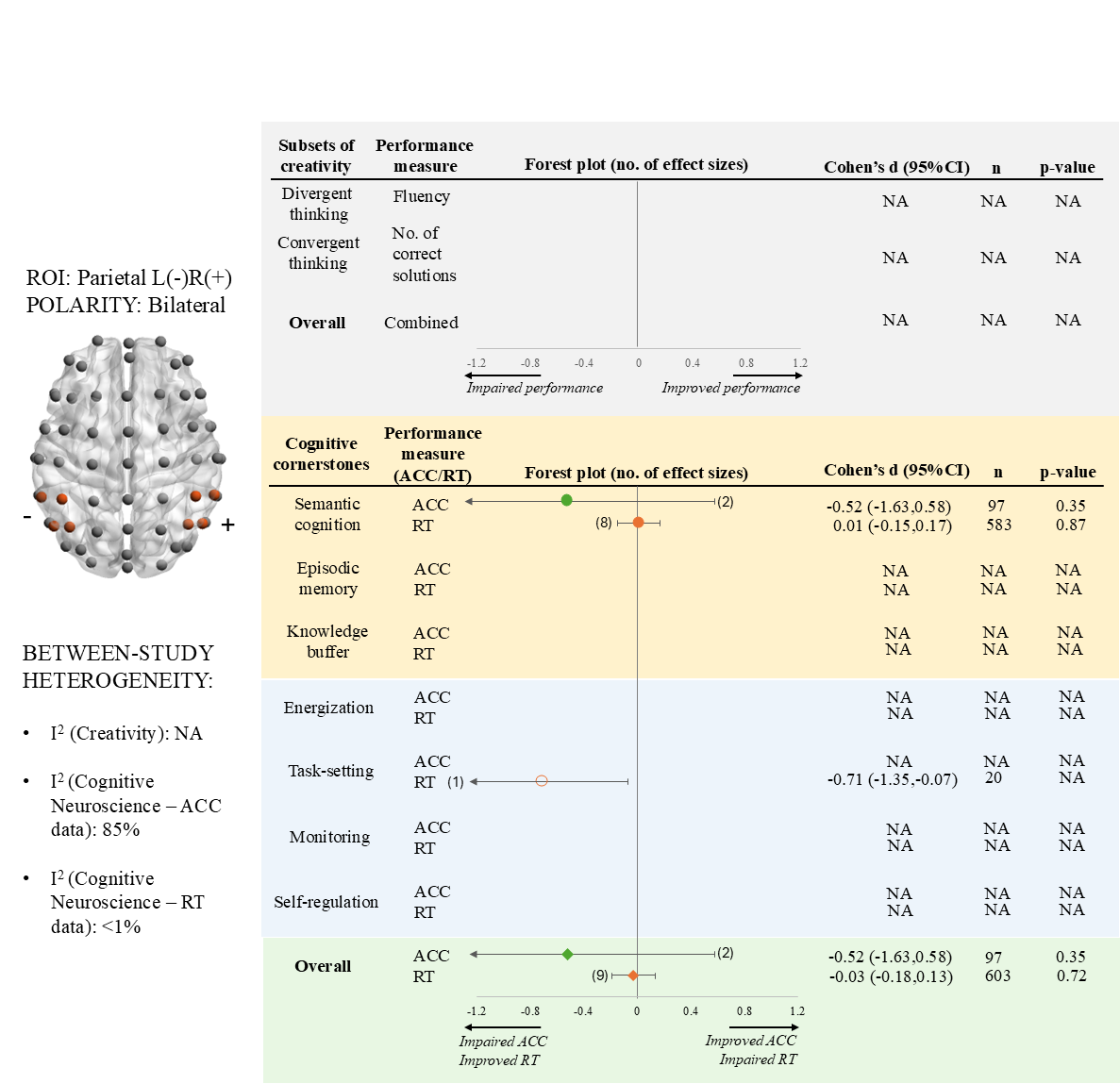

**Figure S11:** A forest plot summarising the left-parietal-cathodal right-parietal-anodal (parietal L-R+) tDCS effect on creativity and cognitive cornerstone components.

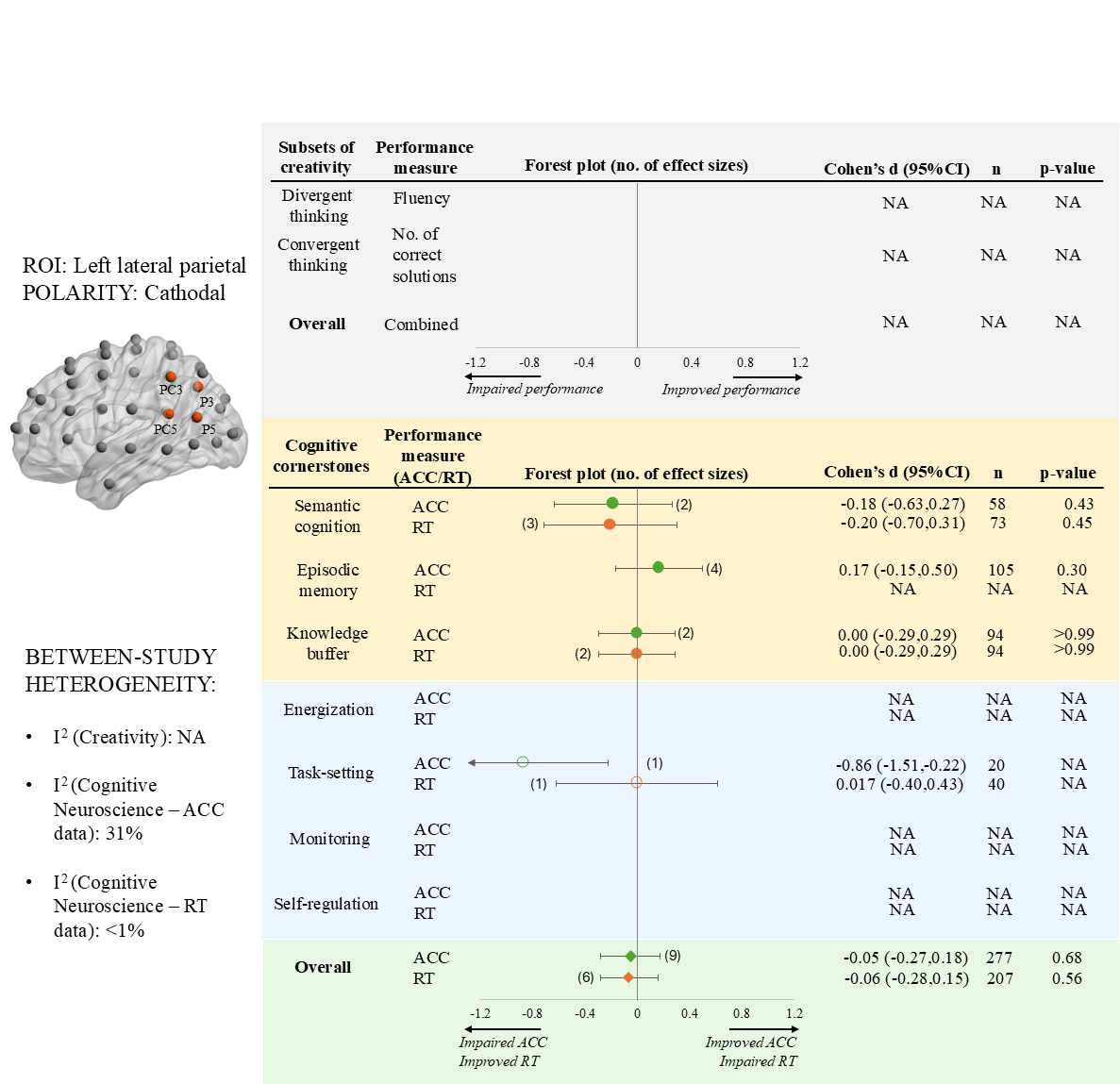

**Figure S12:** A forest plot summarising the left lateral parietal cathodal tDCS effect on creativity and cognitive cornerstone components.

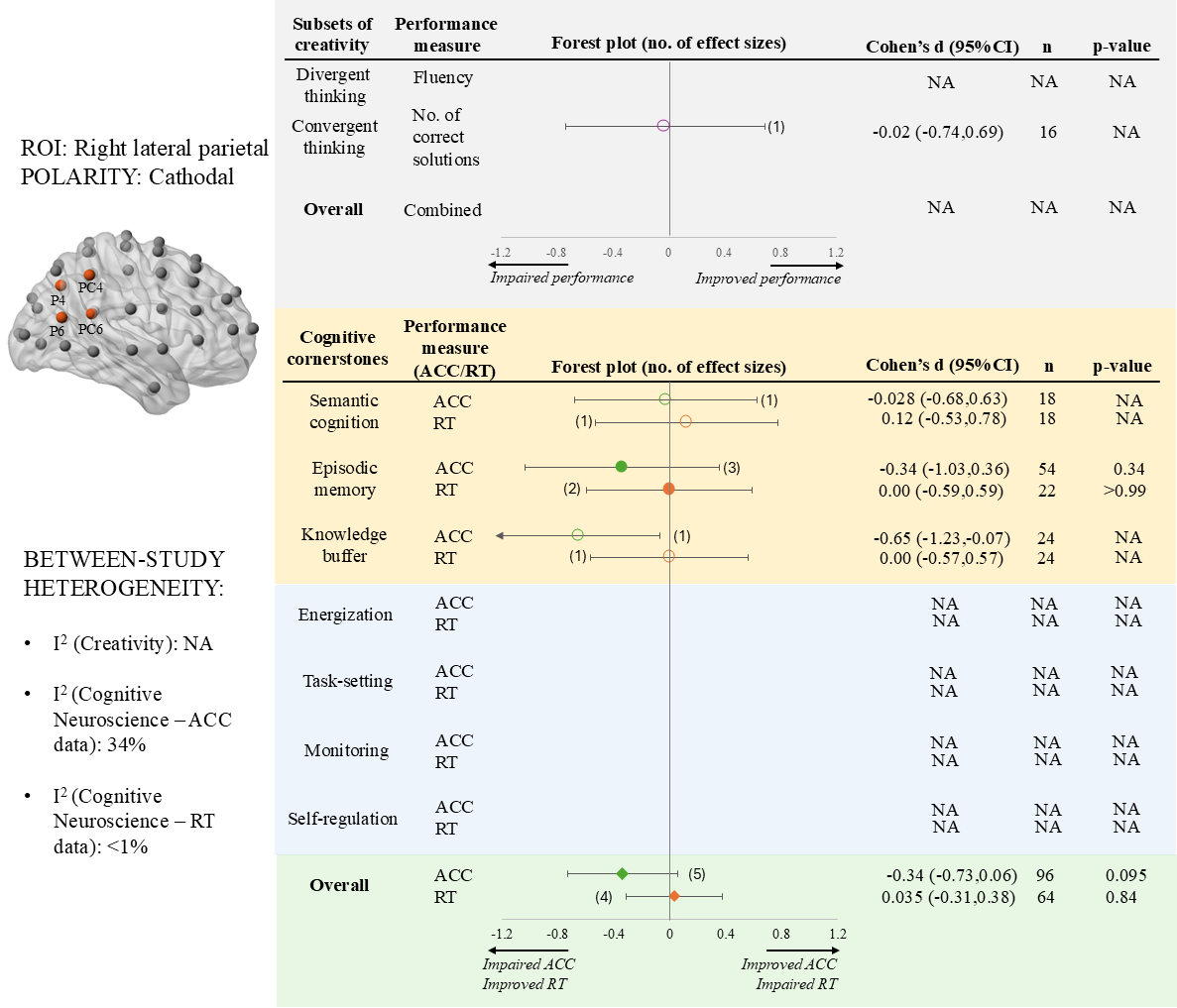

**Figure S13:** A forest plot summarising the right lateral parietal cathodal tDCS effect on creativity and cognitive cornerstone components.

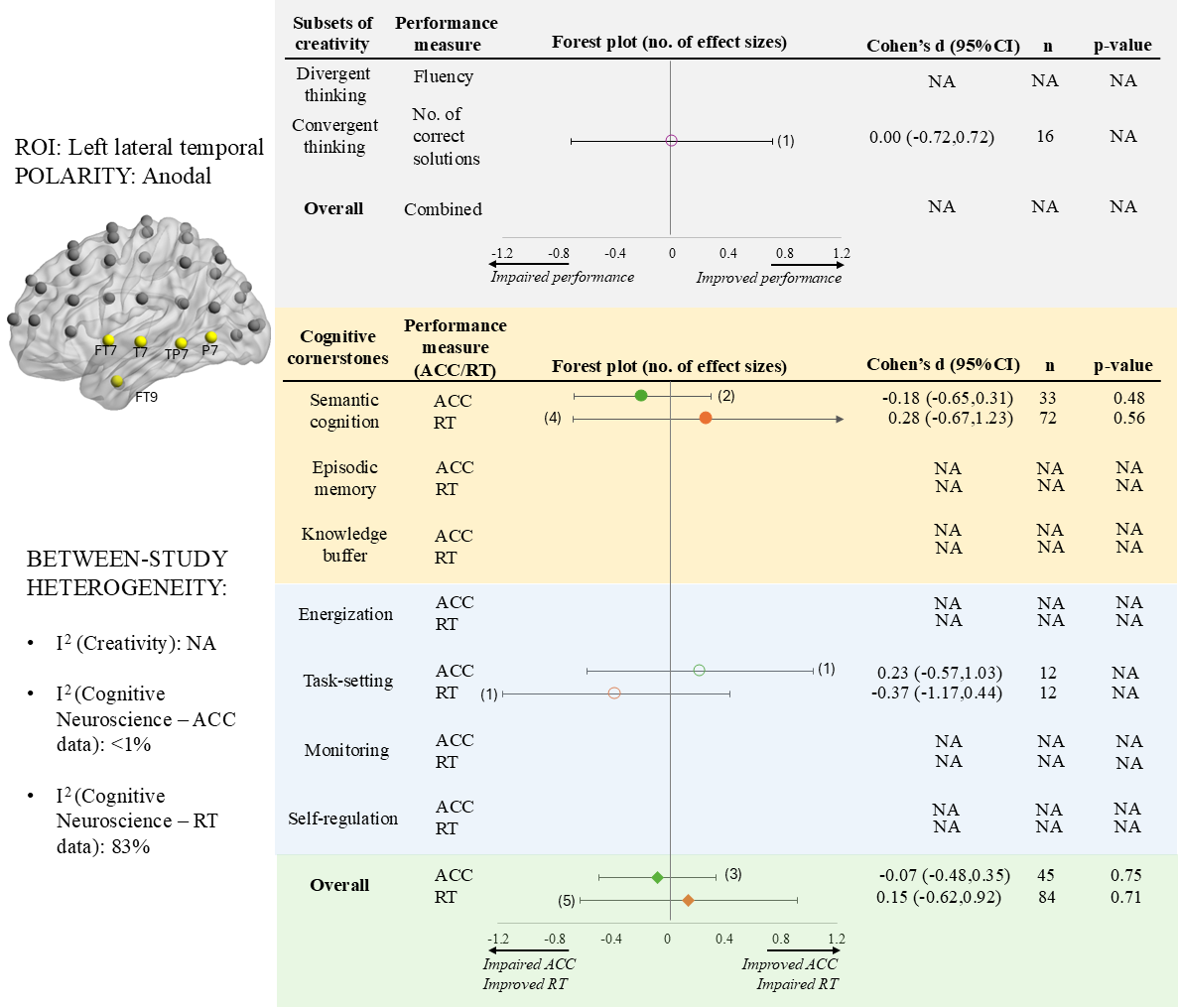

**Figure S14:** A forest plot summarising the left lateral temporal anodal tDCS effect on creativity and cognitive cornerstone components.

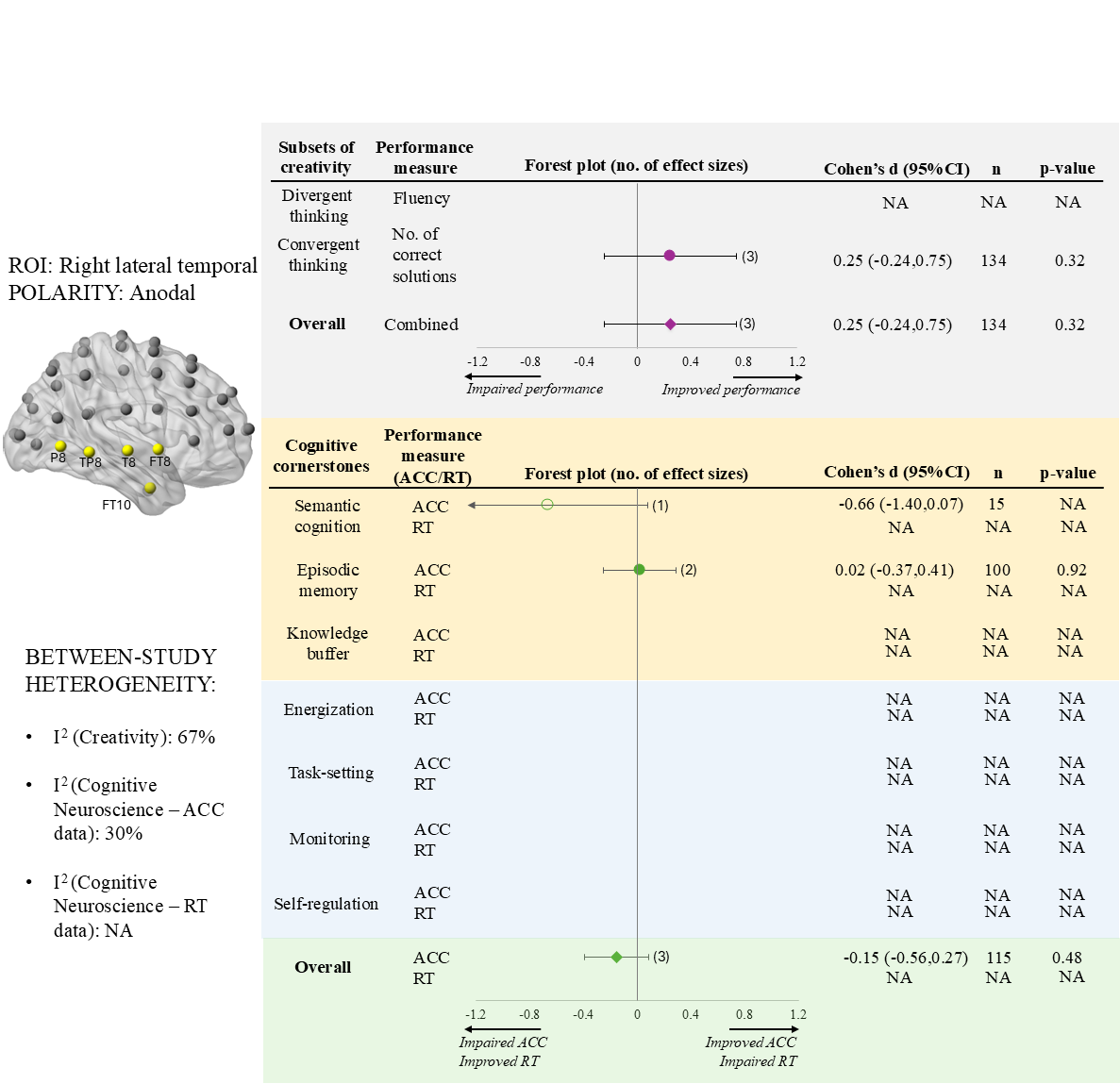

**Figure S15:** A forest plot summarising the right lateral temporal anodal tDCS effect on creativity and cognitive cornerstone components.

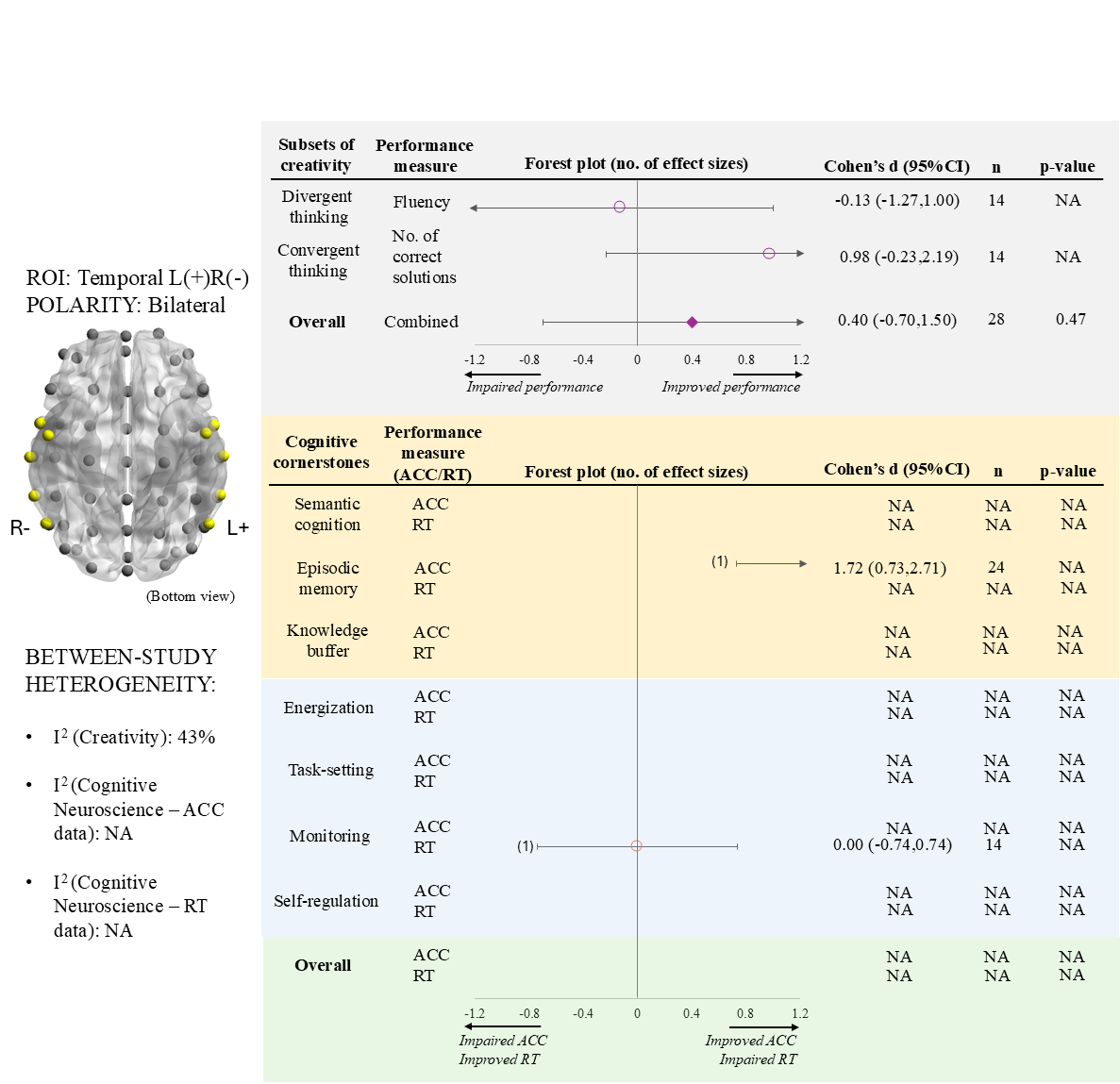

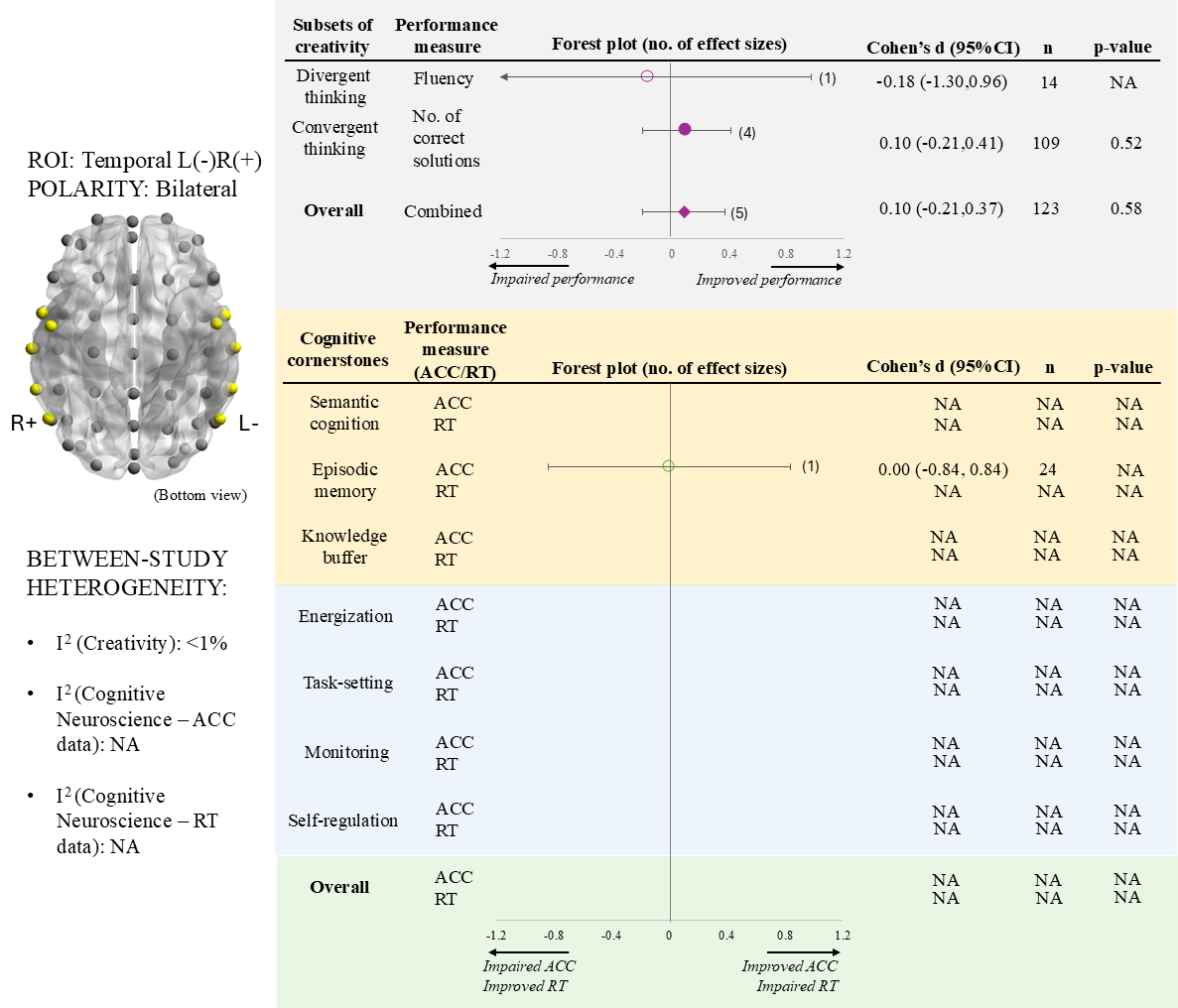

**Figure S17:** A forest plot summarising the left-temporal-cathodal right-temporal-anodal (temporal L-R+) tDCS effect on creativity and cognitive cornerstone components. Note that a bottom view of the brain template is used.

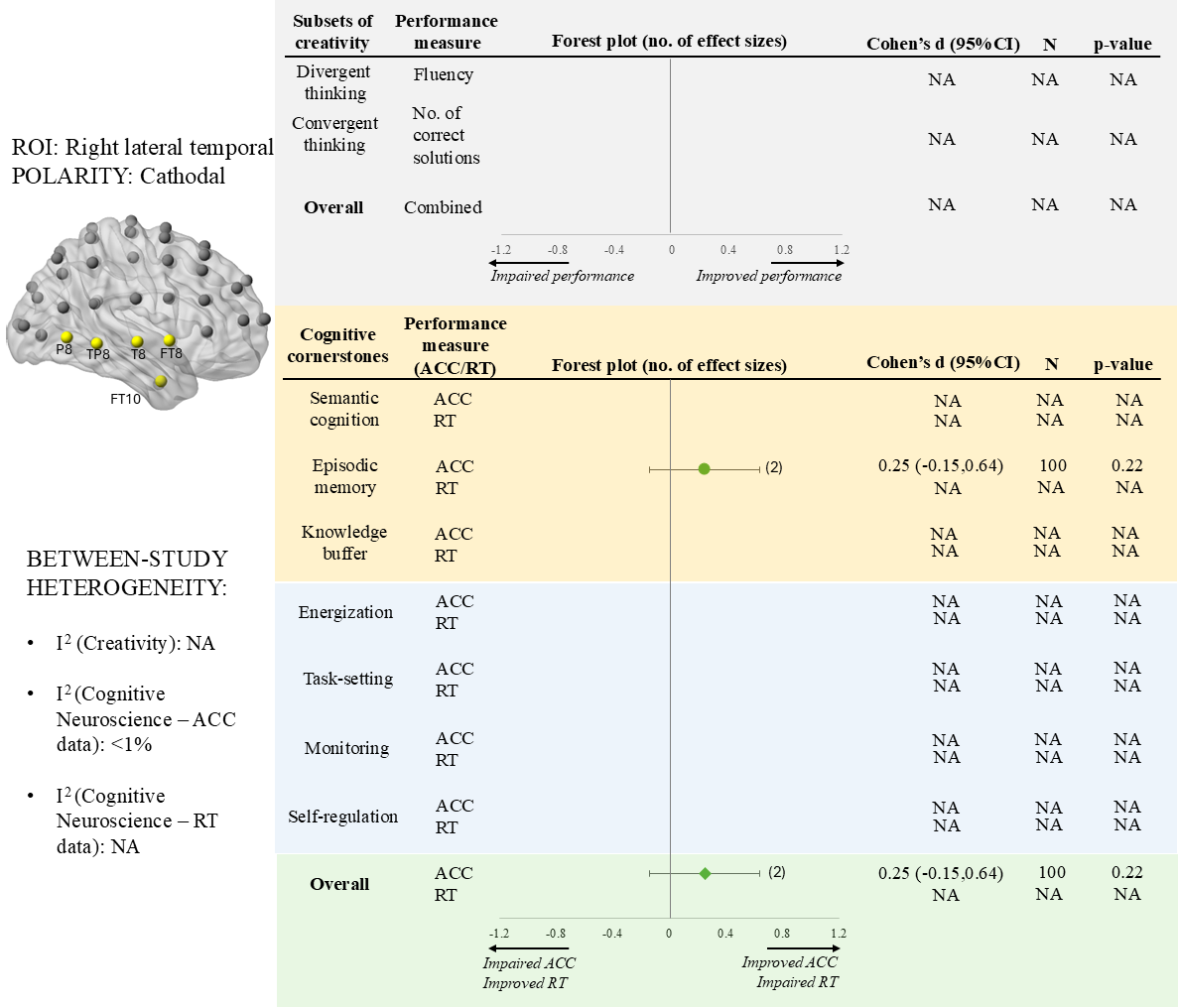

**Figure S18:** A forest plot summarising the right lateral temporal cathodal tDCS effect on creativity and cognitive cornerstone components.
