## Supplementary material for "The cognitive and neural bases of creative thought: a cross-domain meta-analysis of transcranial direct current stimulation studies": Table S1

| **Electronic database** | Embase | MEDLINE (EBSCOHost) | Web of Science Core Collection | APA PsycINFO |
| --- | --- | --- | --- | --- |
| **Search strategy** | creativ*:ti,ab,kw AND (process*:ti,ab,kw OR thought*:ti,ab,kw OR think*:ti,ab,kw OR executive:ti,ab,kw OR semantic:ti,ab,kw OR cognit*:ti,ab,kw OR control:ti,ab,kw) AND (brain:ti,ab,kw OR neural:ti,ab,kw OR neuroscien*:ti,ab,kw OR neurolog*:ti,ab,kw) AND article:it | (TI creativ* OR AB creativ*) AND ((TI (process* OR thought* OR think* OR executive OR semantic OR cognit* OR control) OR AB (process* OR thought* OR think* OR executive OR semantic OR cognit* OR control)) AND ((TI (brain OR neural OR neuroscien* OR neurolog*) OR AB (brain OR neural OR neuroscien* OR neurolog*) ) | creativ* (Topic) and process* OR thought* OR think* OR executive OR semantic OR cognit* OR control (Topic) and brain OR neural OR neuroscien* OR neurolog* (Topic) and Article (Document Types) | tiab,if(creativ*) AND tiab,if(process* OR thought* OR think* OR executive OR semantic OR cognit* OR control) AND tiab,if(brain OR neural OR neuroscien* OR neurolog*)  Additional limits - Record type: Conference Proceedings, Journal Article |

| **Reference (Year)** | **Study Design** | **Participant N** | **Mean Age in Years** | **Stimulation Modality** | **Anode** | **Cathode** | **Polarity of Stimulating Electrode** | **Online/Offline** | **Duration** | **Current Intensity, Electrode Size** | **Experimental Task Description** | **Semantic/**  **Less semantic** | **Visual/**  **Auditory** | **Verbal/**  **Nonverbal** |
| --- | --- | --- | --- | --- | --- | --- | --- | --- | --- | --- | --- | --- | --- | --- |
| ***Executive Functions*** | | | | | | | | | | | | | | |
| Boggio et al. (2008) [1] | Crossover, sham-controlled, double-blind trial | 14 | 23.4 | tDCS | T3 | T4 | Bilateral | Online assessment- performed during stimulation | 8 | 2 mA, 35 cm2 | **Facial expression GNG-** happy, sad, and neutral human faces presented, with happy or sad faces being targets to respond to | Semantic (picture) | Visual | Nonverbal |
| Cerruti & Schlaug (2009)- Exp. 1 [2] | Crossover, sham-controlled, double-blind trial | 18 | 25.5 | tDCS | F3 | Fp2 | Anodal | Online assessment- VFT performed before and during (near the end) of stimulation | 20 | 1 mA, 16.3 cm2 (active), 30 cm2 (reference) | **VFT (phonemic)** | less semantic | Visual | Verbal |
|  |  |  |  |  | Fp2 | F3 | Cathodal |  |  |  |  |  |  |  |
| Cerruti & Schlaug (2009)- Exp. 2 [2] | Crossover, sham-controlled, double-blind trial | 12 | 25.4 | tDCS | F3 | Fp2 | Anodal | Offline assessment- VFT performed before and after stimulation | 20 | 1 mA, 16.3 cm2 (active), 30 cm2 (reference) | **VFT (phonemic)** | less semantic | Visual | Verbal |
|  |  |  |  |  | F4 | Fp1 | Anodal |  |  |  |  |  |  |  |
| Fecteau et al. (2007a) [3] | Parallel, sham-controlled, double-blind trial | LA/RC: 12 RA/LC: 12 Sham: 12 | 20.3 | tDCS | F3 | F4 | Bilateral | Online assessment- performed during stimulation | <15 | 2 mA, 35 cm2 | **Gambling risk task-** choosing the colour of the box which the participant believes to have the 'winning token', where they are told that the token has equal probability of being hidden in any one of the boxes; rewarded with points when picking correct colour, but lose points when picking incorrect colour | Less semantic | Visual | Nonverbal |
|  |  |  |  |  | F4 | F3 | Bilateral |  |  |  |  |  |  |  |
| Ghanavati et al. (2018) [156] | Crossover, sham-controlled, double-blind trial | 20 | 27.55 | tDCS | P4 | Extracephalic | Anodal | Online assessment- performed during stimulation | 20 | 1.5mA, 35cm2 | **FPT-** produce as many different figures as possible by connect the dots within each rectangle | Less semantic | Visual | Nonverbal |
|  |  |  |  |  | F3 | Extracephalic |  |  |  |  |  |  |  |  |
| Iyer et al. (2005)- Exp. 1 [4] | Parallel, sham-controlled, participant-blind trial | Anodal: 10 Cathodal: 10 Sham: 10 | Anodal: 35.9 Cathodal: 36.4 Sham: 36.7 | tDCS | F3 | Fp2 | Anodal | Offline assessment- performed before and after stimulation | 20 | 1 mA, 25 cm2 | **VFT (phonemic)** | Less semantic | Visual | Verbal |
|  |  |  |  |  | Fp2 | F3 | Cathodal |  |  |  |  |  |  |  |
| Iyer et al. (2005)- Exp. 2 [4] | Parallel, sham-controlled, participant-blind trial | Anodal: 14 Cathodal: 14 Sham: 15 | Anodal: 36.4 Cathodal: 39.5 Sham: 36.7 | tDCS | F3 | Fp2 | Anodal | Online assessment- performed before and during stimulation | 20 | 1 mA, 25 cm2 | **VFT (phonemic)** | Less semantic | Visual | Verbal |
|  |  |  |  |  | Fp2 | F3 | Cathodal |  |  |  |  |  |  |  |
| Iyer et al. (2005)- Exp. 3 [4] | Parallel, sham-controlled, participant-blind trial | Anodal: 10 Cathodal: 10 Sham: 10 | Anodal: 38.7 Cathodal: 37.7 Sham: 39.5 | tDCS | F3 | Fp2 | Anodal | Online assessment- performed before and during stimulation | 20 | 2 mA, 25 cm2 | **VFT (phonemic)** | Less semantic | Visual | Verbal |
|  |  |  |  |  | Fp2 | F3 | Cathodal |  |  |  |  |  |  |  |
| Mameli et al. (2010) [5] | Crossover, sham-controlled, double-blind trial | 20 | 29.7 | tDCS | F3, F4 | Extracephalic | Anodal | Offline assessment- performed before (baseline) and after stimulation | 15 | 2 mA, 32 cm2 (active), 64 cm2 (reference) | **Visual attention task (Posner)-** exogenous cue version of Posner task (respond to target which appears at either one of two locations; valid cues, invalid cues, neutral cues) | Less semantic | Visual | Nonverbal |
| Stone & Tesche (2009) [6] | Crossover, sham-controlled trial | 20 | n.r. | tDCS | P3 | Extracephalic | Anodal | Online/offline assessment- performed before (baseline), during, and after (immediately [Post0] and 15 mins after end of tDCS [Post20]) stimulation | 20 | 2 mA, 25 cm2 (active), 25 cm2 or 157.5 cm2 (reference) | **Global/local task (compound letter task)-** respond to the global or local feature ('H' or 'S') of stimuli as instructed by a target cue, in which there are congruent and incongruent trials; after a sequence of compound letters, another target cue ('G' or 'L') is presented that indicates a response switch from local-to-global or global-to-local features | Less semantic | Visual | Nonverbal |
|  |  |  |  |  | Extracephalic | P3 | Cathodal |  |  |  |  |  |  |  |
| Fecteau et al. (2007b)- Exp. 1 [7] | Parallel, sham-controlled, double-blind trial | LA/RC: 10 RA/LC: 10 Sham: 10 | 21 | tDCS | F3 | F4 | Bilateral | Online assessment- BART performed during stimulation  Offline assessment- Stroop task performed before and after stimulation | <20 | 2 mA, 35 cm2 | **BART**  **SCWT** | Less semantic | Visual | Nonverbal |
|  |  |  |  |  | F4 | F3 | Bilateral |  |  |  |  |  |  |  |
| Fecteau et al. (2007b)- Exp. 2 [7] | Parallel, sham-controlled, double-blind trial | L: 6 R: 6 Sham: 10 | 21.7 | tDCS | F3 | Fp2 | Anodal | Online assessment- BART performed during stimulation  Offline assessment- Stroop task performed before and after stimulation | <20 | 2 mA, 35 cm2 | **BART** | Less semantic | Visual | Nonverbal |
|  |  |  |  |  | F4 | Fp1 | Anodal |  |  |  |  |  |  |  |
| Beeli et al. (2008) [8] | Crossover, sham-controlled trial | 35 | 24.9 | tDCS | FC3 | Extracephalic (ipsilateral) | Anodal | Online assessment- performed during stimulation | 5.5 | 1.5 mA, 35 cm2 | **GNG-** stimuli are lines in different directions | Less semantic | Visual | Nonverbal |
|  |  |  |  |  | Extracephalic (ipsilateral) | FC3 | Cathodal |  |  |  |  |  |  |  |
| Dockery et al. (2009) [9] | Crossover, sham-controlled, participant-blind trial | 24 | 24 | tDCS | F3 | Fp2 | Anodal | Online/offline assessment- performed during and after (short-term and long-term) stimulation | 15 | 1 mA, 35 cm2 | **TOL** | Less semantic | Visual | Nonverbal |
|  |  |  |  |  | Fp2 | F3 | Cathodal |  |  |  |  |  |  |  |
| Leite et al. (2011) [10] | Crossover, sham-controlled trial | 15 | n.r. | tDCS | F3 | Fp2 | Anodal | Offline assessment- performed after stimulation | 15 | 1 mA, 35 cm2 | **Set shifting task (cognitive)-** respond either to colour or shape having previously associated two colours and two shapes to the same response buttons; using two consecutive stimuli, the set either remains the same (e.g., colour-colour) or is different (e.g., colour-shape) | Less semantic | Visual | Nonverbal |
|  |  |  |  |  | Fp2 | F3 | Cathodal |  |  |  |  |  |  |  |
| Balconi & Vitaloni (2012) [11] | Crossover, sham-controlled trial | 34 | 23.13 | tDCS | Fp2 | F3 | Cathodal | Offline assessment- performed before (Phase 1) and after (Phase 3) stimulation (Phase 2 is stimulation) | 15 | 2 mA, 35 cm2 | **Semantic congruence processing task-** decide whether action sequences are presented with a congruous (using object properly) or incongruous (not using object properly) ending scene | Semantic (picture) | Visual | Nonverbal |
| Gladwin et al. (2012) [12] | Crossover, sham-controlled, participant-blind trial | 20 | 21.1 | tDCS | F3 | Fp2 | Anodal | Offline assessment- performed after stimulation | 10 | 1 mA, 35 cm2 | **IAT-** respond either congruently or incongruently to pre-existing evaluative associations (e.g., in congruent blocks, negative and insect words are mapped to one key, and positive and flower words to the other key; in incongruent blocks, negative and flower words are mapped to one key, and positive and insect words to the other key) | Semantic (word) | Visual | Nonverbal |
| Jeon & Han (2012) [13] | Parallel, sham-controlled trial | RA: 8 LA: 8 RS: 8 LS: 8 | RA: 35.13 LA: 39.50 RS: 37.88 LS: 36.50 | tDCS | F3 | Fp2 | Anodal | Offline assessment- performed before (baseline), immediately after, and two weeks after stimulation | 20 | 1 mA, 35 cm2 | **SCWT** | Semantic (word) | Visual | Verbal |
|  |  |  |  |  | F4 | Fp1 | Anodal |  |  |  |  |  |  |  |
| Leite et al. (2013) [14] | Crossover, sham-controlled trial | 16 | 24 | tDCS | F3 | F4 | Bilateral | Online assessment- performed during stimulation | Max. 30 | 2 mA, 35 cm2 | **Task-switching paradigms:**  **Letter/digit naming task-** when presented with a pair of targets (i.e., a letter and a digit), respond either to letter or digit depending on the visual colour cue (respond to the letter for green, respond to the digit for red), with there being switch and repeat trials  **Vowel-consonant/parity task-** identical to the above except that the response requires a judgement (in the green cue condition, respond whether the letter is a vowel or a consonant; in the red cue condition, respond whether the number is even or odd) | Less semantic | Visual | Nonverbal |
|  |  |  |  |  | F4 | F3 | Bilateral |  |  |  |  |  |  |  |
| Minati et al. (2012) [16] | Parallel, sham-controlled, participant-blind trial | LA/RC: 16 RA/LC: 15 Sham: 16 | LA/RC: 22.3 RA/LC: 20.9 Sham: 21.8 | tDCS | F3 | F4 | Bilateral | Online assessment- performed during stimulation | 20.5 ± 4.1 across Ps | 2 mA, 35 cm2 | **Gambling task-** monetary gamble defined by potential win, potential loss, and probability of winning; respond according to a four-level scale, i.e., "confident reject", "unsure reject", "unsure accept", and "confident accept" | Less semantic | Visual | Nonverbal |
|  |  |  |  |  | F4 | F3 | Bilateral |  |  |  |  |  |  |  |

| **Reference (Year)** | **Study Design** | **Participant N** | **Mean Age in Years** | **Stimulation Modality** | **Anode** | **Cathode** | **Polarity of Stimulating Electrode** | **Online/Offline** | **Duration** | **Current Intensity, Electrode Size** | **Experimental Task Description** | **Semantic/**  **Less semantic** | **Visual/**  **Auditory** | **Verbal/**  **Nonverbal** |
| --- | --- | --- | --- | --- | --- | --- | --- | --- | --- | --- | --- | --- | --- | --- |
| ***Executive Functions*** | | | | | | | | | | | | | | |
| Vannorsdall et al. (2012) [17] | Crossover, sham-controlled, participant-blind trial  But different groups for polarity (anodal, cathodal) | Anodal: 12 Cathodal: 12 | Anodal: 37.9 Cathodal: 33.5 | tDCS | F3 | Cz | Anodal | Online assessment- performed during stimulation | 30 | 1 mA, 27.04 cm2 | **VFT (phonemic, semantic)-** looking into controlled (i.e., switching) and automatic (i.e., clustering) word retrieval processes | Semantic (word) | Visual | Verbal |
|  |  |  |  |  | Cz | F3 | Cathodal |  |  |  |  |  |  |  |
| Fecteau et al. (2013) [18] | Parallel, sham-controlled, double-blind trial | LA/RC: 12 RA/LC: 12 Sham: 12 | LA/RC: 20.3 RA/LC: 22.2 Sham: 22.4 | tDCS | F3 | F4 | Bilateral | Offline assessment- generation task performed after stimulation; Stroop performed before and after stimulation | 20 | 2 mA, 35 cm2 | **Generation of spontaneous verbal responses task-** provide a viable response to open-ended questions  **SCWT** | Semantic (words) | Auditory (verbal responses task), visual (Stroop) | Verbal (verbal generation task), nonverbal (SCWT) |
|  |  |  |  |  | F4 | F3 | Bilateral |  |  |  |  |  |  |  |
| Iuculano & Cohen Kadosh (2013) [19] | Parallel, sham-controlled, participant-blind trial  No. of sessions: 6 | 19 (subgroup n n.r.) | 20-31 | tDCS | F3 | F4 | Bilateral | Offline assessment- performed after stimulation | 20 | 1 mA, 3 cm2 | **Stroop task (numerical)-** stimuli are artificial and everyday digits (learning task administered first to learn artificial digits) | Less semantic | Visual | Nonverbal |
|  |  |  |  |  | P3 | P4 | Bilateral |  |  |  |  |  |  |  |
| Plewnia et al. (2013) [20] | Crossover, sham-controlled, double-blind trial  But different groups for COMT Met, Val-carriers | 46 | 25.87 | tDCS | F3 | Fp2 | Anodal | Online assessment- performed during stimulation | 20 | 1 mA, 35 cm2 | **PGNG-** stimuli are letters | Less semantic | Visual | Nonverbal |
| Balconi & Vitaloni (2014)- Exp. 1 [21] | Crossover, sham-controlled, participant-blind trial | 30 | 24.22 | tDCS | Fp2 | F3 | Cathodal | Offline assessment- performed before (baseline) and immediately after stimulation | 15 | 2 mA, 35 cm2 | **Semantic congruence processing task (action)-** decide whether the final action target frame represents a congruent or an incongruent ending scene (i.e., correctly or incorrectly used object) | Semantic (picture- video) | Visual | Nonverbal |
| Balconi & Vitaloni (2014)- Exp. 2 [21] | Crossover, sham-controlled, participant-blind trial | 28 | 24.18 | tDCS | Fp2 | F3 | Cathodal | Offline assessment- performed before (baseline) and immediately after stimulation | 15 | 2 mA, 35 cm2 | **Semantic congruence processing task (sentence)-** decide whether the final word of a sentence represents a congruent or an incongruent ending | Semantic (word) | Visual | Nonverbal |
| Weber et al. (2014) [22] | Parallel, sham-controlled trial | Active: 11 Sham: 11 | Active: 19-35 Sham: 19-31 | tDCS | F4 | F3 | Bilateral | Offline assessment- performed before and after stimulation | 15 | 1.5 mA, 25 cm2 | **BART** | Less semantic | Visual | Nonverbal |
| Baumert et al. (2020) [23] | Parallel, sham-controlled, participant-blind trial | Anodal: 31 Cathodal: 34 Sham: 31 | 23.04 | tDCS | F3 | Extracephalic (ipsilateral) | Anodal | Online assessment- performed during stimulation | 20 | 1 mA, 35 cm2 | **SCWT** | Semantic (word) | Visual | Nonverbal |
|  |  |  |  |  | Extracephalic (ipsilateral) | F3 | Cathodal |  |  |  |  |  |  |  |
| Nieratschker et al. (2015) [24] | Crossover, sham-controlled, double-blind trial  But different groups for COMT Met/Met, Val/Val | 41 | 24 | tDCS | Fp2 | F3 | Cathodal | Online assessment- performed during stimulation | 20 | 1 mA, 35 cm2 | **PGNG-** stimuli are letters | Less semantic | Visual | Nonverbal |
| Shen et al. (2016)- Exp. 1 [25] | Crossover, sham-controlled, participant-blind trial | 39 | 22.1 | tDCS | F3 | F4 | Bilateral | Online assessment- performed during stimulation | Approx. 20 | 2 mA, 35 cm2 electrodes | **ICT-** make a choice between a sooner-smaller reward and a later-larger reward | Semantic (word) | Visual | Nonverbal |
|  |  |  |  |  | F4 | F3 | Bilateral |  |  |  |  |  |  |  |
| Shen et al. (2016)- Exp. 2A [25] | Crossover, sham-controlled, participant-blind trial | 39 | 21.1 | HD-tDCS | F3 | C3, FT7, Fp1, Fz | Anodal | Online assessment- performed during stimulation | Approx. 20 | 2 mA, approx. 4 cm2 each | **ICT-** make a choice between a sooner-smaller reward and a later-larger reward | Semantic (word) | Visual | Nonverbal |
|  |  |  |  |  | F4 | C4, FT8, Fp2, Fz | Anodal |  |  |  |  |  |  |  |
| Shen et al. (2016)- Exp. 2B [25] | Crossover, sham-controlled, participant-blind trial | 39 | 21.4 | HD-tDCS | C3, FT7, Fp1, Fz | F3 | Cathodal | Online assessment- performed during stimulation | Approx. 20 | 2 mA, approx. 4 cm2 each | **ICT-** make a choice between a sooner-smaller reward and a later-larger reward | Semantic (word) | Visual | Nonverbal |
|  |  |  |  |  | C4, FT8, Fp2, Fz | F4 | Cathodal |  |  |  |  |  |  |  |
| Albein-Urios et al. (2019) [26] | Parallel, sham-controlled, double-blind trial | vlPFC: 15 dmPFC: 15 Sham: 22 | vlPFC: 23 dmPFC: 22 Sham: 21.36 | HD-tDCS | FP2, F6, FC6 | T8, FT8 (right vlPFC) | Cathodal | Offline assessment- performed after stimulation | 20 | 2 mA, 20 mm diameter electrodes | **PRLT-** choose one of two Hiragana characters, receiving either a reward or punishment feedback; correct stimulus is rewarded on 80% of trials, whereas the incorrect stimulus is rewarded on 20% of trials; rule changes intermittently and contingencies reverse, such that choosing the previously correct stimulus leads to punishment until the new correct stimulus is selected | Less semantic | Visual | Nonverbal |
|  |  |  |  |  | Fz, FCZ, CZ, AF4 | F2 (right dmPFC) | Cathodal |  |  |  |  |  |  |  |
| To et al. (2018) [27] | Parallel, sham-controlled, participant-blind trial | Anodal: 15 Cathodal: 15 Sham: 15 | 23.52 | HD-tDCS | Fz | Fp1, Fp2, F7, F8 | Anodal | Offline assessment- performed before (baseline) and after stimulation | 20 | 1 mA, 1 cm radius (circular electrodes) | **Cognitive counting Stroop-** count the number of words in a display of words that denote a number (e.g., word 'three' written twice) compared to a congruent/neutral condition (e.g., the word 'dog' written once)  **TMT B** | Semantic for Stroop (word), less semantic for TMT B | Visual | Nonverbal |
|  |  |  |  |  | Fp1, Fp2, F7, F8 | Fz | Cathodal |  |  |  |  |  |  |  |
| Guo et al. (2018) [28] | Parallel, sham-controlled, participant-blind trial | Anodal: 20 Cathodal: 16 Sham: 22 | Anodal: 21.30 Cathodal: 19.25 Sham: 20.41 | HD-tDCS | F3 | AF3, F1, F5, FC3 | Anodal | Online assessment- performed during stimulation | Approx. 20 | 1.5 mA, approx. 4 cm2 each | **BART** | Less semantic | Visual | Nonverbal |
|  |  |  |  |  | AF3, F1, F5, FC3 | F3 | Cathodal |  |  |  |  |  |  |  |
| Bender et al. (2017)- Exp. 1 [29] | Crossover, sham-controlled, participant-blind trial | 18 | 24 | tDCS | 1 cm posterior to Fz | Extracephalic | Anodal | Offline assessment- performed before, immediately after, and 20 mins after stimulation | 9 | 0.7 mA, 25 cm2 | **Alternative force CRT task-** discriminate between symbols, coloured circles, or sounds; high and low response selection load conditions | Less semantic | Visual & auditory | Nonverbal |
|  |  |  |  |  | Extracephalic | 1 cm posterior to Fz | Cathodal |  |  |  |  |  |  |  |

| **Reference (Year)** | **Study Design** | **Participant N** | **Mean Age in Years** | **Stimulation Modality** | **Anode** | **Cathode** | **Polarity of Stimulating Electrode** | **Online/Offline** | **Duration** | **Current Intensity, Electrode Size** | **Experimental Task Description** | **Semantic/**  **Less semantic** | **Visual/**  **Auditory** | **Verbal/**  **Nonverbal** |
| --- | --- | --- | --- | --- | --- | --- | --- | --- | --- | --- | --- | --- | --- | --- |
| ***Executive Functions*** | | | | | | | | | | | | | | |
| Bender et al. (2017)- Exp. 2 [29] | Crossover, sham-controlled, participant-blind trial | 18 | 21 | tDCS | 1 cm posterior to Fz | Extracephalic | Anodal | Offline assessment- performed before, immediately after, and 20 mins after stimulation | 9 | 0.7 mA, 25 cm2 | **SST-** stimuli are abstract shapes | Less semantic | Visual & auditory | Nonverbal |
|  |  |  |  |  | Extracephalic | 1 cm posterior to Fz | Cathodal |  |  |  |  |  |  |  |
| Bender et al. (2017)- Exp. 3 [29] | Parallel, sham-controlled, participant-blind trial | Active: 18 Sham: 18 | Active: 22 Sham: 22 | tDCS | Extracephalic | 1 cm posterior to Fz | Cathodal | Offline assessment- performed before, immediately after, and 30 mins after stimulation | 13 | 0.7 mA, 25 cm2 | **Modified SST-** stimuli are colours, symbols, or sounds; alternating blocks of "Never Stop" and "Maybe Stop" trials | Less semantic | Visual & auditory | Nonverbal |
| Campanella et al. (2018) [30] | Parallel, sham-controlled, double-blind trial | Active: 18 Sham: 17 | Active: 22.2 Sham: 21.3 | tDCS | F8 | Extracephalic | Anodal | Offline assessment- performed before (session 1) and after (session 2) stimulation | 20 | Current intensity n.r., 25 cm2 | **GNG-** stimuli are letters | Less semantic | Visual | Nonverbal |
| Castro-Meneses et al. (2016) [31] | Crossover, sham-controlled trial | 14 | 22 | tDCS | Intersection point of a line between T4-Fz and a line between F8-Cz | Extracephalic | Anodal | Online/offline assessment- performed before (Phase 1), during (Phase 2), and after (Phase 3) stimulation | 15 | 1.5 mA, 25 cm2 | **SST-** stimuli are with vowels (manual or vocal responses) | Less semantic | Visual | Verbal for vocal responses (overt), nonverbal for manual responses |
| Cunillera et al. (2014) [32] | Crossover, sham-controlled, participant-blind trial | 22 | 21.2 | tDCS | Crossing point between T4-Fz and F8-Cz | Crossing point between T3-Fz and F7-Cz | Bilateral | Offline assessment- performed after stimulation | 18 | 1.5 mA, 9 cm2 | **GNG-SST-** respond to letters or numbers in two separated and consecutive blocks with the right or left hand depending on the side of the appearance of the Go-stimuli; easy and hard discriminability conditions | Less semantic | Visual | Nonverbal |
| Cunillera et al. (2016) [33] | Crossover, sham-controlled, participant-blind trial | 13 | 25.2 | tDCS | Crossing point between T4-Fz and F8-Cz | Crossing point between T3-Fz and F7-Cz | Bilateral | Online assessment- performed during stimulation | 20 | 1.5 mA, 9 cm2 | **GNG-SST-** respond to letters or numbers in two separated and consecutive blocks with the right or left hand depending on the side of the appearance of the Go-stimuli; easy and hard discriminability conditions | Less semantic | Visual | Nonverbal |
| Dambacher et al. (2015) [34] | Parallel, sham-controlled, participant-blind trial | RA/LC: 22 LA/RC: 22 Sham: 20 | 21.89 | tDCS | F8 | F7 | Bilateral | Online assessment- performed before (baseline) and during stimulation | 21.75 | 1.5 mA, 35 cm2 | **GNG-** stimuli are letters | Less semantic | Visual | Nonverbal |
|  |  |  |  |  | F7 | F8 | Bilateral |  |  |  |  |  |  |  |
| Friehs & Frings (2018) [35] | Parallel, sham-controlled, participant-blind trial | Active: 28 Sham: 28 | Active: 25.25 Sham: 24.39 | tDCS | F4 | Extracephalic | Anodal | Offline assessment- performed before and after stimulation | 19 | 0.5 mA, 9 cm2 (active), 35 cm2 (reference) | **SST-** stimuli are arrows (respond to direction) | Less semantic | Visual | Nonverbal |
| Friehs & Frings (2019b) [36] | Parallel, sham-controlled, participant-blind trial | Active: 20 Sham: 22 | Active: 21.80 Sham: 22.32 | tDCS | Extracephalic | F4 | Cathodal | Offline assessment- performed before and after stimulation | 19 | 0.5 mA, 9 cm2 (active), 35 cm2 (reference) | **SST-** stimuli are arrows (respond to direction) | Less semantic | Visual | Nonverbal |

| **Reference (Year)** | **Study Design** | **Participant N** | **Mean Age in Years** | **Stimulation Modality** | **Anode** | **Cathode** | **Polarity of Stimulating Electrode** | **Online/Offline** | **Duration** | **Current Intensity, Electrode Size** | **Experimental Task Description** | **Semantic/**  **Less semantic** | **Visual/**  **Auditory** | **Verbal/**  **Nonverbal** |
| --- | --- | --- | --- | --- | --- | --- | --- | --- | --- | --- | --- | --- | --- | --- |
| ***Executive Functions*** | | | | | | | | | | | | | | |
| Jacobson et al. (2011)- Main Exp. [37] | Crossover, sham-controlled trial | 11 | 28.3 | tDCS | Crossing point between T4-Fz and F8-Cz | Fp1 | Anodal | Offline assessment- performed after stimulation | 10 | 1 mA, 25 cm2 | **SST-** stimuli are shapes | Less semantic | Visual & auditory | Nonverbal |
|  |  |  |  |  | Fp1 | Crossing point between T4-Fz and F8-Cz | Cathodal |  |  |  |  |  |  |  |
|  |  |  |  |  | Crossing point between T4-Fz and F8-Cz | Crossing point between T3-Fz and F7-Cz | Bilateral |  |  |  |  |  |  |  |
|  |  |  |  |  | Crossing point between T3-Fz and F7-Cz | Crossing point between T4-Fz and F8-Cz | Bilateral |  |  |  |  |  |  |  |
| Jacobson et al. (2011)- Control Exp. [37] | Crossover, sham-controlled trial | 11 | 28.3 | tDCS | P4 | Fp1 | Anodal | Offline assessment- performed after stimulation | 10 | 1 mA, 25 cm2 | **SST-** stimuli are shapes | Less semantic | Visual & auditory | Nonverbal |
| Kwon & Kwon (2013a) [38] | Crossover, sham-controlled, participant-blind trial | 40 | 22.97 | tDCS | 4 cm anterior to Cz | Fp1 | Anodal | Offline assessment- performed before and after stimulation | 10 | 1 mA, 35 cm2 | **SST-** stimuli are shapes | Less semantic | Visual | Nonverbal |
| Lapenta et al. (2012) [39] | Crossover, sham-controlled trial | 28 | 23.2 | tDCS | T3, T4 | Extracephalic | Anodal | Online assessment- performed during stimulation | 14 | 1 mA, 35 cm2 | **GNG-** stimuli are kiki and bouba | Less semantic | Visual | Nonverbal |
|  |  |  |  |  | Extracephalic | T3, T4 | Cathodal |  |  |  |  |  |  |  |
| Leite et al. (2018) [40] | Crossover, sham-controlled trial | 16 | 21.5 | tDCS | F8 | F7 | Bilateral | Offline assessment- performed after stimulation | 30 | 1 mA, 35 cm2 | **Prepotent response inhibition task-** attend to a cue and then respond to a target (arrow pointing to left or right); if the presented cue is green, respond left or right, and if the cue colour is red, respond in the opposite direction of the presented target  **GNG-** stimuli are letters | Less semantic | Visual | Nonverbal |
|  |  |  |  |  | F8 | F7 | Bilateral |  |  | 1 mA, 35 cm2 (anode), 100 cm2 (cathode) |  |  |  |  |
| Nejati et al. (2018) [41] | Crossover, sham-controlled, participant-blind trial | 24 | 26.75 | tDCS | F3 | Fp2 | Anodal | Online assessment- performed during stimulation | 20 | 1 mA, 35 cm2 | **GNG-** stimuli are planes positioned in different directions  **TOH**   **BART**  **Delay discounting task-** choose between a smaller, immediate monetary reward and a larger, delayed monetary reward | Less semantic for all except Delay Discounting task (semantic- word) | Visual (also auditory for GNG) | Nonverbal |
|  |  |  |  |  | Fp2 | F3 | Cathodal |  |  |  |  |  |  |  |
| Ouellet et al. (2015) [42] | Parallel, sham-controlled, participant-blind trial | LA/RC: 15 RA/LC: 15 Sham: 15 | LA/RC: 24.07 RA/LC: 27.2 Sham: 24 | tDCS | Over left supraorbital area and lateral to Fp1 (left OFC) | Over right supraorbital area and lateral to Fp2 (right OFC) | Bilateral | Offline assessment- performed before and after stimulation | 30 | 1.5 mA, 35 cm2 (anode), 55.25 cm2 (cathode) | **IGT-** draw cards from four different decks with the goal of earning as much virtual money as possible  **BART**  **SCWT**  **SST-** stimuli are arrows | Less semantic for all tasks except for SCWT (semantic- word) | Visual (also auditory for SST) | Nonverbal |
|  |  |  |  |  | Over right supraorbital area and lateral to Fp2 (right OFC) | Over left supraorbital area and lateral to Fp1 (left OFC) | Bilateral |  |  |  |  |  |  |  |
| Sallard et al. (2018) [43] | Crossover, sham controlled, double-blind | 16 | 23.8 | tDCS | F6-FC6 | Fp1 | Anodal | Offline assessment- performed after stimulation | 20 | 1.5 mA, 35 cm2 (active), 70 cm2 (reference) | **GNG-** stimuli are letters | Less semantic | Visual | Nonverbal |
| Sedgmond et al. (2019) [44] | Parallel, sham-controlled, double-blind trial | Active: 88 Sham: 84 | Active: 20.51 Sham: 21.1 | tDCS | F4 | F3 | Bilateral | Online training task- GNG performed during stimulation  Offline assessment- speeded GNG performed after stimulation | 20 | 2 mA, 35 cm2 | **Food-related GNG-** stimuli are images of healthy and unhealthy foods and filler images; respond to stimulus location  **Speeded version of GNG-** same stimuli as above | Semantic | Visual | Nonverbal |
| Stramaccia et al. (2015) [45] | Parallel, sham-controlled, participant-blind trial | Anodal-IFG: 20 Cathodal-IFG: 20 Anodal-DLPFC: 20 Cathodal-DLPFC: 20 Sham: 35 | Anodal-IFG: 23.95 Cathodal-IFG: 23.35 Anodal-DLPFC: 23.65 Cathodal-DLPFC: 23.10 Sham: 23.06 | tDCS | Crossing point between T4-Fz and F8-Cz | Fp1 | Anodal | Offline assessment- performed after stimulation | 20 | 1.5 mA, 16 cm2 | **SST-** stimuli are shapes | Less semantic | Visual & auditory | Nonverbal |
|  |  |  |  |  | Fp1 | Crossing point between T4-Fz and F8-Cz | Cathodal |  |  |  |  |  |  |  |
|  |  |  |  |  | F4 | Fp1 | Anodal |  |  |  |  |  |  |  |
|  |  |  |  |  | Fp1 | F4 | Cathodal |  |  |  |  |  |  |  |
| Stramaccia et al. (2017)- Exp. 2 [46] | Parallel, sham-controlled, participant-blind trial | Anodal: 24 Cathodal: 24 Sham: 24 | Anodal: 23.96 Cathodal: 23.33 Sham: 23.42 | tDCS | FC4 | Fp1 | Anodal | Online task- performed during stimulation | 20 | 1.5 mA, 16 cm2 | **SST-** stimuli are shapes | Less semantic | Visual & auditory | Nonverbal |
|  |  |  |  |  | Fp1 | FC4 | Cathodal |  |  |  |  |  |  |  |
| Yu et al. (2015)- Exp. 1 [47] | Crossover, sham-controlled trial | 8 | 20-31 | tDCS | Fz | Extracephalic | Anodal | Offline assessment- performed before and after stimulation | 20 | 2 mA, 16 cm2 (active), 35 cm2 (reference) | **SST-** stimuli are arrows (respond to direction) | Less semantic | Visual | Nonverbal |
| Sandrini et al. (2020) [48] | Parallel, sham-controlled, participant-blind trial | Active: 15 Sham: 15 | Active: 26 Sham: 27 | tDCS | Over right pars opercularis | Fp1 | Anodal | Offline assessment- performed before (baseline; session 1) and after stimulation (session 2) | 20 | 1.5 mA, 25 cm2 | **SST-** stimuli are arrows (respond to direction) | Less semantic | Visual | Nonverbal |
| Strobach et al. (2016) [49] | Crossover, sham-controlled, participant-blind trial | 30 | 25.8 | tDCS | Halfway between F3 and C3 | Fp2 | Anodal | Online assessment- performed during stimulation | Approx. 20 | 1 mA, 35 cm2 (active), 100 cm2 (reference) | **Task switching-** switching between letter task (consonant or vowel) and digit task (odd or even) | Less semantic | Visual & auditory | Nonverbal |
|  |  |  |  |  | Fp2 | Halfway between F3 and C3 | Cathodal |  |  |  |  |  |  |  |
| Tayeb & Lavidor (2016) [50] | Crossover, sham-controlled trial | LA/RC: 10 RA/LC: 9 Sham: 23 | 23.9 | tDCS | F3 | F4 | Bilateral | Online assessment- performed during stimulation | 20 | 1.5 mA, 16 cm2 | **Task switching-** switching between magnitude task (bigger or smaller than 5) and parity task (odd or even) | Less semantic | Visual | Nonverbal |
|  |  |  |  |  | F4 | F3 | Bilateral |  |  |  |  |  |  |  |
| Kwon & Kwon (2013b) [51] | Parallel, sham-controlled trial | Active: 20 Sham: 20 | 23 | tDCS | 4 cm anterior to Cz | Extracephalic | Anodal | Online/offline assessment- performed before, during, and after stimulation | 10 | 1 mA, 35 cm2 | **SST-** stimuli are shapes | Less semantic | Visual | Nonverbal |

| **Reference (Year)** | **Study Design** | **Participant N** | **Mean Age in Years** | **Stimulation Modality** | **Anode** | **Cathode** | **Polarity of Stimulating Electrode** | **Online/Offline** | **Duration** | **Current Intensity, Electrode Size** | **Experimental Task Description** | **Semantic/**  **Less semantic** | **Visual/**  **Auditory** | **Verbal/**  **Nonverbal** |
| --- | --- | --- | --- | --- | --- | --- | --- | --- | --- | --- | --- | --- | --- | --- |
| ***Executive Functions*** | | | | | | | | | | | | | | |
| Loftus et al. (2015) [52] | Parallel, sham-controlled trial | Active: 14 Sham: 14 | 24.5 | tDCS | F3 | F4 | Bilateral | Offline assessment- performed before and after stimulation | 10 | 2 mA, 35 cm2 | **Modified SCWT-** decide whether the colour of the letter string (colour word [incongruent] or nonword [neutral]) matches the meaning (name of colour) of the string presented below it in grey | Semantic (word) | Visual | Nonverbal |
| Zmigrod et al. (2016) [53] | Crossover, sham-controlled trial | 14 | 20 | tDCS | F4 | Fp1 | Anodal | Online assessment- performed during stimulation | 20 | 2 mA, 35 cm2 | **Eriksen Flanker task**  **Simon task-** respond to colour of stimulus regardless of its spatial location | Less semantic | Visual | Nonverbal |
|  |  |  |  |  | Fp1 | F4 | Cathodal |  |  |  |  |  |  |  |
| Cattaneo et al. (2011)- Exp. 1 [54] | Crossover, sham-controlled, participant-blind trial | 10 | 23.6 | tDCS | Crossing point between T3-Fz and F7-Cz | Fp2 | Anodal | Offline assessment- performed after stimulation | 20 | 2 mA, 35 cm2 | **VFT (phonemic, semantic)** | Semantic (word) | Visual | Verbal |
| Cattaneo et al. (2011)- Exp. 2 [54] | Crossover, sham-controlled, participant-blind trial | 8 | 23.8 | tDCS | Crossing point between T4-Fz and F8-Cz | Fp1 | Anodal | Offline assessment- performed after stimulation | 20 | 2 mA, 35 cm2 | **VFT (phonemic, semantic)** | Semantic (word) | Visual | Verbal |
| Meinzer et al. (2012) [55] | Crossover, sham-controlled, double-blind trial | 20 | 26.7 | tDCS | Centre of a line connecting the intersection of T3-F3 and F7-C3 and the midpoint between F7-F3 | Fp2 | Anodal | Online assessment- performed during stimulation | Approx. 17 | 1 mA, 35 cm2 (active), 100 cm2 (reference) | **Overt semantic word generation (fluency) test (category)** | Semantic (word) | Visual | Verbal |
| Penolazzi et al. (2013) [56] | Parallel, sham-controlled, participant-blind trial | Frontal: 18 Fronto-temporal: 18 Bi: 18 Uni: 18 Sham: 18 | Frontal: 23.2 Fronto-temporal: 20.7 Bi: 21.7 Uni: 21.4 Sham: 21 | tDCS | Frontal:  Crossing point between T3–Fz and F7–Cz | Fp2 | Anodal | Offline assessment- performed before and after (immediately after stimulation, and 15 mins after end of first post-stimulation test) stimulation | 20 | 2 mA, 35 cm2 | **VFT (category)** | Semantic (word) | Visual | Verbal |
|  |  |  |  |  | Fronto-temporal:  Crossing point between T3–F3 and F7–C3 | Fp2 | Anodal |  |  | 2 mA, 35 cm2 |  |  |  |  |
|  |  |  |  |  | Bilateral:  Crossing point between T3–F3 and F7–C3 | Right homologue area | Bilateral |  |  | 2 mA, 35 cm2 |  |  |  |  |
|  |  |  |  |  | Unilateral:  Crossing point between T3–F3 and F7–C3 | Fp2 | Anodal |  |  | 2 mA, 35 cm2 (active), 100 cm2 (reference) |  |  |  |  |

| **Reference (Year)** | **Study Design** | **Participant N** | **Mean Age in Years** | **Stimulation Modality** | **Anode** | **Cathode** | **Polarity of Stimulating Electrode** | **Online/Offline** | **Duration** | **Current Intensity, Electrode Size** | **Experimental Task Description** | **Semantic/**  **Less semantic** | **Visual/**  **Auditory** | **Verbal/**  **Nonverbal** |
| --- | --- | --- | --- | --- | --- | --- | --- | --- | --- | --- | --- | --- | --- | --- |
| ***Executive Functions*** | | | | | | | | | | | | | | |
| Cohen-Maximov et al. (2015) [57] | Parallel, sham-controlled, participant-blind trial | RA/LC: 13 LA/RC: 13 Sham: 14 | RA/LC: 24.7 LA/RC: 25.5 Sham: 24 | tDCS | Crossing point between T4-Fz and F8-Cz | Crossing point between T3-Fz and F7-Cz | Bilateral | Online assessment- performed during stimulation | 26 | 1.5 mA, 16 cm2 (anode- active), 35 cm2 (cathode- reference) | **Flanker task (attentional load task)-** stimuli are letters, and there are low-load and high-load conditions | Less semantic | Visual | Nonverbal |
|  |  |  |  |  | Crossing point between T3-Fz and F7-Cz | Crossing point between T4-Fz and F8-Cz | Bilateral |  |  |  |  |  |  |  |
| Ehlis et al. (2016)- Exp. 1 [58] | Crossover, sham-controlled, double-blind trial  But different groups for polarity (anodal, cathodal) | 23 | 32.1 | tDCS | Between C3, F3 and F7, i.e., centre of the electrode located between FC5 and FC3 | Fp2 | Anodal | Offline assessment- performed after stimulation | 20 | 1 mA, 35 cm2 | **VFT (phonemic, semantic)** | Semantic (word) | Visual | Verbal |
| Ehlis et al. (2016)- Exp. 2 [58] | Crossover, sham-controlled, double-blind trial  But different groups for polarity (anodal, cathodal) | 23 | 24.3 | tDCS | Fp2 | Between C3, F3 and F7, i.e., centre of the electrode located between FC5 and FC3 | Cathodal | Offline assessment- performed after stimulation | 20 | 1 mA, 35 cm2 | **VFT (phonemic, semantic)** | Semantic (word) | Visual | Verbal |
| Vannorsdall et al. (2016) [59] | Crossover, sham-controlled, participant-blind trial | 14 | 22.3 | tDCS | Intersection of two lines connecting T3 to Fz and F7 to Cz | Fp2 | Anodal | Offline assessment- performed after stimulation | 20 | 2 mA, 35 cm2 | **VFT (phonemic, semantic)** | Semantic (word) | Visual | Verbal |
| Chrysikou et al. (2013) [60] | Parallel, sham-controlled, double-blind trial | L/CU: 8 R/CU: 8 S/CU: 8 L/UU: 8 R/UU: 8 S/UU: 8 | 23.38 | tDCS | Extracephalic | F7 | Cathodal | Online assessment- performed during stimulation | Max. 20 | 1.5 mA, 25 cm2 | Fluency task – generate common uses of daily objects | Semantic (picture) | Visual | Verbal |
|  |  |  |  |  | Extracephalic | F8 | Cathodal |  |  |  |  |  |  |  |
| Balconi et al. (2014)- Exp. 2 [61] | Crossover, sham-controlled trial | 23 | 25.6 | tDCS | P3 | Fp2 | Anodal | Offline assessment- performed after stimulation | 13 | 2 mA, 35 cm2 | **Error detection task (semantic congruence processing)-** decide whether the final action target frame represents a congruent or an incongruent ending scene (i.e., correctly or incorrectly used object); for incongruent trials, half include actions that represents an instrumental incongruence (incorrect use of the object, as grasping a bat upside-down), and half includes actions that represents a functional incongruence (incongruous use, such as brushing teeth with a comb) | Semantic (picture) | Visual | Nonverbal |

| **Reference (Year)** | **Study Design** | **Participant N** | **Mean Age in Years** | **Stimulation Modality** | **Anode** | **Cathode** | **Polarity of Stimulating Electrode** | **Online/Offline** | **Duration** | **Current Intensity, Electrode Size** | **Experimental Task Description** | **Semantic/**  **Less semantic** | **Visual/**  **Auditory** | **Verbal/**  **Nonverbal** |
| --- | --- | --- | --- | --- | --- | --- | --- | --- | --- | --- | --- | --- | --- | --- |
| ***Executive Functions*** | | | | | | | | | | | | | | |
| Gbadeyan et al. (2016) [62] | Crossover, sham-controlled, double-blind trial  But different groups for stim. site (L/R DLPFC, L/R M1 [latter excluded]) | L: 30 R: 30 | L: 25.96 R: 26.60 | HD-tDCS | F3 | Four electrodes symmetrically around the centre electrode | Anodal | Online assessment- performed during stimulation | 20 | 1 mA, centre electrode- diameter 2.5 cm, ring-shaped reference electrode- inner diameter 9.2 cm; outer diameter 11.5 cm | **Flanker task-** there are four conditions: previous congruent/current congruent, previous congruent/current incongruent, previous incongruent/current congruent, previous incongruent/current incongruent | Less semantic | Visual | Nonverbal |
|  |  |  |  |  | F4 | Four electrodes symmetrically around the centre electrode | Anodal |  |  |  |  |  |  |  |
| Mansouri et al. (2016)- Exp. 1 [63] | Crossover, sham-controlled, participant-blind trial | 19 | 19-28 | tDCS | F3 | Fp2 | Anodal | Offline assessment- performed before and after stimulation | 10 | 1 mA, 7.5 cm2 (active), 20 cm2 (reference) | **WCST** | Less semantic | Visual | Nonverbal |
|  |  |  |  |  | Fp2 | F3 | Cathodal |  |  |  |  |  |  |  |
| Mansouri et al. (2016)- Exp. 2 [63] | Crossover, sham-controlled, participant-blind trial | 23 | 20-32 | tDCS | F3  *Note:* F4 targeted for 2 Ps in SST (left-handed) | Fp2 | Anodal | Offline assessment- performed before and after stimulation | 10 | 1 mA, 7.5 cm2 (active), 20 cm2 (reference) | **SST-** stimuli are horizontal/vertical bars | Less semantic | Visual | Nonverbal |
|  |  |  |  |  | Fp2 | F3 | Cathodal |  |  |  |  |  |  |  |
| Sdoia et al. (2020) [64] | Crossover, sham-controlled trial | 20 | 26.3 | tDCS | F4 | F3 | Bilateral | Online assessment- performed during stimulation | n.r. | 1.5 mA, 3 cm diameter | **Task switching-** switching between magnitude task (smaller or larger than five), parity task (odd or even), and position task (digit centrally or peripherally positioned); either switch back to a previously inhibited task, switch back to a non-inhibited task, or repeat the same task performed on previous trial | Less semantic | Visual | Nonverbal |
|  |  |  |  |  | P4 | P3 | Bilateral |  |  |  |  |  |  |  |
| Lu et al. (2021) [65] | Parallel, sham-controlled, participant-blind trial  No. of sessions: 9 | Active: 22 Sham: 21 | Active: 20.73 Sham: 21.10 | HD-tDCS | F3 | AF3, F1, F5, FC3 | Anodal | Offline assessment- performed before and after stimulation  *Note:* Phase 1 (Stroop and SAT), Phase 2 (9 tDCS sessions were undertaken across 18 days, with SAT completed on the day after the 5th HD-tDCS session), Phase 3 (Stroop and SAT); the three phases are all on different days | 20 | 1.5 mA, electrode size n.r. | **SCWT**  **SAT-** match geometric objects according to colour or shape depending on cue | Semantic (word) | Visual | Nonverbal |
| Hasan et al. (2023) [66] | Parallel, sham-controlled, participant-blind trial  No. of sessions: 5 | Active: 19 Sham: 21 | 22 | tDCS | F3 | F4 | Bilateral | Offline assessment- performed before and after stimulation | 30 | 2 mA, 20 cm2 | **SCWT** | Semantic (word) | Visual | Nonverbal |
| Fujiyama et al. (2022) [67] | Crossover, sham-controlled, double-blind trial  But different groups for pre SMA, rIFG | PreSMA: 20 rIFG: 22 | PreSMA: 22.4 rIFG: 24 | tDCS | Fz | Fp1 | Anodal | Offline assessment- performed before and after stimulation | 20 | 1.5 mA, 24 cm2 | **Flashing grid task**- judge whether stimulus (mixture of blue and orange cells) contains a greater proportion of orange or blue cells (pre-cued as to which response option is more likely), and withhold response to stop signal | Less semantic | Visual | Nonverbal |
|  |  |  |  |  | Crossing point between T4-Fz and F8-Cz | Fp1 | Anodal |  |  |  |  |  |  |  |
| Perrotta et al. (2021)- Exp. 1 [68] | Crossover, sham-controlled trial | 12 | 26 | tDCS | Crossing point between the lines connecting T4-Fz and F8-Cz | Crossing point between the lines connecting T3-Fz and F7-Cz | Bilateral | Offline assessment- performed before and after stimulation | 18 | 1.5 mA, 9 cm2 | **GNG-** stimuli are shapes | Less semantic | Visual | Nonverbal |
| Perrotta et al. (2021)- Exp. 2 [68] | Crossover, sham-controlled trial | 19 | 22.45 | tDCS | Crossing point between the lines connecting T4-Fz and F8-Cz | Fp1 | Anodal | Offline assessment- performed before and after stimulation | 20 | 2 mA, 25 cm2 | **Simple 3-back + SCWT, complex 3-back + SCWT:**  **Simple 3-back-** respond every time the second n-back stimulus (horizontal or vertical) appears  **Complex 3-back-** respond only when visual configuration of second n-back stimulus matches configuration of n-3 stimulus (i.e., horizontal-horizontal, vertical-vertical) | Semantic (word) for Stroop, less semantic for 3-back | Visual | Nonverbal |
| Perrotta et al. (2021)- Exp. 3 [68] | Parallel, sham-controlled trial | Active: 9 Sham: 8 | Active: 31.1 Sham: 23.25 | tDCS | F4 | F3 | Bilateral | Offline assessment- performed before and after stimulation | 20 | 2 mA, 25 cm2 | **SCWT** | Semantic (word) | Visual | Nonverbal |
| Mattavelli et al. (2022) [69] | Crossover, sham-controlled, participant-blind trial | 20 | 23.5 | HD-tDCS | Fz, F1, FCz (dACC) | PO9, O9, O10 | Anodal | Offline assessment- performed after stimulation | 20 | 1 mA, 9.5 mm radius electrodes | **Flanker task**  **Gambling tasks-** chose between a risky mixed-gamble resulting in two equally probable outcomes and a guaranteed alternative; **loss aversion task:** there are gain–loss trials requiring to choose between a gamble with equally probable variable gains or losses and a guaranteed alternative of 0; **risk aversion task:** there are gain-only trials requiring one to choose between a gamble with equally probable variable positive or 0 outcomes and a guaranteed gain | Less semantic | Visual | Nonverbal |
|  |  |  |  |  | PO9, O9, O10 | Fz, F1, FCz (dACC) | Cathodal |  |  |  |  |  |  |  |
| Klaus & Hartwigsen (2020) [70] | Crossover, sham-controlled, double-blind trial | 48 | 27.06 | tDCS | Between FC5 and C5 | Centre of forehead | Anodal | Offline assessment- performed after stimulation | 20 | 2 mA, 25 cm2 (active), 100 cm2 (reference) | **VFT (phonemic, semantic)** | Semantic (word) | Visual | Verbal |
| Westwood & Romani (2018)- Exp. 1 [71] | Crossover, sham-controlled, participant-blind trial  But different groups for recent-probe and semantic-associated probe (in WM section) | 39 | 19.5 | tDCS | F7 | Fp2 | Anodal | Online assessment- performed during stimulation | 25 | 1.5 mA, 25 cm2 (active), 35 cm2 (reference) | **VFT (phonemic, semantic)** | Semantic (word) | Visual | Verbal |
| Lo et al. (2019) [72] | Crossover, sham-controlled trial | 26 | 24.4 | tDCS | P4 | Fp1 | Anodal | Offline assessment- performed before and after stimulation | 20 | 1.5 mA, 35 cm2 | **ANT (only executive effect)-** press arrow (right or left) corresponding to the direction of a target arrow appearing above or below the centre fixation cross; pre-cue conditions: no (only central fixation cross), central (asterisk appears on top of central fixation cross), double (two asterisks appear both above and below central fixation cross), or spatial cues (asterisk appears either above or below central fixation cross); Flanker conditions: neutral, congruent, incongruent | Less semantic | Visual | Nonverbal |
| Marko & Riečanský (2021) [73] | Parallel, sham-controlled, double-blind trial | PFC: 40 TPC: 40 Sham: 41 | 23.1 | tDCS | Between F3 and AF3 | Between T7 and P5 | Bilateral | Online/offline assessment- performed before, during, and after stimulation | 25 | 2 mA, 35 cm2 | **Associative chain test (lexical-semantic retrieval)-** continuously generate word chains according to specific conditions, starting with a word stimulus; associate condition: each new response is semantically associated to the previous one; dissociate condition: words are semantically unrelated; associate–dissociate condition: deliver associations and dissociations in alternation | Semantic (word) | Visual | Nonverbal |
|  |  |  |  |  | Between T8 and P6 | Between T7 and P5 | Bilateral |  |  |  |  |  |  |  |
| Lu et al. (2020) [74] | Parallel, sham-controlled, participant-blind trial  No. of sessions: 12 | Active: 21 Sham: 18 | Active: 21.57 Sham: 20.67 | HD-tDCS | F3 | AF3, F1, F5, FC3 | Anodal | Offline assessment- performed before and after stimulation | 20 | 1.5 mA, electrode size n.r. | **ANT (only executive effect)-** press arrow (right or left) corresponding to the direction of a target arrow appearing above or below the centre fixation cross; pre-cue conditions: no (only central fixation cross), central (asterisk appears on top of central fixation cross), double (two asterisks appear both above and below central fixation cross), or spatial cues (asterisk appears either above or below central fixation cross); Flanker conditions: neutral, congruent, incongruent  **SCWT** | Semantic (word) for Stroop, less semantic for ANT | Visual | Nonverbal |
| Dubreuil-Vall et al. (2019) [75] | Crossover, sham-controlled, double-blind trial | 20 | 32.6 | tDCS | F3 | Fp2 | Anodal | Offline assessment- performed before and after stimulation | 30 | 2 mA, 3.14 cm2 | **Eriksen Flanker task** | Less semantic | Visual | Nonverbal |
|  |  |  |  |  | F4 | Fp1 | Anodal |  |  |  |  |  |  |  |

| **Reference (Year)** | **Study Design** | **Participant N** | **Mean Age in Years** | **Stimulation Modality** | **Anode** | **Cathode** | **Polarity of Stimulating Electrode** | **Online/Offline** | **Duration** | **Current Intensity, Electrode Size** | **Experimental Task Description** | **Semantic/**  **Less semantic** | **Visual/**  **Auditory** | **Verbal/**  **Nonverbal** |
| --- | --- | --- | --- | --- | --- | --- | --- | --- | --- | --- | --- | --- | --- | --- |
| ***Attention*** | | | | | | | | | | | | | | |
| Bolognini et al. (2010a)- Exp. 3 [76] | Crossover, sham-controlled, double-blind trial | 12 | 26 | tDCS | P4 | Extracephalic | Anodal | Offline assessment- performed before and after stimulation | 15 | 2 mA, 35 cm2 | **E-F test-** required to scan visual field, searching for targets embedded among distracters | Less semantic | Visual | Nonverbal |
|  |  |  |  |  | P3 | Extracephalic | Anodal |  |  |  |  |  |  |  |
| Bolognini et al. (2010b) [77] | Parallel, sham-controlled, double-blind trial | Active: 16 Sham: 16 | Active: 24 Sham: 22 | tDCS | P4 | Extracephalic | Anodal | Offline assessment- performed before (baseline) and after stimulation (different days) | 15 | 2 mA, 35 cm2 | **RSE task-** randomly presented with a single auditory stimulus, single red or blue visual stimulus (presented to left or right of fixation), or red or blue audiovisual stimulus (white noise delivered from loudspeakers placed at the same height as the visual target); respond to onset of target and refrain from responding to catch trials | Less semantic | Visual | Nonverbal |
| Iyer et al. (2005)- Exp. 1 [4] | Parallel, sham-controlled, participant-blind trial | Anodal: 10 Cathodal: 10 Sham: 10 | Anodal: 35.9 Cathodal: 36.4 Sham: 36.7 | tDCS | F3 | Fp2 | Anodal | Offline assessment- performed before and after stimulation | 20 | 1 mA, 25 cm2 | **CalCAP-** 1) simple reaction time task, 2) single digit recognition task, 3) one-back cued response task, 4) one-increment task (responding when two numbers are in ascending sequence) | Less semantic | Visual | Nonverbal |
|  |  |  |  |  | Fp2 | F3 | Cathodal |  |  |  |  |  |  |  |
| Iyer et al. (2005)- Exp. 2 [4] | Parallel, sham-controlled, participant-blind trial | Anodal: 14 Cathodal: 14 Sham: 15 | Anodal: 36.4 Cathodal: 39.5 Sham: 36.7 | tDCS | F3 | Fp2 | Anodal | Online assessment- performed before and during stimulation | 20 | 1 mA, 25 cm2 | **CalCAP-** 1) simple reaction time task, 2) single digit recognition task, 3) one-back cued response task, 4) one-increment task (responding when two numbers are in ascending sequence) | Less semantic | Visual | Nonverbal |
|  |  |  |  |  | Fp2 | F3 | Cathodal |  |  |  |  |  |  |  |
| Iyer et al. (2005)- Exp. 3 [4] | Parallel, sham-controlled, participant-blind trial | Anodal: 10 Cathodal: 10 Sham: 10 | Anodal: 38.7 Cathodal: 37.7 Sham: 39.5 | tDCS | F3 | Fp2 | Anodal | Online assessment- performed before and during stimulation | 20 | 2 mA, 25 cm2 | **CalCAP-** 1) simple reaction time task, 2) single digit recognition task, 3) one-back cued response task, 4) one-increment task (responding when two numbers are in ascending sequence) | Less semantic | Visual | Nonverbal |
|  |  |  |  |  | Fp2 | F3 | Cathodal |  |  |  |  |  |  |  |
| Sparing et al. (2009)- Exp. 1 [78] | Crossover, sham-controlled, participant-blind trial  But hemisphere is between-subjects | L: 10 R: 10 | 28.5 | tDCS | P3 | Cz | Anodal | Offline assessment- performed before (baseline) and after stimulation (immediately after and 20 mins following cessation of tDCS) | 10 | 1 mA, 25 cm2 (active), 35 cm2 (reference) | **Visual detection task-** respond with different button presses according to whether stimuli (small dots) are unilateral left visual stimuli, unilateral right stimuli, or bilateral stimuli | Less semantic | Visual | Nonverbal |
|  |  |  |  |  | P4 | Cz | Anodal |  |  |  |  |  |  |  |
|  |  |  |  |  | Cz | P3 | Cathodal |  |  |  |  |  |  |  |
|  |  |  |  |  | Cz | P4 | Cathodal |  |  |  |  |  |  |  |
| Elmer et al. (2009) [79] | Crossover, sham-controlled trial  But stim. side is between-subjects | L: 10 R: 10 | 22.3 | tDCS | F3 | Extracephalic | Anodal | Offline assessment- performed before and after stimulation | 5 | 1.5 mA, 28 cm2 (active), 100 cm2 (reference) | **d2 test** | Less semantic | Visual | Nonverbal |
|  |  |  |  |  | Extracephalic | F3 | Cathodal |  |  |  |  |  |  |  |
|  |  |  |  |  | F4 | Extracephalic | Anodal |  |  |  |  |  |  |  |
|  |  |  |  |  | Extracephalic | F4 | Cathodal |  |  |  |  |  |  |  |
| Leite et al. (2013) [14] | Crossover, sham-controlled trial | 16 | 24 | tDCS | F3 | F4 | Bilateral | Online assessment- performed during stimulation | Max. 30 | 2 mA, 35 cm2 | **SRT task  CRT task** | Less semantic | Visual | Nonverbal |
|  |  |  |  |  | F4 | F3 | Bilateral |  |  |  |  |  |  |  |
| Motohashi et al. (2013) [80] | Crossover, sham-controlled, participant-blind trial  But two different groups regarding stim. order  No. of sessions: 4 | Group A: 6 Group B: 6 | 22 | tDCS | F3 | Fp2 | Anodal | Offline assessment- performed before (baseline; T0) and after (T1, T2) stimulation | 20 | 1 mA, 35 cm2 | **CogHealth:**  **DT-** a single card is presented face-down; respond whenever the card turns face-up  **IT-** respond to the colour of the card (red or black)  **OBT-** respond whether or not the face-up card is exactly the same as the immediately previous card  **CMT-** respond when any one of the cards touches one of the horizontal lines | Less semantic | Visual | Nonverbal |
| Sdoia et al. (2019) [81] | Crossover, sham-controlled, participant-blind trial  But different groups for anodal and cathodal stim. | Anodal: 17 Cathodal: 17 | Anodal: 21.45 Cathodal: 23.05 | tDCS | F3 | Fp2 | Anodal | Online assessment- performed during stimulation | 10 | 1.5 mA, diameter: 3 cm (circular electrodes) | **AB task** | Less semantic | Visual | Nonverbal |
|  |  |  |  |  | Fp2 | F3 | Cathodal |  |  |  |  |  |  |  |
| Brückner & Kammer (2017) [82] | Parallel, sham-controlled trial | Anodal: 20 Cathodal: 20 Sham: 20 | Anodal: 21.5 Cathodal: 23.7 Sham: 22.9 | tDCS | CP5 | Extracephalic (ipsilateral) | Anodal | Offline assessment- performed before and after stimulation | 15 | 1 mA, 35 cm2 | **CRT task** | Less semantic | Visual | Nonverbal |
|  |  |  |  |  | Extracephalic (ipsilateral) | CP5 | Cathodal |  |  |  |  |  |  |  |
| To et al. (2018) [27] | Parallel, sham-controlled, participant-blind trial | Anodal: 15 Cathodal: 15 Sham: 15 | 23.52 | HD-tDCS | Fz | Fp1, Fp2, F7, F8 | Anodal | Offline assessment- performed before (baseline) and after stimulation | 20 | 1 mA, 1 cm radius (circular electrodes) | **TMT A** | Less semantic | Visual | Nonverbal |
|  |  |  |  |  | Fp1, Fp2, F7, F8 | Fz | Cathodal |  |  |  |  |  |  |  |
| Jacobson et al. (2011)- Main Exp. [37] | Crossover, sham-controlled trial | 11 | 28.3 | tDCS | Crossing point between T4-Fz and F8-Cz | Fp1 | Anodal | Offline assessment- performed after stimulation | 10 | 1 mA, 25 cm2 | **Control VDT-** visual discrimination of shapes with no stop signals | Less semantic | Visual | Nonverbal |
|  |  |  |  |  | Fp1 | Crossing point between T4-Fz and F8-Cz | Cathodal |  |  |  |  |  |  |  |
|  |  |  |  |  | Crossing point between T4-Fz and F8-Cz | Crossing point between T3-Fz and F7-Cz | Bilateral |  |  |  |  |  |  |  |
|  |  |  |  |  | Crossing point between T3-Fz and F7-Cz | Crossing point between T4-Fz and F8-Cz | Bilateral |  |  |  |  |  |  |  |
| Jacobson et al. (2011)- Control Exp. [37] | Crossover, sham-controlled trial | 11 | 28.3 | tDCS | P4 | Fp1 | Anodal | Offline assessment- performed after stimulation | 10 | 1 mA, 25 cm2 | **Control VDT-** visual discrimination of shapes with no stop signals | Less semantic | Visual | Nonverbal |

| **Reference (Year)** | **Study Design** | **Participant N** | **Mean Age in Years** | **Stimulation Modality** | **Anode** | **Cathode** | **Polarity of Stimulating Electrode** | **Online/Offline** | **Duration** | **Current Intensity, Electrode Size** | **Experimental Task Description** | **Semantic/**  **Less semantic** | **Visual/**  **Auditory** | **Verbal/**  **Nonverbal** |
| --- | --- | --- | --- | --- | --- | --- | --- | --- | --- | --- | --- | --- | --- | --- |
| ***Attention*** |  |  |  |  |  |  |  |  |  |  |  |  |  |  |
| Leite et al. (2018) [40] | Crossover, sham-controlled trial | 16 | 21.5 | tDCS | F8 | F7 | Bilateral | Offline assessment- performed after stimulation | 30 | 1 mA, 35 cm2 | **CRT task** | Less semantic | Visual | Nonverbal |
|  |  |  |  |  | F8 | F7 | Bilateral |  |  | 1 mA, 35 cm2 (anode), 100 cm2 (cathode) |  |  |  |  |
| Ouellet et al. (2015) [42] | Parallel, sham-controlled, participant-blind trial | LA/RC: 15 RA/LC: 15 Sham: 15 | LA/RC: 24.07 RA/LC: 27.20 Sham: 24 | tDCS | Over left supraorbital area and lateral to Fp1 | Over right supraorbital area and lateral to Fp2 | Bilateral | Offline assessment- performed before and after stimulation | 30 | 1.5 mA, 35 cm2 (anode), 55.25 cm2 (cathode) | **CPT-** respond to a specific letter among consecutive flashes of various letters | Less semantic | Visual | Nonverbal |
|  |  |  |  |  | Over right supraorbital area and lateral to Fp2 | Over left supraorbital area and lateral to Fp1 | Bilateral |  |  |  |  |  |  |  |
| Pisoni et al. (2012)- Exp. 1 [83] | Crossover, sham-controlled, participant-blind trial | 12 | 22.4 | tDCS | Talairach coordinates: X = −50; Y = −46; Z = 1 (left STG- Wernicke's area) | Fp2 | Anodal | Offline assessment- performed after stimulation | 20 | 2 mA, 35 cm2 | **Visual attentive control task-** indicate whether the target (an object that differs in orientation amongst a configuration of stimuli) appears in the left or right hemifield | Less semantic | Visual | Nonverbal |
| Pisoni et al. (2012)- Exp. 2 [83] | Crossover, sham-controlled, participant-blind trial | 12 | 21.8 | tDCS | Crossing point between T3-Fz and F7-Cz (left IFG- Broca's area) | Fp2 | Anodal | Offline assessment- performed after stimulation | 20 | 2 mA, 35 cm2 | **Visual attentive control task-** indicate whether the target (an object that differs in orientation amongst a configuration of stimuli) appears in the left or right hemifield | Less semantic | Visual | Nonverbal |
| Cattaneo et al. (2011)- Exp. 1 [54] | Crossover, sham-controlled, participant-blind trial | 10 | 23.6 | tDCS | Crossing point between T3-Fz and F7-Cz | Fp2 | Anodal | Offline assessment- performed after stimulation | 20 | 2 mA, 35 cm2 | **Spatial DT-** indicate whether target appears in the left or right square | Less semantic | Visual | Nonverbal |
| Cattaneo et al. (2011)- Exp. 2 [54] | Crossover, sham-controlled, participant-blind trial | 8 | 23.8 | tDCS | Crossing point between T4-Fz and F8-Cz | Fp1 | Anodal | Offline assessment- performed after stimulation | 20 | 2 mA, 35 cm2 | **Spatial DT-** indicate whether target appears in the left or right square | Less semantic | Visual | Nonverbal |

| **Reference (Year)** | **Study Design** | **Participant N** | **Mean Age in Years** | **Stimulation Modality** | **Anode** | **Cathode** | **Polarity of Stimulating Electrode** | **Online/Offline** | **Duration** | **Current Intensity, Electrode Size** | **Experimental Task Description** | **Semantic/**  **Less semantic** | **Visual/**  **Auditory** | **Verbal/**  **Nonverbal** |
| --- | --- | --- | --- | --- | --- | --- | --- | --- | --- | --- | --- | --- | --- | --- |
| ***Attention*** | | | | | | | | | | | | | | |
| Penolazzi et al. (2013) [56] | Parallel, sham-controlled, participant-blind trial | Frontal: 18 Fronto-temporal: 18 Bi: 18 Uni: 18 Sham: 18 | Frontal: 23.2 Fronto-temporal: 20.7 Bi: 21.7 Uni: 21.4 Sham: 21 | tDCS | Frontal:  Crossing point between T3–Fz and F7–Cz | Fp2 | Anodal | Offline assessment- performed before and after stimulation | 20 | 2 mA, 35 cm2 | **SRT task** | Less semantic | Visual | Nonverbal |
|  |  |  |  |  | Fronto-temporal:  Crossing point between T3–F3 and F7–C3 | Fp2 | Anodal |  |  | 2 mA, 35 cm2 |  |  |  |  |
|  |  |  |  |  | Bilateral:  Crossing point between T3–F3 and F7–C3 | Right homologue area | Bilateral |  |  | 2 mA, 35 cm2 |  |  |  |  |
|  |  |  |  |  | Unilateral:  Crossing point between T3–F3 and F7–C3 | Fp2 | Anodal |  |  | 2 mA, 35 cm2 (active), 100 cm2 (reference) |  |  |  |  |
| Ihara et al. (2015) [84] | Crossover, sham-controlled trial | 14 | 22-60 | tDCS | Electrode centre over median of Talairach coordinates −45, 28, −16 and −52, 24, 23 (left anterior inferior cortex [LaIFC] and left posterior inferior cortex [LpIFC], respectively) | Cz | Anodal | Online assessment- performed during stimulation | 15 | 1.5 mA, 35 cm2 | **SRT task** | Less semantic | Visual | Nonverbal |
| Price et al. (2016) [85] | Crossover, sham-controlled, participant blind trial | 18 | 25.3 | HD-tDCS | CP5 | C3, T7, P7, P3 | Anodal | Offline assessment- performed after stimulation | 20 | 2 mA, outer diameter: 12 mm, inner diameter: 6 mm (ring electrodes) | **Letter-string control task-** decide whether two letter strings match (e.g., pnqvt pnqvt) or not (e.g., vsbsl vsbql) | Less semantic | Visual | Nonverbal |
|  |  |  |  |  | CP6 | C4, T8, P8, P4 | Anodal |  |  |  |  |  |  |  |
| Giustolisi et al. (2018) [86] | Parallel, sham-controlled, participant-blind trial | Active: 22 Sham: 22 | 22 | tDCS | F5 | Fp2 | Anodal | Online assessment- performed during stimulation | 30 | 0.75 mA, 9 cm2 (active), 35 cm2 (reference) | **Visual pattern task-** decide which of the two checkerboards presented side-by-side is identical to the previously presented single checkerboard | Less semantic | Visual | Nonverbal |
| Gan et al. (2022) [87] | Crossover, sham-controlled, participant-blind trial | 27 | 19.19 | tDCS | 5 cm lateral toward the left and 4 cm anterior from the vertex | Fp2 | Anodal | Online/offline assessment- performed during (blocks 1-3) and after (block 4) stimulation | 28.8 | 2 mA, 25 cm2 | **Vigilance task (visual search)-** presented with search arrays consisting of blue circles and red squares; respond whether or not red circle was detected | Less semantic | Visual | Nonverbal |

| **Reference (Year)** | **Study Design** | **Participant N** | **Mean Age in Years** | **Stimulation Modality** | **Anode** | **Cathode** | **Polarity of Stimulating Electrode** | **Online/Offline** | **Duration** | **Current Intensity, Electrode Size** | **Experimental Task Description** | **Semantic/**  **Less semantic** | **Visual/**  **Auditory** | **Verbal/**  **Nonverbal** |
| --- | --- | --- | --- | --- | --- | --- | --- | --- | --- | --- | --- | --- | --- | --- |
| ***Attention*** | | | | | | | | | | | | | | |
| Paladini et al. (2020) [88] | Crossover, sham-controlled trial | 25 | 23.16 | tDCS | P3 | P4 | Bilateral | Online assessment- performed during stimulation | 20 | 1.5 mA, 35 cm2 | **Dual task- non-spatial verbal WM task** (memorise low or high cognitive load consonants), which allows to manipulate cognitive load, and a **visuospatial attention detection task** (respond when lateralised stimulus appears; **relevant to us**) | Less semantic | Visual | Nonverbal |
|  |  |  |  |  | P4 | P3 | Bilateral |  |  |  |  |  |  |  |
| Orth et al. (2023) [89] | Parallel, sham-controlled, double-blind trial | L: 64 R: 64  Sham: 126 | L: 23.92 R: 22.7 Sham: 23.31 | tDCS | F3 | Fp2 | Anodal | Online task- performed during stimulation | 20 | 1 mA, 35 cm2 | **Alertness task-** respond to appearance of a cross, either preceded by auditory cue or not (phasic or intrinsic alertness) | Less semantic | Visual & auditory | Nonverbal |
|  |  |  |  |  | F4 | Fp1 | Anodal |  |  |  |  |  |  |  |

| **Reference (Year)** | **Study Design** | **Participant N** | **Mean Age in Years** | **Stimulation Modality** | **Anode** | **Cathode** | **Polarity of Stimulating Electrode** | **Online/Offline** | **Duration** | **Current Intensity, Electrode Size** | **Experimental Task Description** | **Semantic/**  **Less semantic** | **Visual/**  **Auditory** | **Verbal/**  **Nonverbal** |
| --- | --- | --- | --- | --- | --- | --- | --- | --- | --- | --- | --- | --- | --- | --- |
| ***Working Memory*** | | | | | | | | | | | | | | |
| Berryhill et al. (2010) [90] | Crossover, sham-controlled, participant-blind trial | 11 | 25 | tDCS | P4 | Extracephalic | Anodal | Offline assessment- performed after stimulation  Online task (practice object WM task)- performed during stimulation; outcome not measured | 10 | 1.5 mA, 35 cm2 | **Object WM task-** a series of images are presented followed by a mask; either have to recall (verbal report) or recognise (old/new) the images held in memory | Semantic (picture) | Visual | Nonverbal stimuli verbal response |
|  |  |  |  |  | Extracephalic | P4 | Cathodal |  |  |  |  |  |  |  |
| Chi et al. (2010) [91] | Parallel, sham-controlled, participant-blind trial | LC/RA: 12 LA/RC : 12 Sham: 12 | 23 | tDCS | Halfway between T8 and FT8 | Halfway between T7 and FT7 | Bilateral | Online assessment- performed before and during stimulation | Approx. 13 | 2 mA, 35 cm2 | **Visual memory task-** during study phase, presented with multiple slides of stimuli (different types of shapes which vary in number, arrangement, colour, and size) that are related by a common theme; in test phase, engage in recognition task which additionally presented slides | Less semantic | Visual | Nonverbal |
|  |  |  |  |  | Halfway between T7 and FT7 | Halfway between T8 and FT8 | Bilateral |  |  |  |  |  |  |  |
| Fregni et al. (2005)- Main Exp. [92] | Crossover, sham-controlled, participant-blind trial | 15 | 20.2 | tDCS | F3 | Fp2 | Anodal | Online assessment- performed during stimulation | 10 | 1 mA, 35 cm2 | ***n*-back task (3-back)-** stimuli are letters | Less semantic | Visual | Nonverbal |
| Fregni et al. (2005)- Control Exp. [92] | Crossover, sham-controlled, participant-blind trial | 7 | 20.2 (for 15 ps) | tDCS | Fp2 | F3 | Cathodal | Online assessment- performed during stimulation | 10 | 1 mA, 35 cm2 | ***n*-back task (3-back)-** stimuli are letters | Less semantic | Visual | Nonverbal |
| Ohn et al. (2008) [93] | Crossover, sham-controlled, participant-blind trial | 15 | 26.5 | tDCS | F3 | Fp2 | Anodal | Online/offline assessment- performed before (baseline), during (three times; T1-T3), and after stimulation (T4) | 30 | 1 mA, 25 cm2 | ***n*-back task (3-back)-** stimuli are Korean letters | Less semantic | Visual | Nonverbal |
| Mulquiney et al. (2011) [94] | Crossover, sham-controlled, participant-blind trial | 10 | 29.42 | tDCS | F3 | Fp2 | Anodal | Online task- Sternberg WM task performed during stimulation | 10 | 1 mA, 35 cm2 | **CogState battery**- OCLT, *n*-back task (1-back, 2-back)- stimuli are cards  **Sternberg WM task-** decide whether a consonant was presented in a previously shown memory set | Less semantic | Visual | Nonverbal |
|  |  |  |  |  |  |  |  | Offline assessment- CogState battery performed before and after stimulation |  |  |  |  |  |  |
| Teo et al. (2011) [95] | Crossover, sham-controlled, double-blind trial  No. of sessions: 2 (different current intensities) | 12 | 27.23 | tDCS | F3 | Fp2 | Anodal | Online task- performed during stimulation (10–15 min into tDCS and 15–20 min into tDCS) | 20 | 1 mA, 2 mA, 35 cm2 | ***n*-back task (3-back)-** stimuli are letters | Less semantic | Visual | Nonverbal |
| Jeon & Han (2012) [13] | Parallel, sham-controlled trial | RA: 8 LA: 8 RS: 8 LS: 8 | RA: 35.13 LA: 39.50 RS: 37.88 LS: 36.50 | tDCS | F3 | Fp2 | Anodal | Offline assessment- performed before (baseline), immediately after, and two weeks after stimulation | 20 | 1 mA, 35 cm2 | **DSB** | Less semantic | Auditory | Verbal |
|  |  |  |  |  | F4 | Fp1 | Anodal |  |  |  |  |  |  |  |
| Mylius et al. (2012) [96] | Crossover, sham-controlled trial  But different groups for stim. side (left, right) | L: 12 R: 12 | L: 25.1 R: 23.5 | tDCS | F3 | Fp2 | Anodal | Online assessment- performed during stimulation | 20 | 2 mA, 35 cm2 | ***n*-back task (2-back)-** stimuli are numbers | Less semantic | Visual | Nonverbal |
|  |  |  |  |  | Fp2 | F3 | Cathodal |  |  |  |  |  |  |  |
|  |  |  |  |  | F4 | Fp1 | Anodal |  |  |  |  |  |  |  |
|  |  |  |  |  | Fp1 | F4 | Cathodal |  |  |  |  |  |  |  |
| Hoy et al. (2013) [97] | Crossover, sham-controlled, double-blind trial  No. of sessions: 2 (different current intensities) | 18 | 24.71 | tDCS | F3 | Fp2 | Anodal | Offline assessment- performed after stimulation (immediately following (T0), 20 mins (T1) and 40 mins (T2) post-stimulation) | 20 | 1 mA, 2 mA, 35 cm2 | ***n*-back task (2-back, 3-back)-** stimuli are letters | Less semantic | Visual | Nonverbal |
| Meiron & Lavidor (2013) [98] | Parallel, sham-controlled, participant-blind trial | L: 14 R: 16 Sham: 11 | L: 24.86 R: 25.38 Sham: 23.09 | tDCS | F3/AF3 | Cz | Anodal | Offline task- performed after stimulation | 15 | 2 mA, 16 cm2 (active), 35 cm2 (reference) | **Modified verbal *n*-back task (2-back)-** decide if a word is part of a word-pair that appeared two study-displays prior to response-display | Less semantic | Visual | Nonverbal |
|  |  |  |  |  | F4/AF4 | Cz | Anodal |  |  |  |  |  |  |  |
| Nozari & Thompson-Schill (2013) [99] | Crossover, sham-controlled, trial | 24 | 19-30 | tDCS | F3 | F4 | Bilateral | Online assessment- performed during stimulation | 20 | 1.5 mA, 25 cm2 | ***n*-back task (1-back, 2-back, 3-back)-** stimuli are numbers | Less semantic | Visual | Nonverbal |
| Tanoue et al. (2013) [100] | Crossover, sham-controlled, participant-blind trial | 24 | 23.7 | tDCS | Extracephalic | F4 | Cathodal | Offline assessment- performed after stimulation | 10 | 1.5 mA, 35 cm2 | **CDT-** remember colour and location conjunction of stimuli and determine whether or not they match the probe | Less semantic | Visual | Nonverbal |
|  |  |  |  |  | Extracephalic | P4 | Cathodal |  |  |  |  |  |  |  |
| Keeser et al. (2011) [101] | Crossover, sham-controlled, double-blind trial | 10 | 28.89 | tDCS | F3 | Fp2 | Anodal | Offline assessment- performed before (separate day; baseline) and after (not immediately) stimulation | 20 | 2 mA, 35 cm2 | ***n*-back task (1-back, 2-back)-** stimuli are numbers | Less semantic | Visual | Nonverbal |
| Motohashi et al. (2013) [80] | Crossover, sham-controlled, participant-blind trial  But two different groups regarding stim. order  No. of sessions: 4 | Group A: 6 Group B: 6 | 22 | tDCS | F3 | Fp2 | Anodal | Offline assessment- performed before (baseline) and after stimulation | 20 | 1 mA, 35 cm2 | **OCLT** | Less semantic | Visual | Nonverbal |
| Baumert et al. (2020) [23] | Parallel, sham-controlled, participant-blind trial | Anodal: 31 Cathodal: 34 Sham: 31 | 23.04 | tDCS | F3 | Extracephalic (ipsilateral) | Anodal | Offline assessment- performed after stimulation | 20 | 1 mA, 35 cm2 | ***n*-back task (1-back, 2-back, 3-back)-** stimuli are letters | Less semantic | Visual | Nonverbal |
|  |  |  |  |  | Extracephalic (ipsilateral) | F3 | Cathodal |  |  |  |  |  |  |  |
| Gill et al. (2015)- Exp. 1 [102] | Crossover, sham-controlled, double-blind trial | 11 | 21.8 | tDCS | F3 | Fp2 | Anodal | Online task- performed during stimulation | 20 | 2 mA, 25 cm2 | ***n*-back task (1-back, 3-back)-** stimuli are letters | Less semantic | Visual | Nonverbal |
| Gill et al. (2015)- Exp. 2 [102] | Crossover, sham-controlled, double-blind trial | 12 | 19.8 | tDCS | F3 | Fp2 | Anodal | Online task- performed during stimulation | 20 | 2 mA, 25 cm2 | ***n*-back task (1-back, 3-back)-** stimuli are letters | Less semantic | Visual | Nonverbal |
| Wu et al. (2014) [103] | Crossover, sham-controlled trial | 20 | 26 | tDCS | F4 | Extracephalic | Anodal | Offline assessment- performed after stimulation | 15 | 1.5 mA, 25 cm2 | **CBT-** memorise the location of targets and the sequence of their presentation, and then recall them either in the forward manner, as the same sequence of the target presentation, or in the backward manner, as the reverse sequence; conditions: forward-without-interference, backward-without-interference, forward-with-interference, backward-with-interference | Less semantic | Visual | Nonverbal |
| Friehs & Frings (2019a) [104] | Parallel, sham-controlled, participant-blind trial | Active/offline: 21 Active/online: 21 Sham: 21 | n.r. | tDCS | F3 | Extracephalic (ipsilateral) | Anodal | Online assessment- performed during stimulation | 19 | 0.5 mA, 9 cm2 (active), 35 cm2 (reference) | ***n*-back task (3-back)-** stimuli are letters | Less semantic | Visual | Nonverbal |
| Lukasik et al. (2018) [105] | Crossover, sham-controlled, double-blind trial | 33 | 22.6 | tDCS | F7 | Fp2 | Anodal | Online/offline- performed before (baseline; Block 1), during (Block 2), and after (Block 3) stimulation | 10 | 1.5 mA, 48 cm2 (active), 30 cm2 (reference) | ***n*-back task (3-back)-** stimuli are numbers | Less semantic | Visual | Nonverbal |
|  |  |  |  |  | Fp2 | F7 | Cathodal |  |  |  |  |  |  |  |
| Mashal & Metzuyanim-Gorelick (2019) [106] | Parallel, sham-controlled, participant-blind trial  No. of sessions: 6 | Active: 16 Sham: 9 | Active: 31.44 Sham: 29.62 | tDCS | F3 | F4 | Bilateral | Offline assessment- performed before (baseline; T1) and after (immediately and one month after; T2 and T3) stimulation | 20 | 2 mA, 35 cm2 | ***n*-back task (1-back, 2-back, 3-back, 4-back)-** stimuli are shapes | Less semantic | Visual | Nonverbal |
| Naka et al. (2018) [107] | Parallel, sham-controlled, participant-blind trial | Active: 10 Sham: 10 | 22.7 | HD-tDCS | F3 | C3, F7, FP1, Fz | Anodal | Online/offline assessment- performed before, during, and after stimulation | 16 | 1.5 mA, approx. 4 cm2 per electrode | ***n*-back task (3-back)-** stimuli are letters (visual and auditory) | Less semantic | Visual & auditory | Nonverbal |
| Nikolin et al. (2017) [108] | Parallel, sham-controlled, participant-blind trial | Active: 10 Sham: 10 | Active: 22.3 Sham: 23.3 | tDCS | F3 | F4 | Bilateral | Online/offline assessment- performed before (baseline), during, and after stimulation | 15 | 2 mA, 16 cm2 | ***n*-back task (3-back)-** stimuli are letters | Less semantic | Visual | Nonverbal |

| **Reference (Year)** | **Study Design** | **Participant N** | **Mean Age in Years** | **Stimulation Modality** | **Anode** | **Cathode** | **Polarity of Stimulating Electrode** | **Online/Offline** | **Duration** | **Current Intensity, Electrode Size** | **Experimental Task Description** | **Semantic/**  **Less semantic** | **Visual/**  **Auditory** | **Verbal/**  **Nonverbal** |
| --- | --- | --- | --- | --- | --- | --- | --- | --- | --- | --- | --- | --- | --- | --- |
| ***Working Memory*** | | | | | | | | | | | | | | |
| Röhner et al. (2018) [109] | Crossover, sham-controlled, participant-blind trial | 30 | 26.2 | tDCS | F3 | Extracephalic (ipsilateral) | Anodal | Online/offline assessment- performed before, during, and after stimulation | 15 | 1 mA, 35 cm2 | ***n*-back task (2-back)-** stimuli are letters | Less semantic | Visual | Nonverbal |
| Talsma et al. (2017) [110] | Parallel, sham-controlled, double-blind trial  No. of sessions: 3 | Active: 15 Sham: 15 | Active: 21.9 Sham: 22.1 | tDCS | F3 | Fp2 | Anodal | Online/offline assessment- performed before (baseline), during (three sessions), and after stimulation | 20 | 1 mA, 35 cm2 | ***n*-back task (3-back, 4-back)-** stimuli are letters | Less semantic | Visual | Nonverbal |
| Trumbo et al. (2016)- Exp. 1 [111] | Parallel, sham-controlled, double-blind trial | L: 12 R: 12 Sham: 12 | L: 18.75 R: 21.42 Sham: 20 | tDCS | F3 | Extracephalic | Anodal | Offline assessment- verbal 3-back performed before (baseline) and after stimulation  Online/offline assessment/task- spatial 3-back performed before (baseline), during (training: TR 2-5), and after stimulation | 30 | 2 mA, 25 cm2 | ***n*-back task (3-back)-** stimuli are location of filled boxes (spatial)  ***n*-back task (3-back)-** stimuli are letters (verbal) | Less semantic | Visual | Nonverbal |
|  |  |  |  |  | F4 | Extracephalic | Anodal |  |  |  |  |  |  |  |
| Trumbo et al. (2016)- Exp. 2 [111] | Parallel, sham-controlled, double-blind trial | L: 12 R: 12 Sham: 12 | L: 22.25 R: 18.92 Sham: 21.5 | tDCS | F3 | Extracephalic | Anodal | Offline assessment- spatial 3-back performed before (baseline) and after stimulation  Online/offline assessment/task- verbal 3-back performed before (baseline), during (training: TR 2-5), and after stimulation | 30 | 2 mA, 25 cm2 | ***n*-back task (3-back)-** stimuli are location of filled boxes (spatial)  ***n*-back task (3-back)-** stimuli are letters (verbal) | Less semantic | Visual | Nonverbal |
|  |  |  |  |  | F4 | Extracephalic | Anodal |  |  |  |  |  |  |  |
| Wang et al. (2018) [112] | Parallel, sham-controlled trial | Anodal: 10 Cathodal: 10 Sham: 10 | 23.2 | tDCS | F4 | Fp1 | Anodal | Offline assessment- performed before (baseline; different day) and after stimulation | 20 | 1.5 mA, 25 cm2 | ***n*-back task (1-back, 2-back)-** stimuli are location of shapes; easy and difficult conditions | Less semantic | Visual | Nonverbal |
|  |  |  |  |  | Fp1 | F4 | Cathodal |  |  |  |  |  |  |  |
| Keshvari et al. (2013) [113] | Crossover, sham-controlled, participant-blind trial  But different groups for polarity (LA/RC, LC/RA) | LA/RC: 30 LC/RA: 30 | LA/RC: 22.3 LC/RA: 21.2 | tDCS | F3 | F4 | Bilateral | Offline assessment- performed before and after stimulation | 20 | 2 mA, 25 cm2 | ***n*-back task (2-back)-** stimuli are figures | Less semantic | Visual | Nonverbal |
|  |  |  |  |  | F4 | F3 | Bilateral |  |  |  |  |  |  |  |
| Lally et al. (2013) [114] | Parallel, sham-controlled, double-blind trial  No. of sessions: 2 | Active: 10 Sham: 11 | 23.09 | tDCS | F3 | Extracephalic | Anodal | Online/offline assessment- performed before (baseline), during, and after stimulation (Day 1); performed during and after stimulation (Day 2) | 10 | 1 mA, 35 cm2 | ***n*-back task (3-back)-** stimuli are letters | Less semantic | Visual | Nonverbal |
| Giglia et al. (2014) [115] | Crossover, sham-controlled, participant-blind trial | 10 | 27 | tDCS | F3 | Fp2 | Anodal | Online assessment- performed before (baseline) and during stimulation | 10 | 1 mA, 35 cm2 | **Delayed recognition task-** indicate whether location of stimuli corresponds to those of previously shown stimuli | Less semantic | Visual | Nonverbal |
|  |  |  |  |  | F4 | Fp1 | Anodal |  |  |  |  |  |  |  |
| Lu et al. (2021) [65] | Parallel, sham-controlled, participant-blind trial  No. of sessions: 9 | Active: 22 Sham: 21 | Active: 20.73 Sham: 21.10 | HD-tDCS | F3 | AF3, F1, F5, FC3 | Anodal | Offline assessment- performed before and after stimulation  *Note:* Phase 1 (n-back), Phase 2 (9 tDCS sessions were undertaken across 18 days, with n-back completed on the day after the 5th HD-tDCS session), Phase 3 (n-back); the three phases are all on different days | 20 | 1.5 mA, electrode size n.r. | ***n*-back task (2-back)-** stimuli are colours | Less semantic | Visual | Nonverbal |
| Hasan et al. (2023) [66] | Parallel, sham-controlled, participant-blind trial  No. of sessions: 5 | Active: 19 Sham: 21 | 22 | tDCS | F3 | F4 | Bilateral | Offline assessment- performed before and after stimulation | 30 | 2 mA, 20 cm2 | ***n*-back task (3-back)-** stimuli are letters | Less semantic | Visual | Nonverbal |
| Pupíková et al. (2021) [116] | Crossover, sham-controlled, double-blind trial | 27 | 27 | tDCS | MNI: 44 40 -10 (right middle frontal gyrus [rMFG]) | MNI: 30 -55 52 (right posterior parietal cortex [rPPC]) | Bilateral | Offline assessment- VOMT performed before and after stimulation  Online task- WM task performed during stimulation | 20 | 2 mA, 25 cm2 | **VOMT-** presented with paired images of common objects, with the second image of each pair either being same or different from the first image (different object identity or object orientation- conventional view, unconventional view); respond whether or not the objects are the same, regardless of spatial orientation  **WM task with faces/outdoor scenes-** upon presentation of images of faces and scenes, attend to one category and ignore irrelevant distractor category; after a retention period, respond whether or not a probe was present in the foregoing series | Semantic (picture) | Visual | Nonverbal |

| **Reference (Year)** | **Study Design** | **Participant N** | **Mean Age in Years** | **Stimulation Modality** | **Anode** | **Cathode** | **Polarity of Stimulating Electrode** | **Online/Offline** | **Duration** | **Current Intensity, Electrode Size** | **Experimental Task Description** | **Semantic/**  **Less semantic** | **Visual/**  **Auditory** | **Verbal/**  **Nonverbal** |
| --- | --- | --- | --- | --- | --- | --- | --- | --- | --- | --- | --- | --- | --- | --- |
| ***Working Memory*** | | | | | | | | | | | | | | |
| Martin et al. (2023) [117] | Crossover, sham-controlled trial  But different groups for stim. region (bi-f, f, par) | Bi: 31 F: 31 P: 31 | Bi: 22.8 F: 23.5 P: 22 | tDCS | F3 | F8 | Bilateral | Online assessment- performed during stimulation | 20 | 2 mA, 35 cm2 | ***n*-back task (1-back, 2-back)-** stimuli are letters | Less semantic | Visual | Nonverbal |
|  |  |  |  |  | F3 | Extracephalic | Anodal |  |  |  |  |  |  |  |
|  |  |  |  |  | P3 | Extracephalic | Anodal |  |  |  |  |  |  |  |
| Nikolin et al. (2018)- Exp. 1 [118] | Parallel, sham-controlled, participant-blind trial | Active: 20 Sham: 20 | Active: 25.1 Sham: 22.9 | tDCS | F3 | F4 | Bilateral | Online/offline assessment- performed before, during, and after stimulation | 15 | 2 mA, 16 cm2 | ***n*-back task (3-back)-** stimuli are letters | Less semantic | Visual | Nonverbal |
| Nikolin et al. (2018)- Exp. 2 [118] | Parallel, sham-controlled, participant-blind trial | Active: 20 Sham: 20 | Active: 22.8 Sham: 21.9 | tDCS | F3 | F4 | Bilateral | Online/offline assessment- performed before, during, and after stimulation | 15 | 1 mA, 16 cm2 | ***n*-back task (3-back)-** stimuli are letters | Less semantic | Visual | Nonverbal |
| Ramaraju et al. (2020) [119] | Crossover, sham-controlled, double-blind trial | 20 | 30 | tDCS | F3 | Fp2 | Anodal | Offline assessment- performed after stimulation | 15 | 1.5 mA, 35 cm2 | ***n*-back task (2-back)-** stimuli are letters and shapes | Less semantic | Visual | Nonverbal |
| Živanović et al. (2021)- Exp. 1 [120] | Crossover, sham-controlled, participant-blind trial | 21 | 26.76 | tDCS | F3 | Extracephalic | Anodal | Offline assessment- performed after stimulation | 20 | 1.8 mA, 25 cm2 | ***n*-back task (3-back)-** stimuli are letters (verbal) and position of squares (spatial) | Less semantic | Visual | Nonverbal |
|  |  |  |  |  | P3 | Extracephalic | Anodal |  |  |  |  |  |  |  |
| Živanović et al. (2021)- Exp. 2 [120] | Crossover, sham-controlled, participant-blind trial | 21 | 26.43 | tDCS | F4 | Extracephalic | Anodal | Offline assessment- performed after stimulation | 20 | 1.8 mA, 25 cm2 | ***n*-back task (3-back)-** stimuli are letters (verbal) and position of squares (spatial) | Less semantic | Visual | Nonverbal |
|  |  |  |  |  | P4 | Extracephalic | Anodal |  |  |  |  |  |  |  |
| Živanović et al. (2021)- Exp. 3 [120] | Crossover, sham-controlled, participant-blind trial | 21 | 24.9 | tDCS | F3 | Extracephalic | Anodal | Online assessment- performed during stimulation | 20 | 1.5 mA, 25 cm2 | ***n*-back task (3-back)-** stimuli are letters (verbal) and position of squares (spatial) | Less semantic | Visual | Nonverbal |
|  |  |  |  |  | P3 | Extracephalic | Anodal |  |  |  |  |  |  |  |
| Papazova et al. (2020)- Exp. 1 [121] | Crossover, sham-controlled, double-blind trial | 16 | 31.06 | tDCS | F3 | Extracephalic | Anodal | Online assessment- performed during stimulation | 21 | 1 mA, 35 cm2 | ***n*-back task (1-back, 2-back, 3-back)-** stimuli are letters | Less semantic | Visual | Nonverbal |
| Papazova et al. (2020)- Exp. 2 [121] | Crossover, sham-controlled, double-blind trial | 16 | 32.13 | tDCS | F3 | Extracephalic | Anodal | Online assessment- performed during stimulation | 21 | 2 mA, 35 cm2 | ***n*-back task (1-back, 2-back, 3-back)-** stimuli are letters | Less semantic | Visual | Nonverbal |
| Zhu et al. (2020) [122] | Parallel, sham-controlled, participant-blind trial | PPC: 17 IFG: 17 Sham: 17 | 18.9 | tDCS | P3 | Extracephalic | Anodal | Offline assessment- performed before and after stimulation | 20 | 1.5 mA, 35 cm2 | ***n*-back task (3-back)-** stimuli are letters (visual and auditory) | Less semantic | Visual & auditory | Nonverbal |
|  |  |  |  |  | Crossing point between F7-Cz and T3-Fz | Extracephalic | Anodal |  |  |  |  |  |  |  |
| Dumont et al. (2021) [123] | Crossover, sham-controlled trial | 47 | 24.15 | tDCS | P3 | Extracephalic | Anodal | Online assessment- performed during stimulation | 20 | 2 mA, 9 cm2 (active), 25 cm2 (reference) | **Visuospatial WM task (visual array probe recognition)-** memorise the position of coloured squares to be recalled later with a probe; load 2 and load 6 **Verbal WM task (letter probe recognition task)-** memorise horizontal sequences of 2 or 6 letters; when a probe letter appears in one of the 2 or 6 possible serial positions, decide if the letter had been in the memory list and if it had appeared in the same position as in the probe array | Less semantic | Visual | Nonverbal |
|  |  |  |  |  | Extracephalic | P3 | Cathodal |  |  |  |  |  |  |  |
| Zhu et al. (2022) [124] | Parallel, sham-controlled, participant-blind trial  No. of sessions: 3 | Active: 18 Sham: 18 | 19 | tDCS | P4 | Extracephalic | Anodal | Offline assessment- performed before and after stimulation | 10 (single tDCS), 20 (repeated tDCS with 5 min and 20 min intervals) | 1.5 mA, 35 cm2 | ***n*-back task (2-back, 3-back)-** stimuli are positions of squares | Less semantic | Visual | Nonverbal |
| Westwood & Romani (2018)- Exp. 2 [71] | Crossover, sham-controlled, participant-blind trial  But different groups for recent-probe and semantic-associated probe | Recent: 20 Semantic: 19 | Recent: 20 Semantic: 19 | tDCS | F7 | Fp2 | Anodal | Online assessment- performed during stimulation | 25 | 1.5 mA, 25 cm2 (active), 35 cm2 (reference) | **Probe tasks-** decide whether or not a probe has appeared in a list or words presented earlier; **recent-probe task** and a **semantic-associated probe task** | Semantic (word) | Visual | Nonverbal |
| Marko & Riečanský (2021) [73] | Parallel, sham-controlled, double-blind trial | PFC: 40 TPC: 40 Sham: 41 | 23.1 | tDCS | Between F3 and AF3 | Between T7 and P5 | Bilateral | Online/offline assessment- performed before, during, and after stimulation | 25 | 2 mA, 35 cm2 | **Interference DS task-** listen to a series of digits presented either in high pitch (remember) or low pitch (ignore), and then report remembered items in reverse order | Less semantic | Auditory | Nonverbal |
|  |  |  |  |  | Between T8 and P6 | Between T7 and P5 | Bilateral |  |  |  |  |  |  |  |

| **Reference (Year)** | **Study Design** | **Participant N** | | **Mean Age in Years** | | **Stimulation Modality** | | **Anode** | | **Cathode** | **Polarity of Stimulating Electrode** | **Online/Offline** | **Duration** | **Current Intensity, Electrode Size** | **Experimental Task Description** | **Semantic/**  **Less semantic** | **Visual/**  **Auditory** | **Verbal/**  **Nonverbal** |
| --- | --- | --- | --- | --- | --- | --- | --- | --- | --- | --- | --- | --- | --- | --- | --- | --- | --- | --- |
| ***Learning & Memory*** | | | | | | | | | | | | | | | | | | |
| Elmer et al. (2009) [79] | Crossover, sham-controlled trial  But stim. side is between-subjects | L: 10 R: 10 | | 22.3 | | tDCS | | F3 | | Extracephalic | Anodal | Offline assessment- LTR performed after stimulation  Online assessment- STL performed during stimulation | 5 | 1.5 mA, 28 cm2 (active), 100 cm2 (reference) | **Verbal learning paradigm:** for STL test, encode and immediately retrieve auditory presented single nouns; then perform a nonverbal recognition test (deciding whether meaningless figures has been presented before) to avoid active memory strategies until later retrieval; after a short pause in which a silent cartoon is presented, complete the LTR test, whereby single-nouns from before have to be verbally retrieved | Semantic (word) | Visual & auditory | Verbal |
|  |  |  |  |  |  |  |  | Extracephalic | | F3 | Cathodal |  |  |  |  |  |  |  |
|  |  |  |  |  |  |  |  | F4 | | Extracephalic | Anodal |  |  |  |  |  |  |  |
|  |  |  |  |  |  |  |  | Extracephalic | | F4 | Cathodal |  |  |  |  |  |  |  |
| Ambrus et al. (2011) [125] | Parallel, sham-controlled, participant-blind trial | RA: 12 LA: 12 RC: 12 Sham: 12 | | 19-40 | | tDCS | | 9 cm to the right and 7 cm anterior relative to the Cz (presumably F4) | | Cz | Anodal | Offline assessment- testing phase performed after stimulation | 10 | 1 mA, 35 cm2 | **Prototype distortion task (categorisation learning)-** during the training phase, presented with low- and high distortions of a prototype dot-pattern, and in the testing phase, decide whether the consecutive high- and low-distortion versions of the prototypes (category members) or random patterns (non-members) that are presented belong to the category established in the training phase | Less semantic | Visual | Nonverbal |
|  |  |  |  |  |  |  |  | 9 cm to the left and 7 cm anterior relative to the Cz (presumably F3) | | Cz | Anodal |  |  |  |  |  |  |  |
|  |  |  |  |  |  |  |  | Cz | | 9 cm to the right and 7 cm anterior relative to the Cz (presumably F4) | Cathodal |  |  |  |  |  |  |  |
| Hammer et al. (2011) [126] | Crossover, sham-controlled, participant-blind trial | | Anodal: 18 Cathodal: 18 | | Anodal: 23.3 Cathodal: 23 | | tDCS | | F3 | Fp2 | Anodal | Online/offline task- learning and recognition phases performed during and after stimulation | 30 | 1 mA, 35 cm2 | **Recognition memory task-** decide whether a presented word is a target word or not (recognition phase), in the context where words have been acquired (learning phase during errorless learning (first three letters of the word are introduced directly followed by the target word in a word-fragment-completion task) and errorful learning (guess several words to complete the word-fragment before the target word is revealed) | Semantic (word) | Visual | Verbal |
|  |  |  |  |  |  |  |  |  | Fp2 | F3 | Cathodal |  |  |  |  |  |  |  |
| Javadi & Walsh (2012)- Exp. 1 [127] | Crossover, sham-controlled, participant-blind trial | | 16 | | 22.46 for all 32 Ps (i.e., Exp. 1 & Exp. 2) | | tDCS | | F3 | Fp2 | Anodal | Online/offline assessment- encoding phase performed during stimulation, recognition phase performed after stimulation | 20 | 1 mA, 12.25 cm2 (active), 30.25 cm2 (reference) | **Word memorisation task-** during the encoding phase, quickly respond to the number of syllables of words, and imagine the words to memorise them; during the retention interval, watch sketches of a television series; during the recognition (retrieval) phase, select the word (among a pair of words) that was seen during the encoding phase | Semantic (word) | Visual | Verbal |
|  |  |  |  |  |  |  |  |  | Fp2 | F3 | Cathodal |  |  |  |  |  |  |  |
| Javadi & Walsh (2012)- Exp. 2 [127] | Crossover, sham-controlled, participant-blind trial | | 16 | | 22.46 for all 32 Ps (i.e., Exp. 1 & Exp. 2) | | tDCS | | F3 | Fp2 | Anodal | Online assessment- encoding phase performed before stimulation, recognition phase performed during stimulation | 20 | 1 mA, 12.25 cm2 (active), 30.25 cm2 (reference) | **Word memorisation task-** during the encoding phase, quickly respond to the number of syllables of words, and imagine the words to memorise them; during the retention interval, watch sketches of a television series; during the recognition (retrieval) phase, select the word (among a pair of words) that was seen during the encoding phase | Semantic (word) | Visual | Verbal |
|  |  |  |  |  |  |  |  |  | Fp2 | F3 | Cathodal |  |  |  |  |  |  |  |
| Asthana et al. (2013) [128] | Parallel, sham-controlled trial | | Anodal: 16 Cathodal: 18 Sham: 15 | | Anodal: 22.18 Cathodal: 23.11 Sham: 22.46 | | tDCS | | F3 | Extracephalic (ipsilateral) | Anodal | Offline assessment- habituation and acquisition phases performed before stimulation, extinction phase performed after stimulation | 12 | 1 mA, 35 cm2 | **Auditory fear-conditioning paradigm-** **habituation** (day 1: coloured squares (blue and yellow) are presented on the monitor to reach a stable response to stimuli), **acquisition** (day 1: coloured squares are presented, and one is paired with the scream), **extinction** (day 2: extinction training, whereby conditioned stimuli are repeatedly presented in the absence of unconditioned stimuli) | Less semantic | Visual & auditory | Nonverbal |
|  |  |  |  |  |  |  |  |  | Extracephalic (ipsilateral) | F3 | Cathodal |  |  |  |  |  |  |  |
| Iuculano & Cohen Kadosh (2013) [19] | Parallel, sham-controlled, participant-blind trial  No. of sessions: 6 | | 19 (subgroup n n.r.) | | 20-31 | | tDCS | | F3 | F4 | Bilateral | Online task- performed during stimulation | 20 | 1 mA, 3 cm2 | **Learning task-** learn artificial digits for a Stroop task by referring to meaningless symbols as though they represent various magnitudes; when two symbols are displayed, choose which has the larger magnitude | Less semantic | Visual | Nonverbal |
|  |  |  |  |  |  |  |  |  | P3 | P4 | Bilateral |  |  |  |  |  |  |  |
| Javadi & Cheng (2013) [129] | Crossover, sham-controlled trial  But two different groups (recon. and control) | | Recon.: 15 Control: 15 | | 22.58 | | tDCS | | F3 | Fp2 | Anodal | **Recognition group:**  Online task- first old-new recognition task performed during stimulation (stimulation session)  Offline assessment- second old-new recognition task performed after stimulation (recognition session)  **Control group:**  Offline assessment- old-new recognition task performed after stimulation | 20 | 1.5 mA, 12.25 cm2 (active), 30.25 cm2 (reference) | **LTM task-** during encoding session, imagine presented words in order to memorise them, and then perform an old–new recognition task; a few hours later, perform the old–new word recognition task again, and also rate confidence of responses | Semantic (word) | Visual | Verbal |
|  |  |  |  |  |  |  |  |  | Fp2 | F3 | Cathodal |  |  |  |  |  |  |  |

| **Reference (Year)** | **Study Design** | **Participant N** | **Mean Age in Years** | **Stimulation Modality** | **Anode** | **Cathode** | **Polarity of Stimulating Electrode** | **Online/Offline** | **Duration** | **Current Intensity, Electrode Size** | **Experimental Task Description** | **Semantic/**  **Less semantic** | **Visual/**  **Auditory** | **Verbal/**  **Nonverbal** |
| --- | --- | --- | --- | --- | --- | --- | --- | --- | --- | --- | --- | --- | --- | --- |
| ***Learning & Memory*** | | | | | | | | | | | | | | |
| Kongthong et al. (2013) [130] | Crossover, sham-controlled trial | 14 | 24.79 | tDCS | T6 | F3 | Bilateral | Offline assessment- performed after stimulation | 20 | 1 mA, 25 cm2 | **Masked-face priming paradigm-** after being presented with a subliminal prime preceding a target stimulus, determine whether the target face is a famous face or not; prime and target pair are either the same person's face (congruent) or different person's faces (incongruent), and are always both famous or both non-famous faces; there are masked prime trials and unmasked prime trials | Semantic (picture) | Visual | Nonverbal |
| Santiesteban et al. (2012) [131] | Parallel, sham-controlled trial | Anodal: 17 Cathodal: 17 Sham: 15 | 26.5 | tDCS | CP6 | Vertex | Anodal | Offline assessment- performed after stimulation | 20 | 1 mA, 35 cm2 | **Surprise recognition memory test-** for judgements made in the self-referential task (main task, not relevant here); presented with a judgement statement and asked to rate confidence of having seen it before on a scale of 1 (no) to 6 (yes); for items rated from 4 to 6, further asked to rate confidence that the statement is in reference to themselves (participants) or to Lady Gaga on a scale of 1 (self) to 6 (Lady Gaga) | Semantic (word) | Visual | Verbal |
|  |  |  |  |  | Vertex | CP6 | Cathodal |  |  |  |  |  |  |  |
| Beharelle et al. (2015) [132] | Parallel, sham-controlled, double-blind trial | Anodal: 26 Cathodal: 27 Sham: 26 | Anodal: 23.35 Cathodal: 22.88 Sham: 22.96 | tDCS | Right frontopolar cortex (FPC) region (MNI peak: x = 27, y = 57, z = 6) | Vertex | Anodal | Online assessment- performed before (baseline) and during stimulation | n.r. (only that 3 mins of stim. passed before task started) | 1 mA, 25 cm2 (active), 100 cm2 (reference) | **Reward-learning task (bandit task)-** select among three virtual slot machines whose payout values drift independently and randomly across trials, whereby the time-varying monetary rewards require participants to learn continuously about the slot machines to maximise their monetary payoffs | Less semantic | Visual | Nonverbal |
|  |  |  |  |  | Vertex | Right FPC region (MNI peak: x = 27, y = 57, z = 6) | Cathodal |  |  |  |  |  |  |  |
| Martin et al. (2017) [133] | Crossover, sham-controlled, double-blind trial  But different groups for anodal and cathodal stim. | Anodal: 20 Cathodal: 20 | Anodal: 22.9 Cathodal: 24 | HD-tDCS | 15% of the distance from the Fz towards the FpZ | Symmetrically around the centre electrode | Anodal | Online assessment- performed during stimulation | 20 | 1 mA, diameter: 2.5 cm (centre electrode), diameter inner/outer: 9.2/11.5 cm (ring-shaped reference electrode) | **Self-referential memory task-** presented with mental attribution words and distractor words, decide whether mental attribution was seen before in the RMET task completed earlier (main task, not relevant here) on a scale of 1 (yes) to 4 (no)  **Source memory task-** if a response is made that the mental attribution was seen before, answered a subsequent question: "Was it on a male or a female face?", on a scale of 1 (male) to 4 (female) | Semantic (word) | Visual | Verbal |
|  |  |  |  |  | Symmetrically around the centre electrode | 15% of the distance from the Fz towards the FpZ | Cathodal |  |  |  |  |  |  |  |

| **Reference (Year)** | **Study Design** | **Participant N** | **Mean Age in Years** | **Stimulation Modality** | | **Anode** | | **Cathode** | **Polarity of Stimulating Electrode** | **Online/Offline** | | **Duration** | | **Current Intensity, Electrode Size** | | **Experimental Task Description** | | **Semantic/**  **Less semantic** | | **Visual/**  **Auditory** | | **Verbal/**  **Nonverbal** |
| --- | --- | --- | --- | --- | --- | --- | --- | --- | --- | --- | --- | --- | --- | --- | --- | --- | --- | --- | --- | --- | --- | --- |
| ***Learning & Memory*** | | | | | | | | | | | | | | | | | | | | | | |
| Stramaccia et al. (2017)- Exp. 2 [46] | Parallel, sham-controlled, participant-blind trial | Anodal: 24 Cathodal: 24 Sham: 24 | Anodal: 23.96 Cathodal: 23.33 Sham: 23.42 | tDCS | | FC4 | | Fp1 | Anodal | Online/offline assessment- study phase performed before stimulation (no outcome measured), practice phase performed during stimulation, test phase performed after stimulation | | 20 | | 1.5 mA, 16 cm2 | | **RPP-** in study phase, memorise category-exemplar word pairs, by relating each exemplar to its category; in practice phase, when the category and the first three letters of each exemplar are provided, provide the name of the specific exemplar associated to the particular cue in full; in test phase, shown the category plus the first letter of an exemplar only, with instructions being same as that of the practice phase | | Semantic (word) | | Visual | | Verbal |
|  |  |  |  |  |  | Fp1 | | FC4 | Cathodal |  |  |  |  |  |  |  |  |  |  |  |  |  |
| Orth et al. (2023) [89] | Parallel, sham-controlled, double-blind trial | L: 64 R: 64 Sham: 126 | L: 23.92 R: 22.7 Sham: 23.31 | tDCS | | F3 | | Fp2 | Anodal | Online/offline assessment- encoding during stimulation, free recall and recognition performed after stimulation | | 20 | | 1 mA, 35 cm2 | | **Incidental learning task (episodic memory)-** first encode nouns (randomly assigned to shallow, deep, or emotional word groups), then make decisions according to group (shallow: indicate whether the word contains an A or whether the word has exactly three syllables; deep: indicate whether the word is “animate” or denoted something “edible”; emotional: indicate whether the word elicits “joy” or “fear”); in retrieval phase, retrieve learned words in the form of free recall and recognition (targets and non-targets) | | Semantic (word) | | Visual | | Verbal |
|  |  |  |  |  | | F4 | | Fp1 | Anodal |  |  |  |  |  |  |  |  |  |  |  |  |  |
| Habich et al. (2020) [134] | Crossover, sham-controlled, double-blind trial | 33 | 24.5 | tDCS | F3 | | Fp2 | | Anodal | Online/offline assessment- encoding phase during stimulation; retrieval performed after stimulation | 20 | | 1 mA, 35 cm2 | | **Verbal episodic memory task-** after encoding words, engage in immediate recall, followed by free delayed recall and recognition | | Semantic (word) | | Visual | | Verbal | |
| Huang et al. (2021) [135] | Parallel, sham-controlled, double-blind trial | Anodal: 13 Cathodal: 14 Sham: 13 | 19 | HD-tDCS | FpZ | | Fz, Fp1, Fp2 | | Anodal | Online assessment- learning before stimulation, recall during stimulation | 10.5 | | 1 mA, 1 cm radius circular electrodes | | **Word association task (episodic/associative memory retrieval)-** during Visit 1, learn Swahili-English word-associations (during test phase provide the correct English translation when Swahili word is presented), and during Visit 2 (a few days after; relevant to us), recall what was previously learned | | Semantic (word) | | Visual | | Verbal | |
|  |  |  |  |  | Fz, Fp1, Fp2 | | FpZ | | Cathodal |  |  |  |  |  |  |  |  |  |  |  |  |  |
| Flöel et al. (2008) [136] | Crossover, sham-controlled, double-blind trial | 19 | 25.6 | tDCS | CP5 | | Fp2 | | Anodal | Online and offline assessment- five blocks of language learning task performed during stimulation; lexical knowledge test performed immediately after stimulation, then again one week later | 20 | | 1 mA, 35 cm2 | | **Language learning paradigm:  Language learning-** indicate whether a picture (object)-pseudoword pairing is correct or not **Lexical knowledge test-** indicate whether the object name-pseudoword pairing is correct or not, testing the ability to explicitly translate the novel words into German (native language); immediate and delayed recall | | Semantic (picture) | | Visual (pictures), auditory (pseudowords) | | Verbal | |
|  |  |  |  |  | Fp2 | | CP5 | | Cathodal |  |  |  |  |  |  |  |  |  |  |  |  |  |

| **Reference (Year)** | **Study Design** | **Participant N** | **Mean Age in Years** | **Stimulation Modality** | **Anode** | **Cathode** | **Polarity of Stimulating Electrode** | **Online/Offline** | **Duration** | **Current Intensity, Electrode Size** | **Experimental Task Description** | **Semantic/**  **Less semantic** | **Visual/**  **Auditory** | **Verbal/**  **Nonverbal** |
| --- | --- | --- | --- | --- | --- | --- | --- | --- | --- | --- | --- | --- | --- | --- |
| ***Language*** |  |  |  |  |  |  |  |  |  |  |  |  |  |  |
| Ross et al. (2010) [137] | Crossover, sham-controlled trial | 15 | 25.6 | tDCS | T3 | Extracephalic | Anodal | Online assessment- performed during stimulation | 15 | 1.5 mA, 35 cm2 | **Picture naming-** famous individuals and landmarks | Semantic (picture) | Visual | Verbal |
|  |  |  |  |  | T4 | Extracephalic | Anodal |  |  |  |  |  |  |  |
| Sparing et al. (2008) [138] | Crossover, sham-controlled, double-blind trial | 15 | 26.9 | tDCS | CP5 | Cz | Anodal | Online/offline assessment- performed before, during, and after (0, 5, 10 mins after) stimulation | 7 | 2 mA, 35 cm2 | **Picture naming-** black-and-white line drawings of everyday objects | Semantic (picture) | Visual | Verbal |
|  |  |  |  |  | Cz | CP5 | Cathodal |  |  |  |  |  |  |  |
|  |  |  |  |  | CP6 | Cz | Anodal |  |  |  |  |  |  |  |
| de Vries et al. (2010) [139] | Parallel, sham-controlled trial | Active: 22 Sham: 22 | 22.6 | tDCS | Left BA 44/45 (Broca's area) | Fp2 | Anodal | Online task- implicit acquisition task performed during stimulation  Offline assessment- classification task performed after stimulation | 20 | 1 mA, 35 cm2 (active), 100 cm2 (reference) | **AGL experiment**:  **Implicit acquisition task-** recall a grammatical string of 5-12 letters by typing **Classification task-** classify novel strings as grammatical or nongrammatical; prior to this task, told that the strings presented during the acquisition task were all grammatical | Less semantic | Visual | Nonverbal |
| Fertonani et al. (2010)- Exp. 1 [140] | Crossover, sham-controlled, participant-blind trial | 12 | 24.1 | tDCS | 8 cm frontal and 6 cm lateral with respect to Cz (presumably F3) | Extracephalic | Anodal | Offline assessment- performed after stimulation | 8 | 2 mA, 35 cm2 | **Picture naming-** actions and objects | Semantic (picture) | Visual | Nonverbal |
|  |  |  |  |  | Extracephalic | 8 cm frontal and 6 cm lateral with respect to Cz (presumably F3) | Cathodal |  |  |  |  |  |  |  |
| Fertonani et al. (2010)- Exp. 2 [140] | Crossover, sham-controlled participant-blind trial | 12 | 21.8 | tDCS | 8 cm frontal and 6 cm lateral with respect to Cz (presumably F3) | Extracephalic | Anodal | Offline assessment- performed after stimulation | 10 | 2 mA, 35 cm2 | **Picture naming-** actions and objects | Semantic (picture) | Visual | Verbal |
|  |  |  |  |  | Extracephalic | 8 cm frontal and 6 cm lateral with respect to Cz (presumably F3) | Cathodal |  |  |  |  |  |  |  |
| Jeon & Han (2012) [13] | Parallel, sham-controlled trial | RA: 8 LA: 8 RS: 8 LS: 8 | RA: 35.13 LA: 39.50 RS: 37.88 LS: 36.50 | tDCS | F3 | Fp2 | Anodal | Offline assessment- performed before (baseline), immediately after, and two weeks after stimulation | 20 | 1 mA, 35 cm2 | **Parallel short forms of K-BNT-** name line drawings of common objects | Semantic (picture) | Visual | Nonverbal stimuli verbal response |
|  |  |  |  |  | F4 | Fp1 | Anodal |  |  |  |  |  |  |  |
| Brückner & Kammer (2017) [82] | Parallel, sham-controlled trial | Anodal: 20 Cathodal: 20 Sham: 20 | Anodal: 21.5 Cathodal: 23.7 Sham: 22.9 | tDCS | CP5 | Extracephalic (ipsilateral) | Anodal | Offline assessment- performed after stimulation | 15 | 1 mA, 35 cm2 | **Lexical decision task** | Semantic (word) | Visual | Verbal |
|  |  |  |  |  | Extracephalic (ipsilateral) | CP5 | Cathodal |  |  |  |  |  |  |  |

| **Reference (Year)** | **Study Design** | **Participant N** | **Mean Age in Years** | **Stimulation Modality** | **Anode** | **Cathode** | | **Polarity of Stimulating Electrode** | | **Online/Offline** | | **Duration** | | **Current Intensity, Electrode Size** | | **Experimental Task Description** | | **Semantic/**  **Less semantic** | | **Visual/**  **Auditory** | | **Verbal/**  **Nonverbal** |
| --- | --- | --- | --- | --- | --- | --- | --- | --- | --- | --- | --- | --- | --- | --- | --- | --- | --- | --- | --- | --- | --- | --- |
| ***Language*** |  |  |  |  |  |  | |  | |  | |  | |  | |  | |  | |  | |  |
| Choi & Perrachione (2019) [141] | Parallel, sham-controlled, participant-blind trial | Anodal: 20 Cathodal: 20 Sham: 20 | 20.4 | HD-tDCS | T7, TP7 | C3, CP3, PO7, F7 | | Anodal | | Online assessment- performed during stimulation | | Approx. 13 | | 2 mA, electrode size n.r. | | **Word identification task-** listen to a phase and indicate whether "boot" or "boat" was heard; factors are talker variability (single-talker vs. mixed-talker) and speech context (isolated words vs. connected speech) | | Semantic (word) | | Auditory | | Verbal |
|  |  |  |  |  | C3, CP3, PO7, F7 | T7, TP7 | | Cathodal | |  |  |  |  |  |  |  |  |  |  |  |  |  |
| Henseler et al. (2014) [142] | Crossover, sham-controlled, participant-blind trial | 36 | 26.2 | tDCS | MNI coordinates −50, 15, 29 (left IFG) | Fp2 | Anodal | | Online assessment- performed during stimulation | | 25 | | 2 mA, 25 cm2 (active), 50 cm2 (reference) | | **PWI paradigm-** name a picture in the presence of a visual distractor word; four experimental distractor conditions: (1) categorically related, (2) associatively related, (3) unrelated control for categorically related distractors, (4) unrelated control for associatively related distractors | | Semantic (picture, word) | | Visual | | Verbal | |
|  |  |  |  |  | MNI coordinates −56, −48, −2 (left pMTG) | Fp2 | Anodal | |  |  |  |  |  |  |  |  |  |  |  |  |  |  |
| Meinzer et al. (2016) [143] | Crossover, sham-controlled, participant-blind trial | 24 | 24.69 | tDCS | Left posterior temporal cortex (pMTG/STG) | Fp2 | Anodal | | Online assessment- performed during stimulation | | 20 | | 1 mA, 35 cm2 (active), 100 cm2 (reference) | | **Blocked cyclic naming paradigm-** common objects; blocks of related and unrelated pictures are used | | Semantic (picture) | | Visual | | Verbal | |
|  |  |  |  |  | Crossing point between T3–Fz and F7–Cz | Fp2 | Anodal | |  |  |  |  |  |  |  |  |  |  |  |  |  |  |
| Pisoni et al. (2012)- Exp. 1 [83] | Crossover, sham-controlled, participant-blind trial | 12 | 22.4 | tDCS | Talairach coordinates: X = −50; Y = −46; Z = 1 (left STG- Wernicke's area) | Fp2 | Anodal | | Offline assessment- performed after stimulation | | 20 | | 2 mA, 35 cm2 | | **Semantic blocked naming paradigm-** blocks consist of objects belonging to the same category (homogeneous sets) or different categories (mixed sets) | | Semantic (picture) | | Visual | | Verbal | |
| Pisoni et al. (2012)- Exp. 2 [83] | Crossover, sham-controlled, participant-blind trial | 12 | 21.8 | tDCS | Crossing point between T3-Fz and F7-Cz (left IFG- Broca's area) | Fp2 | Anodal | | Offline assessment- performed after stimulation | | 20 | | 2 mA, 35 cm2 | | **Semantic blocked naming paradigm-** blocks consist of objects belonging to the same category (homogeneous sets) or different categories (mixed sets) | | Semantic (picture) | | Visual | | Verbal | |
| Westwood et al. (2017)- Exp. 1A [144] | Crossover, sham-controlled, double-blind trial | 18 | 21 | tDCS | F7 | Fp2 | Anodal | | Online assessment- performed during stimulation | | 15 | | 1 mA, 9 cm2 (active), 35 cm2 (reference) | | **Word reading**  **Picture naming** | | Semantic (picture, word) | | Visual | | Verbal | |
| Westwood et al. (2017)- Exp. 1B [144] | Crossover, sham-controlled, double-blind trial | 20 | 21 | tDCS | F7 | Fp2 | Anodal | | Online assessment- performed during stimulation | | 25 | | 1.5 mA, 25 cm2 (active), 35 cm2 (reference) | | **Word reading**  **Picture naming** | | Semantic (picture, word) | | Visual | | Verbal | |
| Westwood et al. (2017)- Exp. 1C [144] | Crossover, sham-controlled, double-blind trial | 18 | 19.8 | tDCS | Halfway between T3 and T5 | Extracephalic | Anodal | | Online assessment- performed during stimulation | | 25 | | 1.5 mA, 25 cm2 (active), 35 cm2 (reference) | | **Word reading**  **Picture naming** | | Semantic (picture, word) | | Visual | | Verbal | |
| Westwood et al. (2017)- Exp. 2 [144] | Crossover, sham-controlled, double-blind trial | 17 | 21 | tDCS | F7 | Fp2 | Anodal | | Online assessment- performed during stimulation | | 25 | | 1.5 mA, 25 cm2 (active), 35 cm2 (reference) | | **Cyclic blocked picture naming-** each set of pictures presented four times in a row | | Semantic (picture) | | Visual | | Verbal | |
| Wirth et al. (2011) [145] | Crossover, sham-controlled, participant-blind trial | 20 | 23.5 | tDCS | Halfway between F3 and AF3 | Extracephalic | Anodal | | Online assessment- performed during stimulation | | 37 | | 1.5 mA, 35 cm2 (active), 49 cm2 (reference) | | **Semantic blocking paradigm-** object pictures of categorically homogeneous and heterogeneous categories are presented in homogeneous and heterogenous blocks; overtly name each object | | Semantic (picture) | | Visual | | Verbal | |
|  |  |  |  |  |  |  |  |  | Offline assessment- performed after stimulation | |  |  |  |  | **Picture naming paradigm** | |  |  |  |  |  |  |
| Younger et al. (2016) [146] | Parallel, sham-controlled, participant-blind trial | Active: 11 Sham: 14 | Active: 26.8 Sham: 26.2 | tDCS | P3 | Fp2 | Anodal | | Offline assessment- performed before and after stimulation (on different days) | | 20 | | 1.5 mA, 25 cm2 | | **SWE subtest of TOWRE** | | Semantic (word) | | Visual | | Verbal | |
| Ihara et al. (2015) [84] | Crossover, sham-controlled, participant-blind trial | 14 | 22-60 | tDCS | Electrode centre over median of Talairach coordinates −45, 28, −16 and −52, 24, 23 (left anterior inferior cortex [LaIFC] and left posterior inferior cortex [LpIFC], respectively) | Cz | Anodal | | Offline assessment- performed after stimulation | | 15 | | 1.5 mA, 35 cm2 | | **SJT-** judge whether target words are semantically related or unrelated to prime words; four conditions: ambiguous words with meanings related to the prime word (RA), ambiguous words with meanings unrelated to the prime word (UA), unambiguous words related to the prime word (RU), unambiguous words unrelated to the prime word | | Semantic (word) | | Visual | | Nonverbal | |
| Cohen-Maximov et al. (2015) [57] | Parallel, sham-controlled, participant-blind trial | RA/LC: 13 LA/RC: 13 Sham: 14 | RA/LC: 24.7 LA/RC: 25.5 Sham: 24 | tDCS | Crossing point between T4-Fz and F8-Cz | Crossing point between T3-Fz and F7-Cz | Bilateral | | Online assessment- performed during stimulation | | 26 | | 1.5 mA, 16 cm2 (anode- active), 35 cm2 (cathode- reference) | | **Gesture task (SJT)-** indicate whether or not a word is related to the gesture performed by an actor in a video clip | | Semantic (picture, word) | | Visual | | Both | |
|  |  |  |  |  | Crossing point between T3-Fz and F7-Cz | Crossing point between T4-Fz and F8-Cz | Bilateral | |  |  |  |  |  |  |  |  |  |  |  |  |  |  |
| Peretz & Lavidor (2013) [147] | Crossover, sham-controlled trial | 17 | 24.4 | tDCS | CP5 | Fp2 | Anodal | | Offline assessment- performed after stimulation | | 10 | | 1 mA, 35 cm2 | | **Explicit semantic relatedness decision task-** decide whether or not prime (ambiguous word) is related to target word; the meaning of prime-target associations can be dominant, subordinate, or unrelated | | Semantic (word) | | Visual | | Verbal | |
|  |  |  |  |  | CP6 | Fp1 | Anodal | |  |  |  |  |  |  |  |  |  |  |  |  |  |  |
| Weltman & Lavidor (2013) [148] | Crossover, sham-controlled, participant-blind trial  But different groups for polarity (LA/RC, RA/LC) | RA/LC: 16 LA/RC: 16 | RA/LC: 25.4 LA/RC: 25.8 | tDCS | CP6 | CP5 | Bilateral (RA/LC) | | Online assessment- performed during stimulation | | 20 | | 1.5 mA, 35 cm2 | | **Lexical decision task**  **Semantic priming task-** presented with a word first and subsequently with a string of characters (word or a pseudo-word); decide whether or not a string of letters is a real word, after first being presented with a real word (prime); prime type: direct, mediated, unrelated | | Semantic (word) | | Visual | | Verbal | |
|  |  |  |  |  | CP5 | CP6 | Bilateral (LA/RC) | |  |  |  |  |  |  |  |  |  |  |  |  |  |  |

| **Reference (Year)** | **Study Design** | **Participant N** | **Mean Age in Years** | **Stimulation Modality** | **Anode** | **Cathode** | **Polarity of Stimulating Electrode** | **Online/Offline** | **Duration** | **Current Intensity, Electrode Size** | **Experimental Task Description** | **Semantic/**  **Less semantic** | **Visual/**  **Auditory** | **Verbal/**  **Nonverbal** |
| --- | --- | --- | --- | --- | --- | --- | --- | --- | --- | --- | --- | --- | --- | --- |
| ***Language*** | | | | | | | | | | | | | | |
| Price et al. (2016) [85] | Crossover, sham-controlled, participant-blind trial | 18 | 25.3 | HD-tDCS | CP5 | C3, T7, P7, P3 | Anodal | Offline assessment- performed after stimulation | 20 | 2 mA, outer diameter: 12 mm, inner diameter: 6 mm (ring electrodes) | **Word-pair task-** decide whether word pair forms a meaningful combination (e.g., tiny radish) or not (e.g., fast blueberry) | Semantic (word) | Visual | Nonverbal |
|  |  |  |  |  | CP6 | C4, T8, P8, P4 | Anodal |  |  |  |  |  |  |  |
| Malyutina et al. (2018) [149] | Crossover, sham-controlled, participant-blind trial  But different groups for polarity (bi, LA, RC) | Bi: 24 Anodal: 24 Cathodal: 24 | 22.9 | tDCS | F7 | F8 | Bilateral | Online/offline- performed during and after stimulation | 20 | 1.5 mA, 25 cm2 | **Single-word-level task (lexical decision)-** decide whether a string of letters is a real word or not  **Sentence-level task (sentence comprehension)-** answer comprehension question with two possible responses after reading a sentence | Semantic (word) | Visual | Nonverbal |
|  |  |  |  |  | F7 | Pz | Anodal |  |  |  |  |  |  |  |
|  |  |  |  |  | Pz | F8 | Cathodal |  |  |  |  |  |  |  |
| Giustolisi et al. (2018) [86] | Parallel, sham-controlled, participant-blind trial | Active: 22 Sham: 22 | 22 | tDCS | F5 | Fp2 | Anodal | Online assessment- performed during stimulation | 30 | 0.75 mA, 9 cm2 (active), 35 cm2 (reference) | **Linguistic task-** judge which of the two picture represents the meaning of the sentence; there are different loads on short-term memory | Semantic (picture, word) | Auditory (sentence), visual (picture) | Verbal and nonverbal |
| Klaus & Hartwigsen (2020) [70] | Crossover, sham-controlled, double-blind trial | 48 | 27.06 | tDCS | Between FC5 and C5 | Centre of forehead | Anodal | Online task- performed during stimulation | 20 | 2 mA, 25 cm2 (active), 100 cm2 (reference) | **Picture naming** | Semantic (picture) | Visual | Nonverbal stimuli verbal response |
| Kurmakaeva et al. (2021) [150] | Parallel, sham-controlled trial | Anodal: 24 Cathodal: 24 Sham: 24 | 17–35 | tDCS | CP5 | Extracephalic (ipsilateral) | Anodal | Offline assessment- performed after stimulation (immediately after training [Day 1], and next day [Day 2]) | 15 | 1.5 mA, 25 cm2 (active), 50 cm2 (reference) | Contextual acquisition of novel concrete and abstract words in short stories, then assessed on lexical and semantic levels:  **Free-form definition task-** provide definitions of 10 new words learned  **SJT-** choose one (out of three) correct definition for each presented word | Semantic (word) | Visual | Verbal stimuli nonverbal response |
|  |  |  |  |  | Extracephalic (ipsilateral) | CP5 | Cathodal |  |  |  |  |  |  |  |
| Fiori et al. (2018) [151] | Crossover, sham-controlled, double-blind trial | 28 | 26.96 | tDCS | FC5 | Fp2 | Anodal | Online assessment- verb learning phase performed during stimulation (T1-T6) | 24 | 1 mA, 35 cm2 | **Verb learning paradigm-** during **picture-word pair learning phase**, read aloud the corresponding verb presented underneath each picture, and for the **verb learning phase**, name the corresponding verb upon the presentation of a picture on its own | Semantic (picture, word) | Visual | Both |
| Marko & Riečanský (2021) [73] | Parallel, sham-controlled, double-blind trial | PFC: 40 TPC: 40 Sham: 41 | 23.1 | tDCS | Between F3 and AF3 | Between T7 and P5 | Bilateral | Online/offline assessment- performed before, during, and after stimulation | 25 | 2 mA, 35 cm2 | **SJT-** decide whether word pair (trial word and current probe) is semantically related or not; types of stimuli: dominant associates, remote associates, unrelated words | Semantic (word) | Visual | Nonverbal |
|  |  |  |  |  | Between T8 and P6 | Between T7 and P5 | Bilateral |  |  |  |  |  |  |  |

| **Reference (Year)** | **Study Design** | **Participant N** | **Mean Age in Years** | **Stimulation Modality** | **Anode** | **Cathode** | **Polarity of Stimulating Electrode** | **Online/Offline** | **Duration** | **Current Intensity, Electrode Size** | **Experimental Task Description** | **Semantic/**  **Less semantic** | **Visual/**  **Auditory** | **Verbal/**  **Nonverbal** |
| --- | --- | --- | --- | --- | --- | --- | --- | --- | --- | --- | --- | --- | --- | --- |
| ***Language*** | | | | | | | | | | | | | | |
| Longo et al. (2022) [152] | Crossover, sham-controlled, double-blind trial  But different groups for polarity (AL, AR, CL, CR) | LA: 18 RA: 18 LC: 18 RC: 18 | 25.49 | tDCS | P3 | Fp2 | Anodal | Online assessment- performed during stimulation | 20 | 1.5 mA, 35 cm2 (active), 100 cm2 (reference) | **Semantic categorisation task-** decide whether a word represents a living or non-living concept; further divided into high number of features (NOF+; semantically rich concepts) and low number of features (NOF-; semantically non-rich concepts) | Semantic (word) | Visual | Nonverbal |
|  |  |  |  |  | P4 | Fp1 | Anodal |  |  |  |  |  |  |  |
|  |  |  |  |  | Fp2 | P3 | Cathodal |  |  |  |  |  |  |  |
|  |  |  |  |  | Fp1 | P4 | Cathodal |  |  |  |  |  |  |  |

| **Reference (Year)** | **Study Design** | **Participant N** | **Mean Age in Years** | **Stimulation Modality** | **Anode** | | **Cathode** | | | | **Polarity of Stimulating Electrode** | **Online/Offline** | **Duration** | **Current Intensity, Electrode Size** | **Experimental Task Description** | **Semantic/**  **Less semantic** | **Visual/**  **Auditory** | **Verbal/**  **Nonverbal** |
| --- | --- | --- | --- | --- | --- | --- | --- | --- | --- | --- | --- | --- | --- | --- | --- | --- | --- | --- |
| ***Creativity*** |  |  |  |  |  | |  | | | |  |  |  |  |  |  |  |  |
| Bartel et al. (2020) [153] | Parallel, sham-controlled, double-blind trial | Active: 20  Sham: 20 | 25.0 | tDCS | Between F3 and F5 | | Fp2 | | | | Anodal | Online assessment- performed before and during stimulation | 20 | 1.5mA, 35cm2 | **Rorschach test-** think of and name as many things as participants could see in the ambiguous inkblot pictures | Less semantic | Visual | Nonverbal |
| Brunye et al. (2015) [154] | Crossover, sham-controlled, double-blind trial | 16 | 20.5 | HD-tDCS | AFZ | | Fp1, AF3, F1, Fz | | | | Anodal | Online assessment- performed during stimulation | n.r. | 2mA, electrode size n.r. | **Cued free association test-** type the first word that come into mind when participants see the target word; informed that there was no right or wrong answer | Semantic (word) | Visual | Verbal |
|  |  |  |  |  | T7 | | TP7, FT7, FC5, CP5 | | | |  |  |  |  |  |  |  |  |
| Cerruti & Schlaug (2009)- Exp. 1 [2] | Crossover, sham-controlled, double-blind trial | 18 | 25.5 | tDCS | F3 | | Fp2 | | | | Anodal | Offline assessment – performed after stimulation | 20 | 1 mA, 16.3 cm2 (active), 30 cm2 (reference) | **RAT** | Semantic (word) | Visual | Verbal |
| Chrysikou et al. (2013) [60] | Parallel, sham-controlled, double-blind trial | L/CU: 8 R/CU: 8 S/CU: 8 L/UU: 8 R/UU: 8 S/UU: 8 | 23.38 | tDCS | Extracephalic | | F7 | | | | Cathodal | Online assessment- performed during stimulation | Max. 20 | 1.5 mA, 25 cm2 | **AUT-** generate unconventional (uncommon) uses of daily objects | Semantic (picture) | Visual | Verbal |
|  |  |  |  |  | Extracephalic | | F8 | | | | Cathodal |  |  |  |  |  |  |  |
| Chrysikou et al. (2021)- Exp. 1 [155] | Parallel, sham-controlled, participant-blind trial | LC: 20  RC: 20  Sham: 20 | 19.20 | tDCS | F7 | | Extracephalic | | | | Cathodal | Online assessment- performed during stimulation | 20 | 1.5mA, 25cm2 | **AUT (common, uncommon uses)** | Semantic (word) | Visual | Verbal |
|  |  |  |  |  | F8 | |  |  |  |  |  |  |  |  |  |  |  |  |
| Chrysikou et al. (2021)- Exp. 2 [155] | Parallel, sham-controlled, participant-blind trial | LA: 20  RA: 20  Sham: 20 | 18.83 | tDCS | F7 | | Extracephalic | | | | Anodal | Online assessment- performed during stimulation | 20 | 1.5mA, 25cm2 | **AUT (common, uncommon uses)** | Semantic (word) | Visual | Verbal |
|  |  |  |  |  | F8 | |  |  |  |  |  |  |  |  |  |  |  |  |
| Chrysikou et al. (2021)- Exp. 3 [155] | Parallel, sham-controlled, participant-blind trial | LA/RC: 20  LC/RA: 20  Sham: 20 | 19.02 | tDCS | F7 | | F8 | | | | Bilateral | Online assessment- performed during stimulation | 20 | 1.5mA, 25cm2 | **AUT (common, uncommon uses)** | Semantic (word) | Visual | Verbal |
|  |  |  |  |  | F8 | | F7 | | | |  |  |  |  |  |  |  |  |
|  |  |  |  |  | F3 | | Extracephalic | | | |  |  |  |  |  |  |  |  |
| Green et al. (2017) [157] | Parallel, sham-controlled, double-blind trial | Active: 15  Sham: 16 | 21.69 | HD-tDCS | AF3 | | Fpz, Fz, F7, FC3 | | | | Anodal | Online assessment- performed during stimulation | 20 | 2mA, electrode size n.r. | **Analogy finding task-** judge if a word pair can form valid analogy or not | Semantic (word) | Visual | Verbal |
| Hertenstein et al. (2019) [158] | Parallel, sham-controlled, double-blind trial | LA/RC: 30  LC/RA: 30  Sham: 30 | 23.8 | tDCS | Crossing point between T4-Fz and F8-Cz | | Crossing point between T3-Fz and F7-Cz | | | | Bilateral | Online assessment- performed before and during stimulation | 20 | 1mA, 35cm2 | **AUT**  **RAT**  **WCST** | Semantic (word) for AUT & RAT, less semantic for WCST | Visual | Verbal for RAT, AUT, nonverbal for WCST |
|  |  |  |  |  | Crossing point between T3-Fz and F7-Cz | | Crossing point between T4-Fz and F8-Cz | | | |  |  |  |  |  |  |  |  |
| Koizumi et al. (2020) [159] | Crossover, sham-controlled trial | 14 | 23.0 | tDCS | F3 | | P4 | | | | Bilateral | Offline assessment- performed before and after stimulation | 20 | 2mA, 35cm2 | **AUT** | Semantic (word) | Visual | Verbal |
| Li et al. (2022) [160] | Crossover, sham-controlled trial | 26 | 19.31 | tDCS | F3 | | F4 | | | | Bilateral | Online assessment- performed during stimulation | 21 | 1.5mA, 25cm2 | **AUT**  **RAT** | Semantic (word) | Visual | Verbal |
|  |  |  |  |  | F4 | | F3 | | | |  |  |  |  |  |  |  |  |
| Metuki et al. (2012) [15] | Crossover, sham-controlled trial | 21 | 23.1 | tDCS | F3 | | Fp2 | | | | Anodal | Online assessment- performed during stimulation | Max. 11 | 1 mA, 35 cm2 | **CRAT-** easy and difficult items | Semantic (word) | Visual | Verbal |
| ***Creativity*** | | | | | | | | | | | | | | | | | | |
| Mayseless & Shamay-Tsoory (2015)- Exp. 1 [161] | Crossover, sham-controlled trial  But different groups for polarity (LA/RC, LC/RA) | LA/RC: 15  LC/RA: 15 | 24.44 | tDCS | Crossing point between T4-Fz and F8-Cz | Crossing point between T3-Fz and F7-Cz | | | | Bilateral | | Online assessment- performed during stimulation | 21 | 1.5mA, 25cm2 | **AUT**  **VFT (phonemic)** | Semantic (word) | Visual | Verbal |
|  |  |  |  |  | Crossing point between T3-Fz and F7-Cz | Crossing point between T4-Fz and F8-Cz | | | |  |  |  |  |  |  |  |  |  |
| Mayseless & Shamay-Tsoory (2015)- Exp. 2 [161] | Crossover, sham-controlled trial  But different groups for polarity (anodal, cathodal) | Anodal: 15  Cathodal: 15 | 24.17 | tDCS | Crossing point between T4-Fz and F8-Cz | Fp1 | | | Anodal | | | Online assessment- performed during stimulation | 21 | 1.5mA, 25cm2 | **AUT**  **VFT (phonemic)** | Semantic (word) | Visual | Verbal |
|  |  |  |  |  | Crossing point between T3-Fz and F7-Cz | Fp2 | | | Cathodal | | |  |  |  |  |  |  |  |
| Pick & Lavidor (2019)- Exp. 1 [162] | Crossover, sham-controlled trial  But different groups for polarity (bilateral, cathodal) | Bi: 16  Cathodal: 16 | 22.81 | tDCS | P3, P4 | Extracephalic | | Bilateral (anodal) | | | | Online assessment- performed during stimulation | 17 | 1mA, 9cm2 (active), 35cm2 (reference) | **RAT** | Semantic (word) | Visual | Verbal |
|  |  |  |  |  | P4 | Extracephalic | | Cathodal | | | |  |  |  |  |  |  |  |
| Ruggiero et al. (2018) [163] | Parallel, sham-controlled trial | LA/RC: 7  LC/RA: 7  Sham: 7 | 26.58 | tDCS | Halfway between T7 and FT7 | Halfway between T8 and FT8 | | | | Bilateral | | Online assessment: DTT performed during stimulation  Offline assessment: RAT performed before and after stimulation | **20** | 1.5mA, 35cm2 | **RAT**  **DTT-** given a line in a frame, asked to produce drawing within the frame; a title has to be assigned to each frame | Semantic (word) for RAT, less semantic for DTT | Visual | Verbal for RAT, nonverbal for DTT |
|  |  |  |  |  | Halfway between T8 and FT8 | Halfway between T7 and FT7 | | | |  |  |  |  |  |  |  |  |  |
| Salvi et al. (2020) [164] | Parallel, sham-controlled, double-blind trial | T8: 36  AF3: 37  Sham: 36 | n.r. | HD-tDCS | T8 | FC6, FT10, TP10, CP6 | | | | Anodal | | Online assessment- performed before and during stimulation | 20 | 2mA, electrode size n.r. | **RAT** | Semantic (word) | Visual | Verbal |
|  |  |  |  |  | AF3 | FPz, Fz, F7, FC3 | | | |  |  |  |  |  |  |  |  |  |
| Xiang et al. (2021) [165] | Crossover, sham-controlled trial | 20 | 19.60 | tDCS | F4 | F3 | | | | Bilateral | | Online assessment- performed during stimulation | 20 | 1.5mA, 25cm2 | **AUT**  **RAT** | Semantic (word) | Visual | Nonverbal |
|  |  |  |  |  | F3 | F4 | | | |  |  |  |  |  |  |  |  |  |
| Aihara et al. (2017) [166] | Crossover, sham-controlled trial | 31 | 29.3 | tDCS | 1.5cm anterior to T4 | 1.5cm anterior to T3 | | | | Bilateral | | Online assessment – performed during stimulation | 25 | 1.6mA, 35cm2 | **Matchstick arithmetic task**  **Verbal insight problem solving** | Less semantic  Semantic (word) | Visual | Nonverbal  Verbal |
|  |  |  |  |  | 1.5cm anterior to T3 | 1.5cm anterior to T4 | | | |  |  |  |  |  |  |  |  |  |
| Luft et al. (2017) [167] | Parallel, sham-controlled trial | 60 | 23.0 | HD-tDCS | F3 | T7, Cz, Fp2 | | | | Anodal | | Offline assessment – performed before and after stimulation | 15 | 1mA | **Matchstick arithmetic task** | Less semantic | Visual | Nonverbal |
|  |  |  |  |  | T7, Cz, Fp2 | F3 | | | | Cathodal | |  |  |  |  |  |  |  |
| Chi and Snyder (2012) [168] | Parallel, sham-controlled trial | 22 | 30.6 | tDCS | T8 | T7 | | | | Bilateral | | Offline assessment – performed before and after stimulation | 10 | 2mA, 35cm2 | **Nine dot problem** | Less semantic | Visual | Nonverbal |
| Guo et al. (2023) [169] | Parallel, sham-controlled trial | 36 | 21.6 | tDCS | F4 | Fp1 | | | | Anodal | | Offline assessment –after stimulation | 20 | 1.5mA, 25cm2 | **Product design task** | Semantic (picture) | Visual | Verbal (annotation) |
| Lundie et al. (2022) [170] | Parallel, sham-controlled trial | 48 | 21.7 | HD-tDCS | AF3 | FpZ, FZ, FC1 | | | | Anodal | | Online assessment – performed during stimulation | 21 | 2mA | **Radiation problem solving (analogical reasoning)** | Semantic | Visual | Verbal |
| Wang et al. (2022) [171] | Parallel, sham-controlled trial | 70 | 19.6 | tDCS | F4 | F3 | | | | Bilateral | | Online assessment – performed during stimulation | 20 | 1.5mA, 25cm2 | **RAT**  **AUT** | Semantic | Visual | Verbal |
| Wang et al. (2023) [172] | Parallel, sham-controlled trial | 60 | 19.5 | tDCS | F3 | F4 | | | | Bilateral | | Offline assessment – performed before and after stimulation | 20 | 1.5mA, 25cm2 | **AUT** | Semantic | Visual | Verbal |

Abbreviations: n.r. = not reported; mA = milliamperes, HD = high-definition, LA = left hemisphere – anodal, LC = left hemisphere – cathodal, RA = right hemisphere – anodal, RC = right hemisphere – cathodal, LS = left sham, RS = right sham, Bi = bilateral, L = left, R = right, GNG = go/no-go, VFT = verbal fluency test, RAT = Remote Associates Test, BART = Balloon Analogue Risk Task, SCWT = Stroop Colour and Word Test, AUT = alternate uses task, TOL = Tower of London, IAT = Implicit Association Test, CRAT = Compound Remote Associates Test, PGNG = parametric go/no-go, ICT = Intertemporal Choice Task, PRLT = probabilistic reversal learning task, TMT B = Trail Making Test B, CRT = choice reaction time, SST = stop signal task, GNG-SST = go/no-go-stop signal task, TOH = Tower of Hanoi, IGT = Iowa Gambling Task, WCST = Wisconsin Card Sorting Test, SAT = shifting attention test, ANT = Attention Network Test, RSE = redundant signal effect, CalCAP = California Computerized Assessment Package, SRT = simple reaction time, DT = detection task, IT = identification task, OBT = one back task, CMT = continuous monitoring task, AB = attentional blink, TMT A = Trail Making Test A, VDT = visual discrimination task, CPT = continuous performance task, WM = working memory, OCLT = one card learning task, DSF = digit span forward, DSB = digit span backward, DS = digit span, CDT = change detection task, CBT = Corsi block-tapping test, VOMT = visual-object matching task, STL = short-term learning, LTR = long-term retrieval, LTM = long-term memory, RPP = retrieval practice paradigm, TAVEC = Complutense Verbal Learning Test, AGL = artificial grammar learning, K-BNT = Korean-Boston Naming Test, PWI = picture-word interference, SWE = Sight word efficiency, TOWRE = Test of Word Reading Efficiency, SJT = semantic judgement task, recon. = reconsolidation, DTT = divergent thinking test, FPT = five-point test

**References**

[1] Boggio PS, Rocha RR, da Silva MT, Fregni F. Differential modulatory effects of transcranial direct current stimulation on a facial expression go-no-go task in males and females. Neurosci Lett 2008;447(2-3):101-5. https://doi.org/10.1016/j.neulet.2008.10.009.

[2] Cerruti C, Schlaug G. Anodal transcranial direct current stimulation of the prefrontal cortex enhances complex verbal associative thought. J Cogn Neurosci 2009;21(10):1980-7. https://doi.org/10.1162/jocn.2008.21143.

[3] Fecteau S, Knoch D, Fregni F, Sultani N, Boggio P, Pascual-Leone A. Diminishing risk-taking behavior by modulating activity in the prefrontal cortex: A direct current stimulation study. J Neurosci 2007a;27(46):12500-5. https://doi.org/10.1523/JNEUROSCI.3283-07.2007.

[4] Iyer MB, Mattu U, Grafman J, Lomarev M, Sato S, Wassermann EM. Safety and cognitive effect of frontal DC brain polarization in healthy individuals. Neurology 2005;64(5):872-5. https://doi.org/10.1212/01.WNL.0000152986.07469.E9.

[5] Mameli F, Mrakic-Sposta S, Vergari M, Fumagalli M, Macis M, Ferrucci R, et al. Dorsolateral prefrontal cortex specifically processes general – but not personal – knowledge deception: Multiple brain networks for lying. Behav Brain Res 2010;211(2):164-8. https://doi.org/10.1016/j.bbr.2010.03.024.

[6] Stone DB, Tesche CD. Transcranial direct current stimulation modulates shifts in global/local attention. Neuroreport 2009;20(12):1115-9. https://doi.org/10.1097/WNR.0b013e32832e9aa2.

[7] Fecteau S, Pascual-Leone A, Zald DH, Liguori P, Theoret H, Boggio PS, et al. Activation of prefrontal cortex by transcranial direct current stimulation reduces appetite for risk during ambiguous decision making. J Neurosci 2007b;27(23):6212-8. https://doi.org/10.1523/JNEUROSCI.0314-07.2007.

[8] Beeli G, Casutt G, Baumgartner T, Jäncke L. Modulating presence and impulsiveness by external stimulation of the brain. Behav Brain Funct 2008;4(1):33-. https://doi.org/10.1186/1744-9081-4-33.

[9] Dockery CA, Hueckel-Weng R, Birbaumer N, Plewnia C. Enhancement of planning ability by transcranial direct current stimulation. J Neurosci 2009;29(22):7271-7. https://doi.org/10.1523/JNEUROSCI.0065-09.2009.

[10] Leite J, Carvalho S, Fregni F, Gonçalves ÓF. Task-specific effects of tDCS-induced cortical excitability changes on cognitive and motor sequence set shifting performance. PLOS One 2011;6(9):e24140-e. https://doi.org/10.1371/journal.pone.0024140.

[11] Balconi M, Vitaloni S. The tDCS effect on alpha brain oscillation for correct vs. incorrect object use. The contribution of the left DLPFC. Neurosci Lett 2012;517(1):25-9. https://doi.org/10.1016/j.neulet.2012.04.010.

[12] Gladwin TE, den Uyl TE, Wiers RW. Anodal tDCS of dorsolateral prefontal cortex during an Implicit Association Test. Neurosci Lett 2012;517(2):82-6. https://doi.org/10.1016/j.neulet.2012.04.025.

[13] Jeon SY, Han SJ. Improvement of the working memory and naming by transcranial direct current stimulation. Ann Rehabil Med 2012;36(5):585-95. https://doi.org/10.5535/arm.2012.36.5.585.

[14] Leite J, Carvalho S, Fregni F, Boggio PS, Gonçalves ÓF. The effects of cross-hemispheric dorsolateral prefrontal cortex transcranial direct current stimulation (tDCS) on task switching. Brain Stimul 2013;6(4):660-7. https://doi.org/10.1016/j.brs.2012.10.006.

[15] Metuki N, Sela T, Lavidor M. Enhancing cognitive control components of insight problems solving by anodal tDCS of the left dorsolateral prefrontal cortex. Brain Stimul 2012;5(2):110-5. https://doi.org/10.1016/j.brs.2012.03.002.

[16] Minati L, Campanhã C, Critchley HD, Boggio PS. Effects of transcranial direct-current stimulation (tDCS) of the dorsolateral prefrontal cortex (DLPFC) during a mixed-gambling risky decision-making task. Cogn Neurosc 2012;3(2):80-8. https://doi.org/10.1080/17588928.2011.628382.

[17] Vannorsdall TD, Schretlen DJ, Andrejczuk M, Ledoux K, Bosley LV, Weaver JR, et al. Altering automatic verbal processes with transcranial direct current stimulation. Front Psychiatry 2012;3:73-. https://doi.org/10.3389/fpsyt.2012.00073.

[18] Fecteau S, Boggio P, Fregni F, Pascual-Leone A. Modulation of untruthful responses with non-invasive brain stimulation. Front Psychiatry 2013;3:97-. https://doi.org/10.3389/fpsyt.2012.00097.

[19] Iuculano T, Kadosh RC. The mental cost of cognitive enhancement. J Neurosci 2013;33(10):4482-6. https://doi.org/10.1523/JNEUROSCI.4927-12.2013.

[20] Plewnia C, Zwissler B, Längst I, Maurer B, Giel K, Krüger R. Effects of transcranial direct current stimulation (tDCS) on executive functions: Influence of COMT Val/Met polymorphism. Cortex 2013;49(7):1801-7. https://doi.org/10.1016/j.cortex.2012.11.002.

[21] Balconi M, Vitaloni S. Dorsolateral pFC and the representation of the incorrect use of an object: The transcranial direct current stimulation effect on N400 for visual and linguistic stimuli. J Cogn Neurosci 2014;26(2):305-18. https://doi.org/10.1162/jocn_a_00500.

[22] Weber MJ, Messing SB, Rao H, Detre JA, Thompson-Schill SL. Prefrontal transcranial direct current stimulation alters activation and connectivity in cortical and subcortical reward systems: A tDCS-fMRI study. Hum Brain Mapp 2014;35(8):3673-86. https://doi.org/10.1002/hbm.22429.

[23] Baumert A, Buchholz N, Zinkernagel A, Clarke P, MacLeod C, Osinsky R, et al. Causal underpinnings of working memory and Stroop interference control: Testing the effects of anodal and cathodal tDCS over the left DLPFC. Cogn Affect Behav Neurosci 2020;20(1):34-48. https://doi.org/10.3758/s13415-019-00726-y.

[24] Nieratschker V, Kiefer C, Giel K, Krüger R, Plewnia C. The COMT Val/Met polymorphism modulates effects of tDCS on response inhibition. Brain Stimul 2015;8(2):283-8. https://doi.org/10.1016/j.brs.2014.11.009.

[25] Shen B, Yin Y, Wang J, Zhou X, McClure SM, Li J. High-definition tDCS alters impulsivity in a baseline-dependent manner. NeuroImage 2016;143:343-52. https://doi.org/10.1016/j.neuroimage.2016.09.006.

[26] Albein-Urios N, Chase H, Clark L, Kirkovski M, Davies C, Enticott PG. Increased perseverative errors following high-definition transcranial direct current stimulation over the ventrolateral cortex during probabilistic reversal learning. Brain Stimul 2019;12(4):959-66. https://doi.org/10.1016/j.brs.2019.02.013.

[27] To WT, Eroh J, Hart J, Vanneste S. Exploring the effects of anodal and cathodal high definition transcranial direct current stimulation targeting the dorsal anterior cingulate cortex. Sci Rep 2018;8(1):4454-16. https://doi.org/10.1038/s41598-018-22730-x.

[28] Guo H, Zhang Z, Da S, Sheng X, Zhang X. High‐definition transcranial direct current stimulation (HD‐tDCS) of left dorsolateral prefrontal cortex affects performance in Balloon Analogue Risk Task (BART). Brain Behav 2018;8(2):e00884-n/a. https://doi.org/10.1002/brb3.884.

[29] Bender AD, Filmer HL, Dux PE. Transcranial direct current stimulation of superior medial frontal cortex disrupts response selection during proactive response inhibition. NeuroImage 2017;158:455-65. https://doi.org/10.1016/j.neuroimage.2016.10.035.

[30] Campanella S, Schroder E, Vanderhasselt M-A, Baeken C, Kornreich C, Verbanck P, et al. Short-term impact of tDCS over the right inferior frontal cortex on impulsive responses in a go/no-go task. Clin EEG Neurosci 2018;49(6):398-406. https://doi.org/10.1177/1550059418777404.

[31] Castro-Meneses LJ, Johnson BW, Sowman PF. Vocal response inhibition is enhanced by anodal tDCS over the right prefrontal cortex. Exp Brain Res 2016;234(1):185-95. https://doi.org/10.1007/s00221-015-4452-0.

[32] Cunillera T, Fuentemilla L, Brignani D, Cucurell D, Miniussi C. A simultaneous modulation of reactive and proactive inhibition processes by anodal tDCS on the right inferior frontal cortex. PLOS One 2014;9(11):e113537-e. https://doi.org/10.1371/journal.pone.0113537.

[33] Cunillera T, Brignani D, Cucurell D, Fuentemilla L, Miniussi C. The right inferior frontal cortex in response inhibition: A tDCS–ERP co-registration study. NeuroImage 2016;140:66-75. https://doi.org/10.1016/j.neuroimage.2015.11.044.

[34] Dambacher F, Schuhmann T, Lobbestael J, Arntz A, Brugman S, Sack AT. No effects of bilateral tDCS over inferior frontal gyrus on response inhibition and aggression. PLOS One 2015;10(7):e0132170-e. https://doi.org/10.1371/journal.pone.0132170.

[35] Friehs MA, Frings C. Pimping inhibition: Anodal tDCS enhances stop-signal reaction time. J Exp Psychol Hum Percept Perform 2018;44(12):1933-45. https://doi.org/10.1037/xhp0000579.

[36] Friehs MA, Frings C. Cathodal tDCS increases stop-signal reaction time. Cogn Affect Behav Neurosci 2019b;19(5):1129-42. https://doi.org/10.3758/s13415-019-00740-0.

[37] Jacobson L, Javitt DC, Lavidor M. Activation of inhibition: Diminishing impulsive behavior by direct current stimulation over the inferior frontal gyrus. J Cogn Neurosci 2011;23(11):3380-7. https://doi.org/10.1162/jocn_a_00020.

[38] Kwon YH, Kwon JW. Response inhibition induced in the stop-signal task by transcranial direct current stimulation of the pre-supplementary motor area and primary sensoriomotor cortex. J Phys Ther Sci 2013a;25(9):1083-6. https://doi.org/10.1589/jpts.25.1083.

[39] Lapenta OM, Fregni F, Oberman LM, Boggio PS. Bilateral temporal cortex transcranial direct current stimulation worsens male performance in a multisensory integration task. Neurosci Lett 2012;527(2):105-9. https://doi.org/10.1016/j.neulet.2012.08.076.

[40] Leite J, Gonçalves ÓF, Pereira P, Khadka N, Bikson M, Fregni F, et al. The differential effects of unihemispheric and bihemispheric tDCS over the inferior frontal gyrus on proactive control. Neurosci Res 2018;130:39-46. https://doi.org/10.1016/j.neures.2017.08.005.

[41] Nejati V, Salehinejad MA, Nitsche MA. Interaction of the left dorsolateral prefrontal cortex (l-DLPFC) and right orbitofrontal cortex (OFC) in hot and cold executive functions: Evidence from transcranial direct current stimulation (tDCS). Neurosci 2018;369:109-23. https://doi.org/10.1016/j.neuroscience.2017.10.042.

[42] Ouellet J, McGirr A, Van den Eynde F, Jollant F, Lepage M, Berlim MT. Enhancing decision-making and cognitive impulse control with transcranial direct current stimulation (tDCS) applied over the orbitofrontal cortex (OFC): A randomized and sham-controlled exploratory study. J Psychiatr Res 2015;69:27-34. https://doi.org/10.1016/j.jpsychires.2015.07.018.

[43] Sallard E, Mouthon M, De Pretto M, Spierer L. Modulation of inhibitory control by prefrontal anodal tDCS: A crossover double-blind sham-controlled fMRI study. PLOS One 2018;13(3):e0194936-e. https://doi.org/10.1371/journal.pone.0194936.

[44] Sedgmond J, Lawrence NS, Verbruggen F, Morrison S, Chambers CD, Adams RC. Prefrontal brain stimulation during food-related inhibition training: Effects on food craving, food consumption and inhibitory control. R Soc Open Sci 2019;6(1):181186-. https://doi.org/10.1098/rsos.181186.

[45] Stramaccia DF, Penolazzi B, Sartori G, Braga M, Mondini S, Galfano G. Assessing the effects of tDCS over a delayed response inhibition task by targeting the right inferior frontal gyrus and right dorsolateral prefrontal cortex. Exp Brain Res 2015;233(8):2283-90. https://doi.org/10.1007/s00221-015-4297-6.

[46] Stramaccia DF, Penolazzi B, Altoè G, Galfano G. TDCS over the right inferior frontal gyrus disrupts control of interference in memory: A retrieval-induced forgetting study. Neurobiol Learn Mem 2017;144:114-30. https://doi.org/10.1016/j.nlm.2017.07.005.

[47] Yu J, Tseng P, Hung D, L. , Wu SW, Juan CH. Brain stimulation improves cognitive control by modulating medial-frontal activity and preSMA-vmPFC functional connectivity: Brain stimulation improves cognitive control. Hum Brain Mapp 2015;36:4004-15. https://doi.org/10.1002/hbm.22893.

[48] Sandrini M, Xu B, Volochayev R, Awosika O, Wang W-T, Butman JA, et al. Transcranial direct current stimulation facilitates response inhibition through dynamic modulation of the fronto-basal ganglia network. Brain Stimul 2020;13(1):96-104. https://doi.org/10.1016/j.brs.2019.08.004.

[49] Strobach T, Antonenko D, Schindler T, Flöel A, Schubert T. Modulation of executive control in the task switching paradigm with transcranial direct current stimulation (tDCS). J Psychophysiol 2016;30(2):55-65. https://doi.org/10.1027/0269-8803/a000155.

[50] Tayeb Y, Lavidor M. Enhancing switching abilities: Improving practice effect by stimulating the dorsolateral pre frontal cortex. Neurosci 2016;313:92-8. https://doi.org/10.1016/j.neuroscience.2015.11.050.

[51] Kwon YH, Kwon JW. Is transcranial direct current stimulation a potential method for improving response inhibition? Neural Regen Res 2013b;8(11):1048-54. https://doi.org/10.3969/j.issn.1673-5374.2013.11.011.

[52] Loftus AM, Yalcin O, Baughman FD, Vanman EJ, Hagger MS. The impact of transcranial direct current stimulation on inhibitory control in young adults. Brain Behav 2015;5(5):e00332-n/a. https://doi.org/10.1002/brb3.332.

[53] Zmigrod S, Zmigrod L, Hommel B. Transcranial direct current stimulation (tDCS) over the right dorsolateral prefrontal cortex affects stimulus conflict but not response conflict. Neurosci 2016;322:320-5. https://doi.org/10.1016/j.neuroscience.2016.02.046.

[54] Cattaneo Z, Pisoni A, Papagno C. Transcranial direct current stimulation over Broca's region improves phonemic and semantic fluency in healthy individuals. Neurosci 2011;183:64-70. https://doi.org/10.1016/j.neuroscience.2011.03.058.

[55] Meinzer M, Antonenko D, Lindenberg R, Hetzer S, Ulm L, Avirame K, et al. Electrical brain stimulation improves cognitive performance by modulating functional connectivity and task-specific activation. J Neurosci 2012;32(5):1859-66. https://doi.org/10.1523/JNEUROSCI.4812-11.2012.

[56] Penolazzi B, Pastore M, Mondini S. Electrode montage dependent effects of transcranial direct current stimulation on semantic fluency. Behav Brain Res 2013;248:129-35. https://doi.org/10.1016/j.bbr.2013.04.007.

[57] Cohen-Maximov T, Avirame K, Flöel A, Lavidor M. Modulation of gestural-verbal semantic integration by tDCS. Brain Stimul 2015;8(3):493-8. https://doi.org/10.1016/j.brs.2014.12.001.

[58] Ehlis A-C, Haeussinger FB, Gastel A, Fallgatter AJ, Plewnia C. Task-dependent and polarity-specific effects of prefrontal transcranial direct current stimulation on cortical activation during word fluency. NeuroImage 2016;140:134-40. https://doi.org/10.1016/j.neuroimage.2015.12.047.

[59] Vannorsdall TD, Van Steenburgh JJ, Schretlen DJ, Jayatillake R, Skolasky RL, Gordon B. Reproducibility of tDCS results in a randomized trial: Failure to replicate findings of tDCS-induced enhancement of verbal fluency. Cogn Behav Neurol 2016;29(1):11-7. https://doi.org/10.1097/WNN.0000000000000086.

[60] Chrysikou EG, Hamilton RH, Coslett HB, Datta A, Bikson M, Thompson-Schill SL. Noninvasive transcranial direct current stimulation over the left prefrontal cortex facilitates cognitive flexibility in tool use. Cogn Neurosc 2013;4(2):81-9. https://doi.org/10.1080/17588928.2013.768221.

[61] Balconi M, Canavesio Y, Vitaloni S. Activation of the prefrontal cortex and posterior parietal cortex increases the recognition of semantic violations in action representation. Brain Stimul 2014;7(3):435-42. https://doi.org/10.1016/j.brs.2014.01.011.

[62] Gbadeyan O, McMahon K, Steinhauser M, Meinzer M. Stimulation of dorsolateral prefrontal cortex enhances adaptive cognitive control: A high-definition transcranial direct current stimulation study. J Neurosci 2016;36(50):12530-6. https://doi.org/10.1523/JNEUROSCI.2450-16.2016.

[63] Mansouri FA, Fehring DJ, Feizpour A, Gaillard A, Rosa MGP, Rajan R, et al. Direct current stimulation of prefrontal cortex modulates error-induced behavioral adjustments. Eur J Neurosci 2016;44(2):1856-69. https://doi.org/10.1111/ejn.13281.

[64] Sdoia S, Zivi P, Ferlazzo F. Anodal tDCS over the right parietal but not frontal cortex enhances the ability to overcome task set inhibition during task switching. PLOS One 2020;15(2):e0228541-e. https://doi.org/10.1371/journal.pone.0228541.

[65] Lu H, Gong Y, Huang P, Zhang Y, Guo Z, Zhu X, et al. Effect of repeated anodal HD-tDCS on executive functions: Evidence from a pilot and single-blinded fNIRS study. Front Hum Neurosci 2021;14:583730-. https://doi.org/10.3389/fnhum.2020.583730.

[66] Hasan MA, Shahid H, Ahmed Qazi S, Ejaz O, Danish Mujib M, Vuckovic A. Underpinning the neurological source of executive function following cross hemispheric tDCS stimulation. Int J Psychophysiol 2023;185:1-10. https://doi.org/10.1016/j.ijpsycho.2023.01.004.

[67] Fujiyama H, Tan J, Puri R, Hinder MR. Influence of tDCS over right inferior frontal gyrus and pre-supplementary motor area on perceptual decision-making and response inhibition: A healthy ageing perspective. Neurobiol Aging 2022;109:11-21. https://doi.org/10.1016/j.neurobiolaging.2021.09.014.

[68] Perrotta D, Bianco V, Berchicci M, Quinzi F, Perri RL. Anodal tDCS over the dorsolateral prefrontal cortex reduces Stroop errors. A comparison of different tasks and designs. Behav Brain Res 2021;405:113215-. https://doi.org/10.1016/j.bbr.2021.113215.

[69] Mattavelli G, Lo Presti S, Tornaghi D, Canessa N. High-definition transcranial direct current stimulation of the dorsal anterior cingulate cortex modulates decision-making and executive control. Brain Struct Funct 2022;227(5):1565-76. https://doi.org/10.1007/s00429-022-02456-3.

[70] Klaus J, Hartwigsen G. Failure to improve verbal fluency with transcranial direct current stimulation. Neurosci 2020;449:123-33. https://doi.org/10.1016/j.neuroscience.2020.09.003.

[71] Westwood SJ, Romani C. Null effects on working memory and verbal fluency tasks when applying anodal tDCS to the inferior frontal gyrus of healthy participants. Front Neurosc 2018;12:166-. https://doi.org/10.3389/fnins.2018.00166.

[72] Lo OY, Donkelaar P, Chou LS. Effects of transcranial direct current stimulation over right posterior parietal cortex on attention function in healthy young adults. Eur J Neurosci 2019;49(12):1623-31. https://doi.org/10.1111/ejn.14349.

[73] Marko M, Riečanský I. The left prefrontal cortex supports inhibitory processing during semantic memory retrieval. Cortex 2021;134:296-306. https://doi.org/10.1016/j.cortex.2020.11.001.

[74] Lu H, Liu Q, Guo Z, Zhou G, Zhang Y, Zhu X, et al. Modulation of repeated anodal HD-tDCS on attention in healthy young adults. Front Psychol 2020;11:564447-. https://doi.org/10.3389/fpsyg.2020.564447.

[75] Dubreuil-Vall L, Chau P, Ruffini G, Widge AS, Camprodon JA. tDCS to the left DLPFC modulates cognitive and physiological correlates of executive function in a state-dependent manner. Brain Stimul 2019;12(6):1456-63. https://doi.org/10.1016/j.brs.2019.06.006.

[76] Bolognini N, Fregni F, Casati C, Olgiati E, Vallar G. Brain polarization of parietal cortex augments training-induced improvement of visual exploratory and attentional skills. Brain Res 2010a;1349:76-89. https://doi.org/10.1016/j.brainres.2010.06.053.

[77] Bolognini N, Olgiati E, Rossetti A, Maravita A. Enhancing multisensory spatial orienting by brain polarization of the parietal cortex. Eur J Neurosci 2010b;31(10):1800-6. https://doi.org/10.1111/j.1460-9568.2010.07211.x.

[78] Sparing R, Thimm M, Hesse MD, Küst J, Karbe H, Fink GR. Bidirectional alterations of interhemispheric parietal balance by non-invasive cortical stimulation. Brain 2009;132(11):3011-20. https://doi.org/10.1093/brain/awp154.

[79] Elmer S, Burkard M, Renz B, Meyer M, Jancke L. Direct current induced short-term modulation of the left dorsolateral prefrontal cortex while learning auditory presented nouns. Behav Brain Funct 2009;5(1):29-. https://doi.org/10.1186/1744-9081-5-29.

[80] Motohashi N, Yamaguchi M, Fujii T, Kitahara Y. Mood and cognitive function following repeated transcranial direct current stimulation in healthy volunteers: A preliminary report. Neurosci Res 2013;77(1-2):64-9. https://doi.org/10.1016/j.neures.2013.06.001.

[81] Sdoia S, Conversi D, Pecchinenda A, Ferlazzo F. Access to consciousness of briefly presented visual events is modulated by transcranial direct current stimulation of left dorsolateral prefrontal cortex. Sci Rep 2019;9(1):10950-9. https://doi.org/10.1038/s41598-019-47527-4.

[82] Brückner S, Kammer T. Both anodal and cathodal transcranial direct current stimulation improves semantic processing. Neurosci 2017;343:269-75. https://doi.org/10.1016/j.neuroscience.2016.12.015.

[83] Pisoni A, Papagno C, Cattaneo Z. Neural correlates of the semantic interference effect: New evidence from transcranial direct current stimulation. Neurosci 2012;223:56-67. https://doi.org/10.1016/j.neuroscience.2012.07.046.

[84] Ihara AS, Mimura T, Soshi T, Yorifuji S, Hirata M, Goto T, et al. Facilitated lexical ambiguity processing by transcranial direct current stimulation over the left inferior frontal cortex. J Cogn Neurosci 2015;27(1):26-34. https://doi.org/10.1162/jocn_a_00703.

[85] Price AR, Peelle JE, Bonner MF, Grossman M, Hamilton RH. Causal evidence for a mechanism of semantic integration in the angular gyrus as revealed by high-definition transcranial direct current stimulation. J Neurosci 2016;36(13):3829-38. https://doi.org/10.1523/JNEUROSCI.3120-15.2016.

[86] Giustolisi B, Vergallito A, Cecchetto C, Varoli E, Romero Lauro LJ. Anodal transcranial direct current stimulation over left inferior frontal gyrus enhances sentence comprehension. Brain Lang 2018;176:36-41. https://doi.org/10.1016/j.bandl.2017.11.001.

[87] Gan T, Huang Y, Hao X, Hu L, Zheng Y, Yang Z. Anodal tDCS over the left frontal eye field improves sustained visual search performance. Perception 2022;51(4):263-75. https://doi.org/10.1177/03010066221086446.

[88] Paladini RE, Wieland FAM, Naert L, Bonato M, Mosimann UP, Nef T, et al. The impact of cognitive load on the spatial deployment of visual attention: Testing the role of interhemispheric balance with biparietal transcranial direct current stimulation. Front Neurosc 2020;13:1391-. https://doi.org/10.3389/fnins.2019.01391.

[89] Orth M, Wagnon C, Neumann-Dunayevska E, Kaller CP, Kloeppel S, Meier B, et al. The left prefrontal cortex determines relevance at encoding and governs episodic memory formation. Cereb Cortex 2023;33(3):612-21. https://doi.org/10.1093/cercor/bhac088.

[90] Berryhill ME, Wencil EB, Branch Coslett H, Olson IR. A selective working memory impairment after transcranial direct current stimulation to the right parietal lobe. Neurosci Lett 2010;479(3):312-6. https://doi.org/10.1016/j.neulet.2010.05.087.

[91] Chi RP, Fregni F, Snyder AW. Visual memory improved by non-invasive brain stimulation. Brain Res 2010;1353:168-75. https://doi.org/10.1016/j.brainres.2010.07.062.

[92] Fregni F, Boggio PS, Nitsche M, Bermpohl F, Antal A, Feredoes E, et al. Anodal transcranial direct current stimulation of prefrontal cortex enhances working memory. Exp Brain Res 2005;166(1):23-30. https://doi.org/10.1007/s00221-005-2334-6.

[93] Ohn SH, Park C-I, Yoo W-K, Ko M-H, Choi KP, Kim G-M, et al. Time-dependent effect of transcranial direct current stimulation on the enhancement of working memory. Neuroreport 2008;19(1):43-7. https://doi.org/10.1097/WNR.0b013e3282f2adfd.

[94] Mulquiney PG, Hoy KE, Daskalakis ZJ, Fitzgerald PB. Improving working memory: Exploring the effect of transcranial random noise stimulation and transcranial direct current stimulation on the dorsolateral prefrontal cortex. Clin Neurophysiol 2011;122(12):2384-9. https://doi.org/10.1016/j.clinph.2011.05.009.

[95] Teo F, Hoy KE, Daskalakis Z, Fitzgerald PB. Investigating the role of current strength in tdcs modulation of working memory performance in healthy controls. Front Psychiatry 2011;2:45-. https://doi.org/10.3389/fpsyt.2011.00045.

[96] Mylius V, Jung M, Menzler K, Haag A, Khader PH, Oertel WH, et al. Effects of transcranial direct current stimulation on pain perception and working memory. Eur J Pain 2012;16(7):974-82. https://doi.org/10.1002/j.1532-2149.2011.00105.x.

[97] Hoy KE, Emonson MRL, Arnold SL, Thomson RH, Daskalakis ZJ, Fitzgerald PB. Testing the limits: Investigating the effect of tDCS dose on working memory enhancement in healthy controls. Neuropsychologia 2013;51(9):1777-84. https://doi.org/10.1016/j.neuropsychologia.2013.05.018.

[98] Meiron O, Lavidor M. Unilateral prefrontal direct current stimulation effects are modulated by working memory load and gender. Brain Stimul 2013;6(3):440-7. https://doi.org/10.1016/j.brs.2012.05.014.

[99] Nozari N, Thompson-Schill SL. More attention when speaking: Does it help or does it hurt? Neuropsychologia 2013;51(13):2770-80. https://doi.org/10.1016/j.neuropsychologia.2013.08.019.

[100] Tanoue RT, Jones KT, Peterson DJ, Berryhill ME. Differential frontal involvement in shifts of internal and perceptual attention. Brain Stimul 2013;6(4):675-82. https://doi.org/10.1016/j.brs.2012.11.003.

[101] Keeser D, Padberg F, Reisinger E, Pogarell O, Kirsch V, Palm U, et al. Prefrontal direct current stimulation modulates resting EEG and event-related potentials in healthy subjects: A standardized low resolution tomography (sLORETA) study. NeuroImage 2011;55(2):644-57. https://doi.org/10.1016/j.neuroimage.2010.12.004.

[102] Gill J, Shah-Basak PP, Hamilton R. It's the thought that counts: Examining the task-dependent effects of transcranial direct current stimulation on executive function. Brain Stimul 2015;8(2):253-9. https://doi.org/10.1016/j.brs.2014.10.018.

[103] Wu Y-J, Tseng P, Chang C-F, Pai M-C, Hsu K-S, Lin C-C, et al. Modulating the interference effect on spatial working memory by applying transcranial direct current stimulation over the right dorsolateral prefrontal cortex. Brain Cogn 2014;91:87-94. https://doi.org/10.1016/j.bandc.2014.09.002.

[104] Friehs MA, Frings C. Offline beats online: Transcranial direct current stimulation timing influences on working memory. Neuroreport 2019a;30(12):795-9. https://doi.org/10.1097/WNR.0000000000001272.

[105] Lukasik KM, Lehtonen M, Salmi J, Meinzer M, Joutsa J, Laine M. No effects of stimulating the left ventrolateral prefrontal cortex with tDCS on verbal working memory updating. Front Neurosc 2018;11:738-. https://doi.org/10.3389/fnins.2017.00738.

[106] Mashal N, Metzuyanim-Gorelick S. New information on the effects of transcranial direct current stimulation on n-back task performance. Exp Brain Res 2019;237(5):1315-24. https://doi.org/10.1007/s00221-019-05500-7.

[107] Naka M, Matsuzawa D, Ishii D, Hamada H, Uchida T, Sugita K, et al. Differential effects of high-definition transcranial direct current stimulation on verbal working memory performance according to sensory modality. Neurosci Lett 2018;687:131-6. https://doi.org/10.1016/j.neulet.2018.09.047.

[108] Nikolin S, Boonstra TW, Loo CK, Martin D. Combined effect of prefrontal transcranial direct current stimulation and a working memory task on heart rate variability. PLOS One 2017;12(8):e0181833-e. https://doi.org/10.1371/journal.pone.0181833.

[109] Röhner F, Breitling C, Rufener KS, Heinze HJ, Hinrichs H, Krauel K, et al. Modulation of working memory using transcranial electrical stimulation: A direct comparison between TACS and TDCS. Front Neurosc 2018;12:761-. https://doi.org/10.3389/fnins.2018.00761.

[110] Talsma LJ, Kroese HA, Slagter HA. Boosting cognition: Effects of multiple-session transcranial direct current stimulation on working memory. J Cogn Neurosci 2017;29(4):755-68. https://doi.org/10.1162/jocn_a_01077.

[111] Trumbo MC, Matzen LE, Coffman BA, Hunter MA, Jones AP, Robinson CSH, et al. Enhanced working memory performance via transcranial direct current stimulation: The possibility of near and far transfer. Neuropsychologia 2016;93(Pt A):85-96. https://doi.org/10.1016/j.neuropsychologia.2016.10.011.

[112] Wang J, Tian J, Hao R, Tian L, Liu Q. Transcranial direct current stimulation over the right DLPFC selectively modulates subprocesses in working memory. PeerJ 2018;2018(5):e4906-e. https://doi.org/10.7717/peerj.4906.

[113] Keshvari F, Hamid-Reza P, Ekhtiari H. The polarity-dependent effects of the bilateral brain stimulation on working memory. Basic Clin Neurosci 2013;4(3):224-31.

[114] Lally N, Nord CL, Walsh V, Roiser JP. Does excitatory fronto-extracerebral tDCS lead to improved working memory performance? F1000Research 2013;2:219-. https://doi.org/10.12688/f1000research.2-219.v2.

[115] Giglia G, Brighina F, Rizzo S, Puma A, Indovino S, Maccora S, et al. Anodal transcranial direct current stimulation of the right dorsolateral prefrontal cortex enhances memory-guided responses in a visuospatial working memory task. Funct Neurol 2014;29(3):189-93. https://doi.org/10.11138/FNeur/2014.29.3.189.

[116] Pupikova M, Simko P, Gajdos M, Rektorova I. Modulation of working memory and resting-state fMRI by tDCS of the right frontoparietal network. Neural Plast 2021;2021:5594305-9. https://doi.org/10.1155/2021/5594305.

[117] Martin DM, Rushby JA, De Blasio FM, Wearne T, Osborne-Crowley K, Francis H, et al. The effect of tDCS electrode montage on attention and working memory. Neuropsychologia 2023;179:108462-. https://doi.org/10.1016/j.neuropsychologia.2022.10846t2.

[118] Nikolin S, Martin D, Loo CK, Boonstra TW. Effects of TDCS dosage on working memory in healthy participants. Brain Stimul 2018;11(3):518-27. https://doi.org/10.1016/j.brs.2018.01.003.

[119] Ramaraju S, Roula MA, McCarthy PW. Transcranial direct current stimulation and working memory: Comparison of effect on learning shapes and English letters. PLOS One 2020;15(7):e0222688-e. https://doi.org/10.1371/journal.pone.0222688.

[120] Zivanovic M, Paunovic D, Konstantinovic U, Vulic K, Bjekic J, Filipovic SR. The effects of offline and online prefrontal vs parietal transcranial direct current stimulation (tDCS) on verbal and spatial working memory. Neurobiol Learn Mem 2021;179:107398-. https://doi.org/10.1016/j.nlm.2021.107398.

[121] Papazova I, Strube W, Wienert A, Henning B, Schwippel T, Fallgatter AJ, et al. Effects of 1 mA and 2 mA transcranial direct current stimulation on working memory performance in healthy participants. Conscious Cogn 2020;83:102959. https://doi.org/doi.org/10.1016/j.concog.2020.102959.

[122] Zhu R, Luo Y, Wang Z, You X. Modality effects in verbal working memory updating: Transcranial direct current stimulation over human inferior frontal gyrus and posterior parietal cortex. Brain Cogn 2020;145:105630-. https://doi.org/10.1016/j.bandc.2020.105630.

[123] Dumont R, Majerus S, Hansenne M. Transcranial direct current stimulation (tDCS) over the intraparietal sulcus does not influence working memory performance. Psychol Belg 2021;61(1):200-11. https://doi.org/10.5334/PB.534.

[124] Zhu R, Luo Y, Wang Z, You X. Within-session repeated transcranial direct current stimulation of the posterior parietal cortex enhances spatial working memory. Cogn Neurosc 2022;13(1):26-37. https://doi.org/10.1080/17588928.2021.1877648.

[125] Ambrus GG, Zimmer M, Kincses ZT, Harza I, Kovács G, Paulus W, et al. The enhancement of cortical excitability over the DLPFC before and during training impairs categorization in the prototype distortion task. Neuropsychologia 2011;49(7):1974-80. https://doi.org/10.1016/j.neuropsychologia.2011.03.026.

[126] Hammer A, Mohammadi B, Schmicker M, Saliger S, Münte TF. Errorless and errorful learning modulated by transcranial direct current stimulation. BMC Neurosci 2011;12(1):72-. https://doi.org/10.1186/1471-2202-12-72.

[127] Javadi AH, Walsh V. Transcranial direct current stimulation (tDCS) of the left dorsolateral prefrontal cortex modulates declarative memory. Brain Stimul 2012;5(3):231-41. https://doi.org/10.1016/j.brs.2011.06.007.

[128] Asthana M, Nueckel K, Mühlberger A, Neueder D, Polak T, Domschke K, et al. Effects of transcranial direct current stimulation on consolidation of fear memory. Front Psychiatry 2013;4:107-. https://doi.org/10.3389/fpsyt.2013.00107.

[129] Javadi AH, Cheng P. Transcranial direct current stimulation (tDCS) enhances reconsolidation of long-term memory. Brain Stimul 2013;6(4):668-74. https://doi.org/10.1016/j.brs.2012.10.007.

[130] Kongthong N, Minami T, Nakauchi S. Semantic processing in subliminal face stimuli: An EEG and tDCS study. Neurosci Lett 2013;544:141-6. https://doi.org/10.1016/j.neulet.2013.04.002.

[131] Santiesteban I, Banissy Michael J, Catmur C, Bird G. Enhancing social ability by stimulating right temporoparietal junction. Curr Biol 2012;22(23):2274-7. https://doi.org/10.1016/j.cub.2012.10.018.

[132] Beharelle AR, Polanía R, Hare TA, Ruff CC. Transcranial stimulation over frontopolar cortex elucidates the choice attributes and neural mechanisms used to resolve exploration–exploitation trade-offs. J Neurosci 2015;35(43):14544-56. https://doi.org/10.1523/JNEUROSCI.2322-15.2015.

[133] Martin AK, Dzafic I, Ramdave S, Meinzer M. Causal evidence for task-specific involvement of the dorsomedial prefrontal cortex in human social cognition. Soc Cogn Affect Neurosci 2017;12(8):1209-18. https://doi.org/10.1093/scan/nsx063.

[134] Habich A, Slotboom J, Peter J, Wiest R, Klöppel S. No effect of anodal tDCS on verbal episodic memory performance and neurotransmitter levels in young and elderly participants. Neural Plast 2020;2020:8896791-15. https://doi.org/10.1155/2020/8896791.

[135] Huang Y, Mohan A, McLeod SL, Luckey AM, Hart J, Vanneste S. Polarity-specific high-definition transcranial direct current stimulation of the anterior and posterior default mode network improves remote memory retrieval. Brain Stimul 2021;14(4):1005-14. https://doi.org/10.1016/j.brs.2021.06.007.

[136] Flöel A, Rösser N, Michka O, Knecht S, Breitenstein C. Noninvasive brain stimulation improves language learning. J Cogn Neurosci 2008;20(8):1415-22. https://doi.org/10.1162/jocn.2008.20098.

[137] Ross LA, McCoy D, Wolk DA, Coslett HB, Olson IR. Improved proper name recall by electrical stimulation of the anterior temporal lobes. Neuropsychologia 2010;48(12):3671-4. https://doi.org/10.1016/j.neuropsychologia.2010.07.024.

[138] Sparing R, Dafotakis M, Meister IG, Thirugnanasambandam N, Fink GR. Enhancing language performance with non-invasive brain stimulation—A transcranial direct current stimulation study in healthy humans. Neuropsychologia 2008;46(1):261-8. https://doi.org/10.1016/j.neuropsychologia.2007.07.009.

[139] de Vries MH, Barth ACR, Maiworm S, Knecht S, Zwitserlood P, Flöel A. Electrical stimulation of Broca's area enhances implicit learning of an artificial grammar. J Cogn Neurosci 2010;22(11):2427-36. https://doi.org/10.1162/jocn.2009.21385.

[140] Fertonani A, Rosini S, Cotelli M, Rossini PM, Miniussi C. Naming facilitation induced by transcranial direct current stimulation. Behav Brain Res 2010;208(2):311-8. https://doi.org/10.1016/j.bbr.2009.10.030.

[141] Choi JY, Perrachione TK. Noninvasive neurostimulation of left temporal lobe disrupts rapid talker adaptation in speech processing. Brain Lang 2019;196:104655-. https://doi.org/10.1016/j.bandl.2019.104655.

[142] Henseler I, Mädebach A, Kotz SA, Jescheniak JD. Modulating brain mechanisms resolving lexico-semantic interference during word production: A transcranial direct current stimulation study. J Cogn Neurosci 2014;26(7):1403-17. https://doi.org/10.1162/jocn_a_00572.

[143] Meinzer M, Yetim Ö, McMahon K, de Zubicaray G. Brain mechanisms of semantic interference in spoken word production: An anodal transcranial direct current stimulation (atDCS) study. Brain Lang 2016;157-158:72-80. https://doi.org/10.1016/j.bandl.2016.04.003.

[144] Westwood SJ, Olson A, Miall RC, Nappo R, Romani C. Limits to tDCS effects in language: Failures to modulate word production in healthy participants with frontal or temporal tDCS. Cortex 2017;86:64-82. https://doi.org/10.1016/j.cortex.2016.10.016.

[145] Wirth M, Rahman RA, Kuenecke J, Koenig T, Horn H, Sommer W, et al. Effects of transcranial direct current stimulation (tDCS) on behaviour and electrophysiology of language production. Neuropsychologia 2011;49(14):3989-98. https://doi.org/10.1016/j.neuropsychologia.2011.10.015.

[146] Younger JW, Wagner MR, Booth JR. Weighing the cost and benefit of transcranial direct current stimulation on different reading subskills. Front Neurosc 2016;10:262-. https://doi.org/10.3389/fnins.2016.00262.

[147] Peretz Y, Lavidor M. Enhancing lexical ambiguity resolution by brain polarization of the right posterior superior temporal sulcus. Cortex 2013;49(4):1056-62. https://doi.org/10.1016/j.cortex.2012.03.015.

[148] Weltman K, Lavidor M. Modulating lexical and semantic processing by transcranial direct current stimulation. Exp Brain Res 2013;226(1):121-35. https://doi.org/10.1007/s00221-013-3416-5.

[149] Malyutina S, Zelenkova V, Buivolova O, Oosterhuis EJ, Zmanovsky N, Feurra M. Modulating the interhemispheric balance in healthy participants with transcranial direct current stimulation: No significant effects on word or sentence processing. Brain Lang 2018;186:60-6. https://doi.org/10.1016/j.bandl.2018.09.004.

[150] Kurmakaeva D, Blagovechtchenski E, Gnedykh D, Mkrtychian N, Kostromina S, Shtyrov Y. Acquisition of concrete and abstract words is modulated by tDCS of Wernicke’s area. Sci Rep 2021;11(1):1508-. https://doi.org/10.1038/s41598-020-79967-8.

[151] Fiori V, Kunz L, Kuhnke P, Marangolo P, Hartwigsen G. Transcranial direct current stimulation (tDCS) facilitates verb learning by altering effective connectivity in the healthy brain. NeuroImage 2018;181:550-9. https://doi.org/10.1016/j.neuroimage.2018.07.040.

[152] Longo F, Braun M, Hutzler F, Richlan F. Impaired semantic categorization during transcranial direct current stimulation of the left and right inferior parietal lobule. J Neurolinguistics 2022;62:101058. https://doi.org/10.1016/j.jneuroling.2022.101058.

[153] Bartel G, Marko M, Rameses I, Lamm C, Riečanský I. Left prefrontal cortex supports the recognition of meaningful patterns in ambiguous stimuli. Front Neurosc 2020;14:152-. https://doi.org/10.3389/fnins.2020.00152.

[154] Brunyé TT, Moran JM, Cantelon J, Holmes A, Eddy MD, Mahoney CR, et al. Increasing breadth of semantic associations with left frontopolar direct current brain stimulation: A role for individual differences. Neuroreport 2015;26(5):296-301. https://doi.org/10.1097/WNR.0000000000000348.

[155] Chrysikou EG, Morrow HM, Flohrschutz A, Denney L. Augmenting ideational fluency in a creativity task across multiple transcranial direct current stimulation montages. Sci Rep 2021;11(1):8874-. https://doi.org/10.1038/s41598-021-85804-3.

[156] Ghanavati E, Nejati V, Salehinejad MA. Transcranial direct current stimulation over the posterior parietal cortex (PPC) enhances figural fluency: Implications for creative cognition. J Cogn Enhanc 2018;2(1):88-96. https://doi.org/10.1007/s41465-017-0059-7.

[157] Green AE, Spiegel KA, Giangrande EJ, Weinberger AB, Gallagher NM, Turkeltaub PE. Thinking cap plus thinking zap: tDCS of frontopolar cortex improves creative analogical reasoning and facilitates conscious augmentation of state creativity in verb generation. Cereb Cortex 2017;27(4):2628-39. https://doi.org/10.1093/cercor/bhw080.

[158] Hertenstein E, Waibel E, Frase L, Riemann D, Feige B, Nitsche MA, et al. Modulation of creativity by transcranial direct current stimulation. Brain Stimul 2019;12(5):1213-21. https://doi.org/10.1016/j.brs.2019.06.004.

[159] Koizumi K, Ueda K, Li Z, Nakao M. Effects of transcranial direct current stimulation on brain networks related to creative thinking. Front Hum Neurosci 2020;14:541052-. https://doi.org/10.3389/fnhum.2020.541052.

[160] Li Y, Beaty RE, Luchini S, Dai DY, Xiang S, Qi S, et al. Accelerating creativity: Effects of transcranial direct current stimulation on the temporal dynamics of divergent thinking. Creat Res J 2023;35(2):169-88. https://doi.org/10.1080/10400419.2022.2068297.

[161] Mayseless N, Shamay-Tsoory SG. Enhancing verbal creativity: Modulating creativity by altering the balance between right and left inferior frontal gyrus with tDCS. Neurosci 2015;291:167-76. https://doi.org/10.1016/j.neuroscience.2015.01.061.

[162] Pick H, Lavidor M. Modulation of automatic and creative features of the Remote Associates Test by angular gyrus stimulation. Neuropsychologia 2019;129:348-56. https://doi.org/10.1016/j.neuropsychologia.2019.04.010.

[163] Ruggiero F, Lavazza A, Vergari M, Priori A, Ferrucci R. Transcranial direct current stimulation of the left temporal lobe modulates insight. Creat Res J 2018;30(2):143-51. https://doi.org/10.1080/10400419.2018.1446817.

[164] Salvi C, Beeman M, Bikson M, McKinley R, Grafman J. TDCS to the right anterior temporal lobe facilitates insight problem-solving. Sci Rep 2020;10(1):946-. https://doi.org/10.1038/s41598-020-57724-1.

[165] Xiang S, Qi S, Li Y, Wang L, Dai DY, Hu W. Trait anxiety moderates the effects of tDCS over the dorsolateral prefrontal cortex (DLPFC) on creativity. Pers Individ Differ 2021;177:110804. https://doi.org/10.1016/j.paid.2021.110804.

[166] Aihara T, Ogawa T, Shimokawa T, Yamashita O. Anodal transcranial direct current stimulation of the right anterior temporal lobe did not significantly affect verbal insight. PLoS One. 2017 Sep 13;12(9):e0184749.

[167] Luft CD, Zioga I, Banissy MJ, Bhattacharya J. Relaxing learned constraints through cathodal tDCS on the left dorsolateral prefrontal cortex. Scientific reports. 2017 Jun 7;7(1):2916.

[168] Chi RP, Snyder AW. Brain stimulation enables the solution of an inherently difficult problem. Neuroscience letters. 2012 May 2;515(2):121-4.

[169] Guo J, Luo J, An Y, Xia T. tDCS Anodal Stimulation of the Right Dorsolateral Prefrontal Cortex Improves Creative Performance in Real-World Problem Solving. Brain Sciences. 2023 Mar 6;13(3):449.

[170] Lundie M, Dasara H, Beeghly C, Kazmi A, Krawczyk D. High‐Definition Transcranial Direct Current Stimulation Over the Left Frontopolar Cortex Promotes Analogical Reasoning. Mind, Brain, and Education. 2022 Aug;16(3):209-20.

[171] Wang Y, Guo X, Wang M, Kan Y, Zhang H, Zhao H, Meilin W, Duan H. Transcranial direct current stimulation of bilateral dorsolateral prefrontal cortex eliminates creativity impairment induced by acute stress. International Journal of Psychophysiology. 2022 Jan 1;171:1-1.

[172] Wang Y, Zhang J, Li Y, Qi S, Zhang F, Ball LJ, Duan H. Preventing prefrontal dysfunction by tDCS modulates stress-induced creativity impairment in women: an fNIRS study. Cerebral Cortex. 2023 Oct 15;33(20):10528-45.
